# Unheralded high MHC Class II polymorphism in the abundant Atlantic herring resolved by long-read sequencing

**DOI:** 10.1101/2025.06.08.658498

**Authors:** Minal Jamsandekar, Fahime M Sangdehi, Florian Berg, Michael F. Criscitiello, Brian W Davis, Mårten Larsson, Jingyi Li, Mats E Pettersson, Leif Andersson

**Affiliations:** Department of Veterinary Integrative Biosciences, Texas A&M University, College Station, USA; Department of Medical Biochemistry and Microbiology, Uppsala University, Uppsala, Sweden; Institute of Marine Research, Norway; Department of Veterinary Pathobiology, Texas A&M University, College Station, USA; Department of Pharmaceutical Biosciences, Uppsala University, Uppsala, Sweden

## Abstract

Major Histocompatibility Complex (MHC) genes are the most polymorphic in vertebrate genomes due to their important function of presenting a diversity of peptides from pathogens to initiate an adaptive immune response. Here, we have characterized MHC class II genes and explored their polymorphism and evolution in one of the most abundant vertebrates in the world, Atlantic herring (*Clupea harengus*). The vast population size and schooling behavior make Atlantic herring an attractive target for pathogens. Hence, we hypothesized that it would maintain exceptionally high MHC polymorphism. We used PacBio HiFi long-read whole genome sequencing data of 14 individuals from three different geographic regions in the Atlantic Ocean and Baltic Sea. The analysis identified nine MHC class II loci distributed across four chromosomes. We found two distinct lineages of class II genes, a highly polymorphic one denoted *DA* and a non-polymorphic *DB* lineage, arranged as alpha-beta gene pairs (*DAA*-*DAB* or *DBA-DBB*). The *DA* genes showed extremely high nucleotide diversity in exon 2, strong signatures of positive selection (*dN/dS* >> 1) as well as copy number variation. Structure prediction revealed that all highly polymorphic amino acid residues occurred in the predicted peptide binding cleft. Two of the most polymorphic loci showed distinct allelic groupings (supertypes), with high sequence similarity within supertypes and high sequence divergence between supertypes. Most of the haplotypes had genes from different supertypes, thus maintaining diverse class II repertoire in a single individual. There was also a highly significant nonrandom association of *DAA* and *DAB* alleles within supertypes, strongly suggesting coevolution of *DAA* and *DAB* alleles forming the peptide binding domain. This study reveals that the herring MHC class II genes are among the most, if not the most, polymorphic so far described in vertebrates. Their exceptional polymorphism surpasses that of human due to a larger gene repertoire, copy number variation, and pronounced sequence divergence among alleles. This unheralded polymorphism is most likely explained by the combined effects of the vast population size, minimizing genetic drift, and strong pathogen-driven balancing selection.

## Introduction

The major histocompatibility complex (MHC) is present in all jawed vertebrates^1^. MHC genes are key components of the vertebrate immune system and encode MHC class I and class II molecules that present peptides to other cells in the immune system. MHC class I and class II genes show extremely high levels of genetic polymorphism in most vertebrates^2^. Characteristic features of MHC polymorphisms are (1) large number of alleles at multiple loci^3^, (2) alleles differ by many nonsynonymous substitutions, and (3) deep allele divergences that may predate speciation events, so called trans-species polymorphisms^2^. This extreme diversity allows the immune system to recognize peptides from a wide range of pathogens and thereby enhance disease resistance^4–7^. There is evidence that this extensive polymorphism is maintained by balancing selection mechanisms which includes heterozygote advantage, negative frequency-dependent selection, and fluctuating selection^8^. The consensus view is that pathogen-mediated selection is the major mechanism for the maintenance of MHC polymorphisms as allelic variants differ in their capacity to present peptides to T lymphocytes of the immune system^1,2,8^. However, it has also been proposed that MHC-based mating preferences may contribute to enhance MHC diversity due to sexual selection^9^.

The present study characterizes MHC class II genes in Atlantic herring (*Clupea harengus*) at locus and haplotype level, and tests the prediction that this species will show high MHC diversity, if pathogen-mediated selection is the major mechanism causing allele diversity. Atlantic herring occur in the North Atlantic Ocean and the Baltic Sea. It is one of the most abundant vertebrates on earth with a census population size of about 10^12^ individuals and an estimated effective population size of 4.4 x 10^5^ (Ref. ^10^); a single school may include a billion individuals. There are two major reasons for expecting unusually high levels of MHC polymorphism in herring. Firstly, their large effective population size and high fecundity leave large room for natural selection to operate. Loss of genetic diversity due to drift is minute^11^. Secondly, the large schools of Atlantic herring, and their abundance, make them an attractive target for pathogens. However, if MHC-based mating preferences are a major driver for high MHC polymorphism, it is expected that Atlantic herring will show low levels of MHC polymorphisms. This is because it is a broadcast spawner with no mate choice; instead, spawning aggregations occur where sperm and eggs are randomly mixed.

Here, we have used PacBio long read sequencing of 14 individuals to robustly characterize the genetic diversity at MHC class II loci of Atlantic herring. We report that this species has one of the most, if not the most, polymorphic MHC class II allele sets so far described. Results strongly support the notion that pathogen-mediated selection is the primary driver for extreme polymorphism of MHC genes in vertebrates.

## Results

### Genome organization and copy number variation of MHC II genes

PacBio long read sequencing was carried out using six Atlantic herring sampled in the Celtic Sea, two Atlantic herring from the Norwegian Sea, and six Baltic herring, considered a distinct subspecies, from the Baltic Sea. Haplotype-phased genome assemblies were constructed using hifiasm for each indiviual^12^ resulting in 28 high quality PacBio haplotype assemblies from 14 individuals, all of which, together with the reference assembly, were used in the downstream analyses (Extended Data Table 1). Manual annotation of MHC class II genes in the current herring reference assembly^13^ resulted in the detection of 9 alpha (including one pseudogene) and 10 beta genes at six different loci, designated 1 to 6, distributed across three chromosomes (Chr5, 7, and 13). Each of these loci contained one or more closely linked alpha-beta gene pairs. Applying this to the 28 new haplotype assemblies (Extended Data Table 1) showed that almost all had a larger number of MHC class II genes than present in the reference assembly, suggesting that some MHC regions have been collapsed during the construction of the reference assembly but are more resolved in the new haplotype assemblies. In addition to chromosomes 5, 7, and 13, MHC class II genes were also present on chromosome 8 in some new assemblies. Thus, the total number of MHC class II loci increased from 6 in the reference assembly to 9, of which Locus 1 to 6 were present in all assemblies while Locus 7 to 9 were present in only a few (Fig. 1a; Supplementary Fig. 1).

**Fig. 1.**
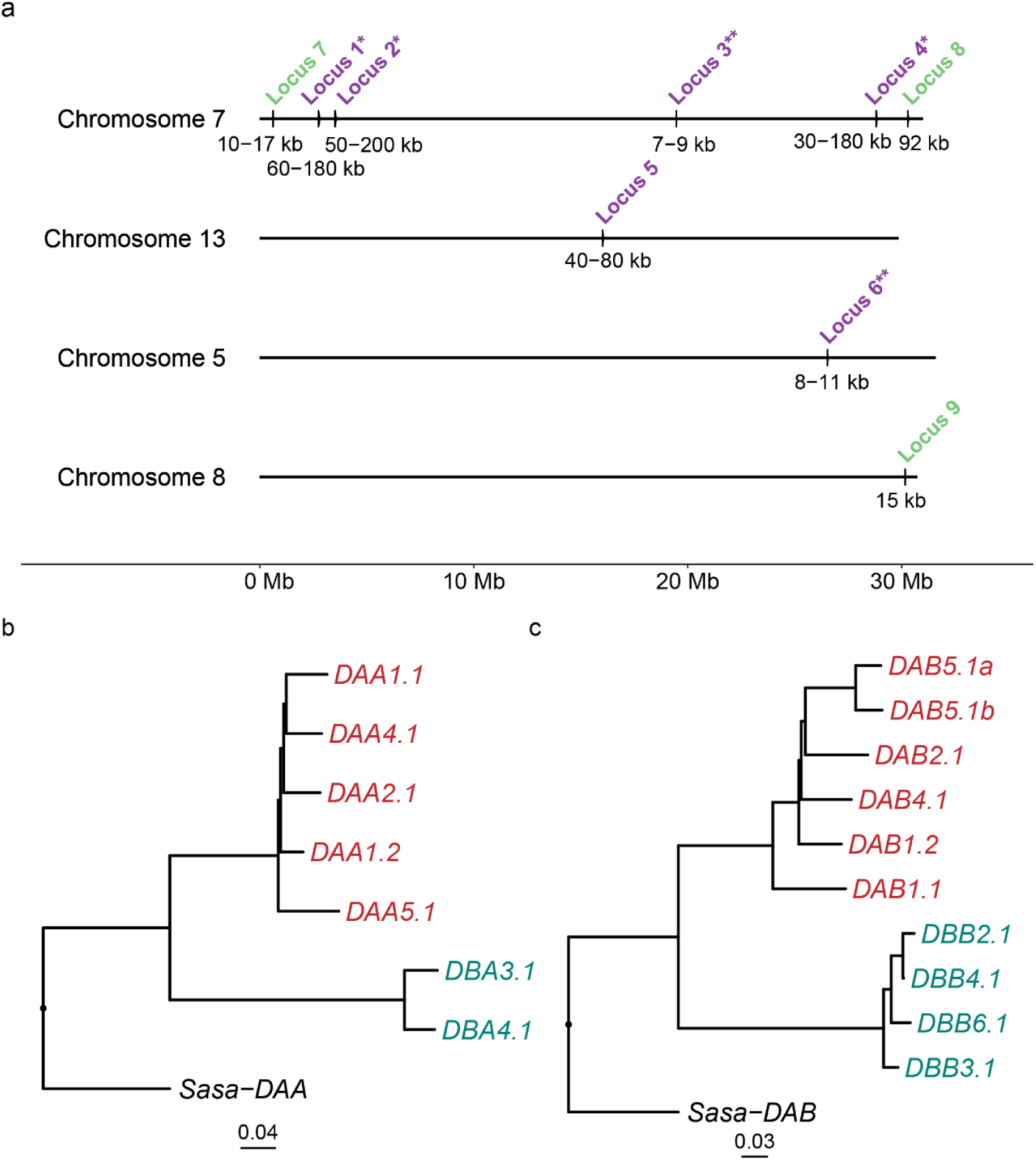
Overview of MHC class II genes in Atlantic herring. **a** Genomic organization. Each locus (#1-9) contains one or more closely linked alpha-beta gene pairs. Loci present in all haplotypes are labeled in purple while loci present in only some haplotypes are labeled in green. Numbers below the locus represent its genomic size. *Loci with genes from both DA and DB lineages. **Loci with genes exclusively from DB lineage. The genomic distances between loci were estimated as follows: Locus 7 and 1: 2.1 Mb; Locus 1 and 2: 0.7 Mb; Locus 2 and 3: 15.9 Mb; Locus 3 and 4: 9.3 Mb; Locus 4 and 8: 1.4 Mb. **b, c** Genetic distance trees constructed using Neighbor-Joining method based on coding sequences from the reference assembly for alpha (**b)** and beta **(c)** chain genes. *Sasa-DAA*\*0101 and *Sasa-DAB*\*0101 from IPD-MHC database (https://www.ebi.ac.uk/ipd/mhc/) from Atlantic salmon (*Salmo salar*) are used as outgroup for alpha and beta trees, respectively. The numbers at the bottom represent the branch length indicative of the genetic divergence between genes.

**Table 1.** Number of MHC class II genes at each locus among haplotypes based on 28 PacBio based haplotype assemblies from 14 individual Atlantic and Baltic herring. The genomic intervals for Locus 1-6 correspond to the reference assembly. As Locus 7-9 are absent in the reference assembly, genomic intervals are from PacBio assemblies of BS4_hap2, CS5_hap1, and BS2_hap1, respectively.

| **Locus** | **Chr: interval (Mb)** | **Number of genes** |  |  |  |
| --- | --- | --- | --- | --- | --- |
|  |  | ***DAA*** | ***DAB*** | ***DBA*** | ***DBB*** |
| 1 | 7: 2.73 - 2.80 | 1-3 | 1-3 | 0 | 0-2 |
| 2 | 7: 3.52 - 3.58 | 1-3 | 1-3 | 1-2 | 1-2 |
| 3 | 7: 19.46 - 19.47 | 0 | 0 | 1 | 1 |
| 4 | 7: 28.81 - 28.84 | 1-2 | 1-2 | 0-2 | 0-2 |
| 5 | 13: 16.02 - 16.07 | 1 | 2 | 0 | 0 |
| 6 | 5: 26.53 - 26.54 | 0 | 0 | 1 | 1 |
| 7 | 7: 0.62 - 0.63 | 0-1 | 0-1 | 0 | 0 |
| 8 | 7: 30.30 - 30.37 | 0-2 | 0-2 | 0 | 0 |
| 9 | 8: 31.93 - 31.94 | 0-1 | 0-1 | 0 | 0 |

Genetic distance trees for MHC class II alpha (Fig. 1b) and beta (Fig. 1c) genes present in the reference assembly showed that both alpha and beta genes formed two distinct clades that we denoted DA and DB, respectively. *DAA* (alpha) and *DAB* (beta) genes as well as *DBA* (alpha) and *DBB* (beta) genes were present as alpha-beta tandem gene pairs (Fig. 2). This formed the basis for our proposed nomenclature, for instance *DAA1.1* is a DA alpha gene belonging to Locus 1 with the last digit (.1) used to handle the situation where multiple copies of alpha or beta genes were present at a given locus. A maximum likelihood phylogeny using full length amino acid sequences of MHC class II genes from other fish and tetrapods did not establish clear orthologous relationship to herring genes (Extended Data Fig. 1), reflecting the deep splits in the teleost phylogeny and/or concerted evolution of class II genes in different lineages of fish. The gene composition at the nine class II loci is summarized in Table 1. An important component of the genetic diversity is the extensive copy number variation, particularly at loci 1, 2, and 4. As an example, the size of Locus 1 varies from about 70 kb to more than 300 kb among haplotypes due to copy number variation of genes and variable length of non-coding sequences (Fig. 2; Supplementary Fig. 2). For this reason, it is difficult to define MHC loci in Atlantic herring as harboring allelic series for single MHC class II genes. We found it more appropriate to define each locus as a cluster of closely linked MHC class II genes at a given genomic location as illustrated in Fig. 2. Hence, we labeled genes according to their order of occurrence on the chromosome. For example, [*DAA1.1*, *DAB1.1*], [*DAA1.2, DAB1.2*], [*DAA1.3, DAB1.3*], and [*DBA1.1, DBB1.1*] denote three DA and one DB gene pairs for alpha and beta chain heterodimers at Locus 1, respectively.

**Fig. 2.**
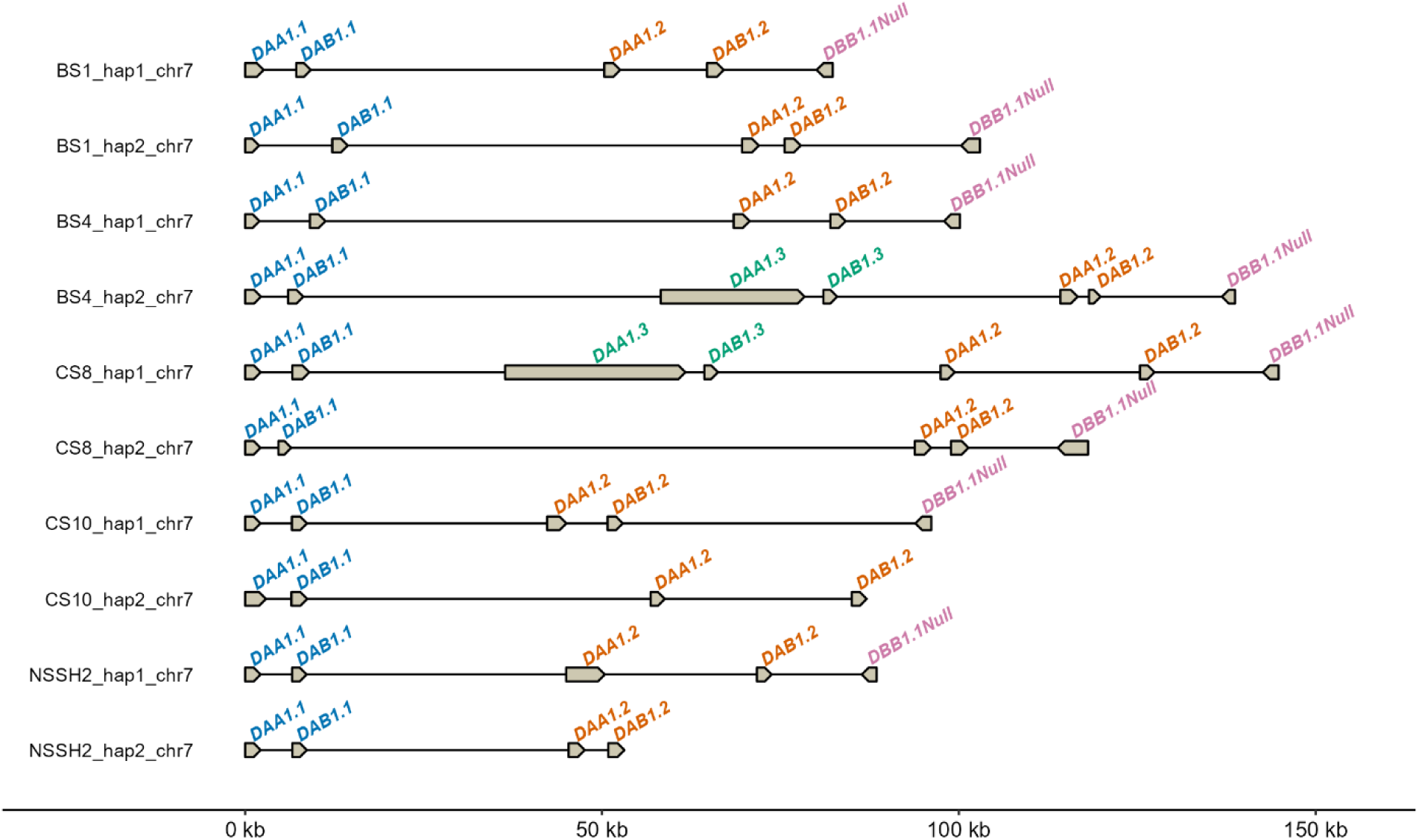
Overview of the polymorphic MHC class II locus 1 on chromosome 7 in Atlantic herring. Gene organization on the two haplotypes (hap1 and hap2) from five individual fish. The designation *DBB1.1Null* indicates that this gene encodes a mutation that disrupts the coding sequence.

A characteristic of MHC class II genes in Atlantic herring is that for highly polymorphic genes it is not possible to assign a gene to a specific locus only based on sequence identities because there is an overlap in intra- and inter-locus sequence identities (Fig. 3; Extended Data Fig. 2). Therefore, the gene designations used throughout this paper are a combination of sequence identity and genomic location. Interestingly, high inter-locus identities only occurred for genes located on the same chromosomes implying that intrachromosomal recombination may have contributed to sequence exchange between genes.

**Fig. 3.**
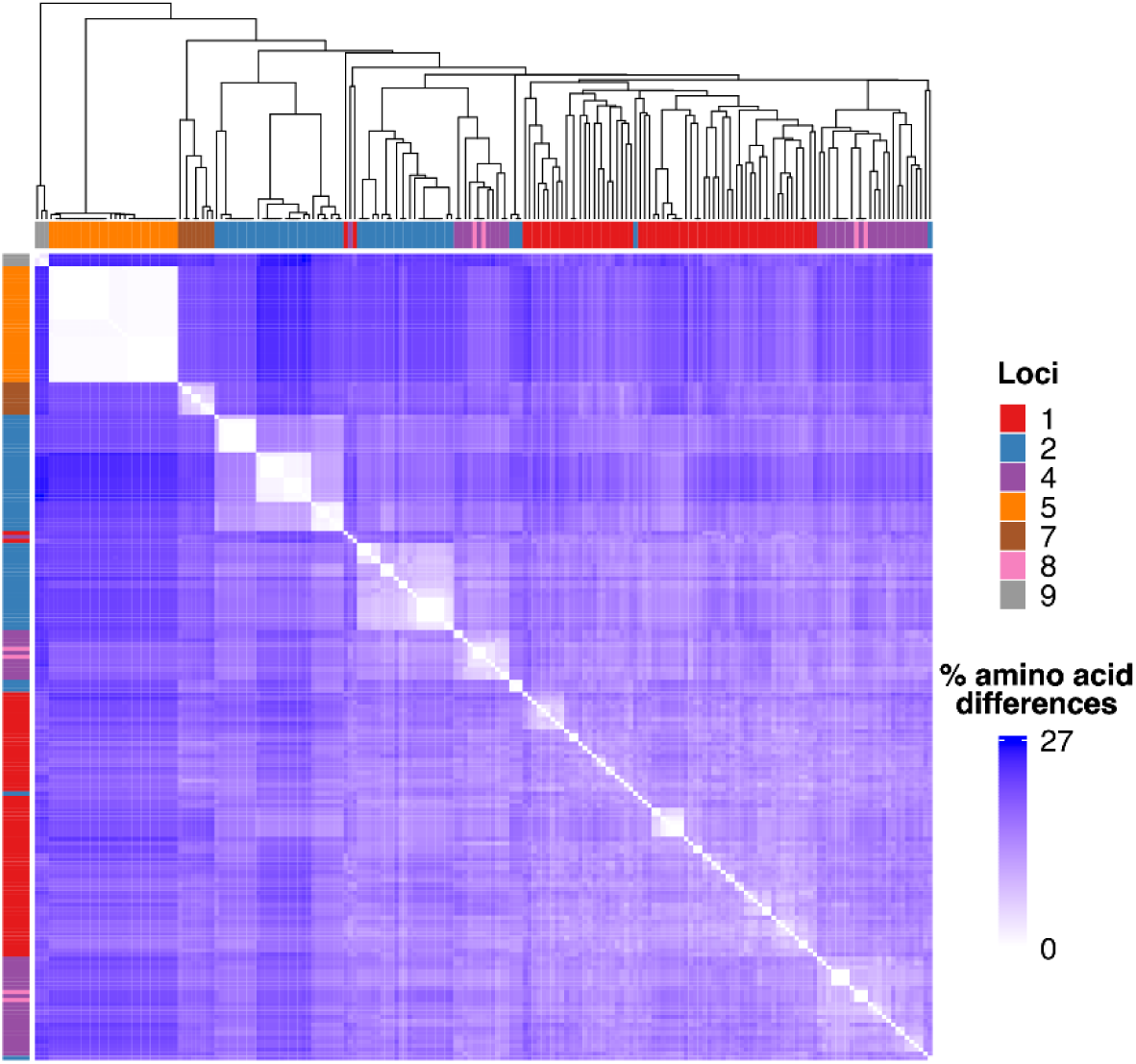
Clustering of amino acid sequences from DA alpha genes. Heatmap and dendrogram showing hierarchical clustering of all DAA amino acid sequences across the genome. Heatmap gradients correspond to pairwise percent amino acid differences. Source loci are color-coded in the annotation bar.

In summary, our analyses using 28 assemblies identified nine MHC class II loci present on four chromosomes, two different clades of class II genes (DA and DB), and that copy number variation is an important component of the observed genetic diversity.

### Extensive sequence polymorphism of MHC class II genes

Analysis of nucleotide diversity (*π*) per exon of herring MHC class II genes revealed a clear pattern (Fig. 4a, b). The majority of DA genes are highly polymorphic whereas none of the DB genes are highly polymorphic, implying that the members of these two lineages have different functions. Possibly, DA functionally corresponds to the classical class II genes, while DB may represent non-classical genes as defined in other species^14^. Remarkably, we identified up to 29 different alleles among 29 assembled haplotypes at the most polymorphic DA loci (Fig. 4a, b). The functional significance of the polymorphism is evident by the fact that exon 2, encoding the peptide binding site in MHC class II molecules^15^, is by far the most polymorphic exon. The genome-wide average *π* in the herring genome is ∼0.3% whereas *π* for exon 2 in the most polymorphic DA genes are in the range 10-20%. However, not all DA genes are highly polymorphic, the *DAA* and *DAB* genes at Locus 5 and 8 showed no or limited polymorphism in our sample of haplotypes (Fig. 4a, b).

**Fig. 4.**
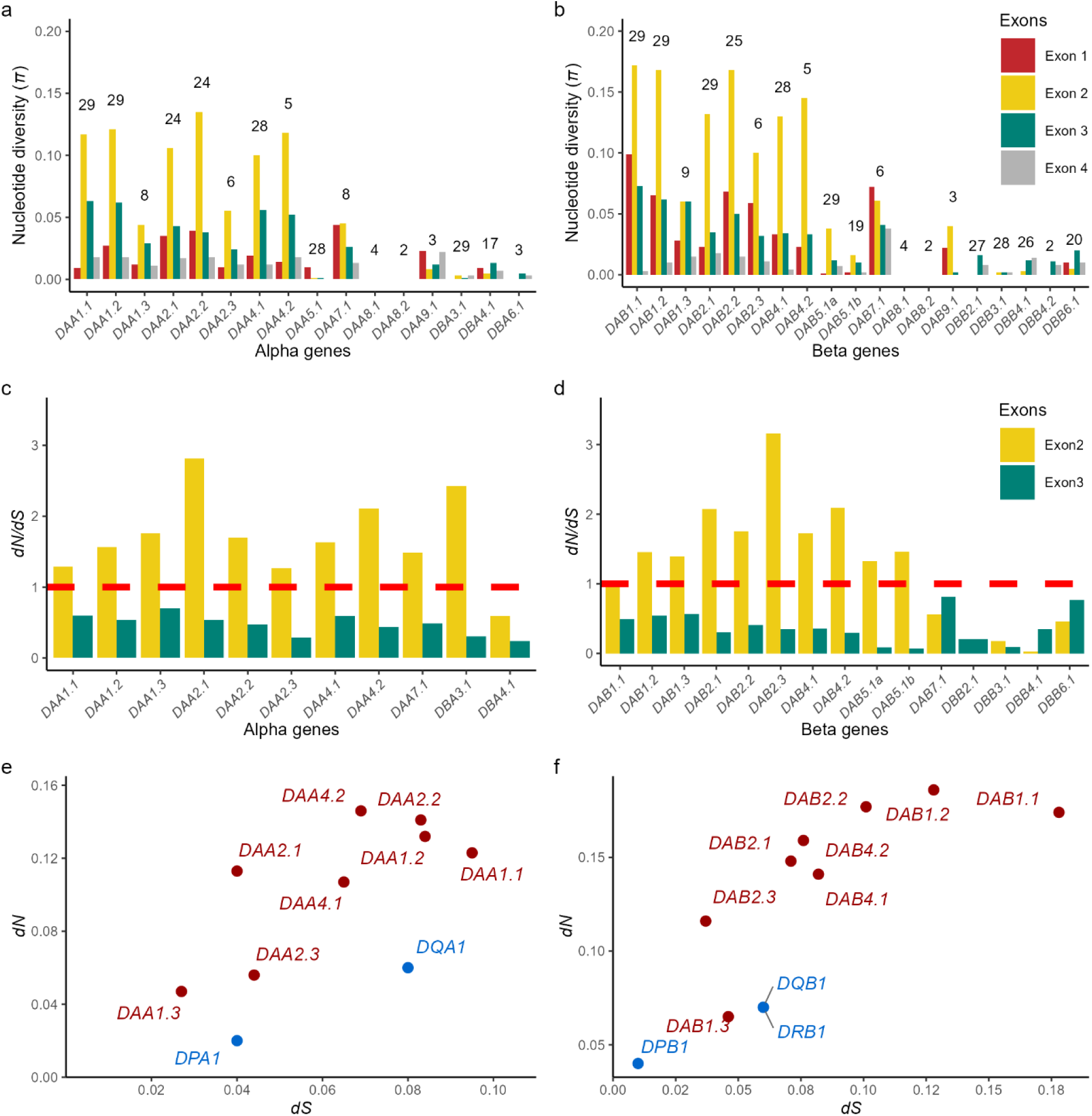
Nucleotide diversity (*π*) and the ratio of non-synonymous substitutions over synonymous substitutions (*dN/dS*) in MHC class II genes in Atlantic herring. a, **b** *π* for all four exons of all alpha (**a**) and beta (**b**) genes. The number of sequences used to calculate *π* are shown above the bars for each gene. c, d *dN/dS* for exon 2 and exon 3 of alpha (c) and beta (**d**) genes. The red dashed line indicates *dN/dS* = 1. **e, f** Scatter plot of exon 2 *dN* and *dS* values of human and herring MHC class II alpha (**e**) and beta (**f**) genes. Human genes are colored in blue while herring genes are colored in red.

We next calculated the rate of non-synonymous (*dN*) and synonymous (*dS*) substitutions for exon 2 and exon 3 sequences of alpha and beta genes (Fig. 4c, d; Supplementary Table 1); a *dN/dS* ratio >1.0 provides evidence for positive or balancing selection whereas a *dN/dS* ratio <1.0 is consistent with purifying selection. This analysis demonstrated that herring *DAA* and *DAB* genes are under strong balancing selection. This is indicated by *dN/dS* ratios higher than 1.0 for exon 2, while purifying selection (*dN/dS* <<1.0) dominates for exon 3 sequences. Non-polymorphic DB genes showed *dN/dS* ratios lower than 1 except for *DBA3.1*, but this was caused by very low *dS* rather than high *dN*. Positively selected sites classified by PAML analysis^16^ are tabulated in Supplementary File 1 and shown in the amino acid alignment in Supplementary Fig. 3.

We noted that *dN/dS* for the most polymorphic genes in herring are much higher than those for the most polymorphic human MHC class II genes (*dN/dS* = ∼1.0)^17^. To understand if the higher *dN/dS* values are due to higher *dN* or lower *dS*, we compared the herring data with corresponding human *dN* and *dS* estimates^17^. The most striking difference is that most herring genes have a much higher *dN* than their human counterparts (Fig. 4e, f; Supplementary Table 1). For class II beta chain genes, there is also on average higher *dS* in herring. Thus, herring alleles tend to have a deeper ancestry (higher *dS*) and natural selection has had a more prominent effect on polymorphism in herring (higher *dN/dS*).

### Sequence diversity is highly enriched at predicted peptide binding residues

To further characterize the DA polymorphism across exon 2 and how it relates to the peptide binding residues (PBRs), we first predicted the PBRs in herring MHC class II genes using known PBRs in human^18^ and chicken^19^ (Fig. 5a, b; Supplementary Fig. 4). We then calculated normalized Shannon entropy values per individual amino acid residues, where higher values indicate a greater degree of polymorphism. Interestingly, Shannon entropy peaks were observed at approximately every three to four residues along the alpha helix, suggesting that these residues may be positioned on the same side of the helix and potentially correspond to the PBRs (Fig. 5). This pattern aligns with the structure of alpha helices, where every n+3.6^th^ residue is oriented toward the same side of the helix^20^, in this case possibly facing peptide-binding pocket.

**Fig. 5.**
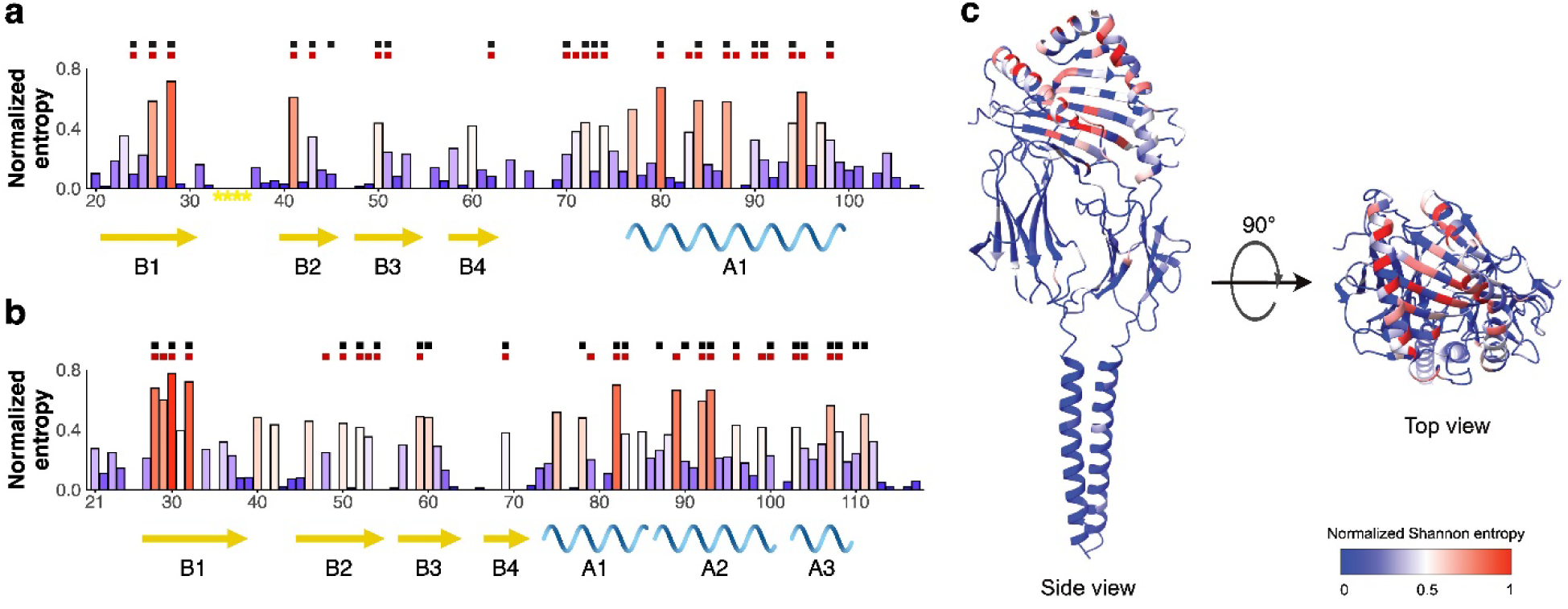
Amino acid and protein structure properties of MHC class II peptide binding residues (PBRs). **a, b** Bar plot of normalized Shannon entropy values for each amino acid site in the α1 domain (**a**) and β1 domain (**b**) of Locus 2 gene. Shannon entropy values are colored with red-white-blue gradient indicating highest to lowest diversity. The scale below the bar plot indicates amino acid positions beginning at the first residue of exon 2 in the full-length protein. The black and red squares above the bar plot represent human^18^ and chicken PBRs^19^, respectively. Arrows and helices below the bar plot correspond to the protein secondary structures beta sheet and alpha helix, respectively. **c** Predicted quaternary structure of an MHC class II protein modeled with AlphaFold3 using protein sequences encoded by *CS4_hap1_DA2.1* as template. The color gradient from red-white-blue corresponds to the normalized Shannon entropy values as in **a** and **b**.

We next built a protein quaternary structure model using Locus 2 gene sequences Extended Data Fig. 3; Supplementary Fig. 5). The normalized Shannon entropy values revealed a remarkably strong association, where the most polymorphic residues are oriented toward the presumed peptide-binding pocket formed by the flanking alpha helices and beta sheet floor of the MHC class II alpha and beta chains (Fig. 5c; Supplementary Video 1).

### Nonrandom combinations of gene copies enhance diversity at a haplotype level

We performed inter- and intra-locus comparisons of the amino acid sequences of MHC class II alleles to understand how the polymorphism is distributed across loci. In the inter-locus comparison of DA lineage sequences, Locus 5, 7, and 9 showed strict locus-specific clustering of alleles, as opposed to Locus 1, 2, 4, and 8, indicating that the latter set of loci shared alleles with each other (Fig. 3; Extended Data Fig. 2). We further made intra-locus comparisons of polymorphic DA lineage sequences and found a distinct pattern for Locus 2 and 4 with groups of highly similar sequences (Fig. 3; Extended Data Fig. 4). To confirm these findings based on amino acid differences, we constructed tanglegrams comparing the hierarchical clustering of alpha and beta sequences (Fig. 6a; Extended Data Fig. 5a). These allelic groups were named as a combination of a number (locus) and a letter. In this way, 51 of 54 alpha and beta sequences from Locus 2 were clustered into five distinct groups, 2a-e (Fig. 6; Supplementary Fig.9b). Groups 2f-h contained only one sequence each, which could be due to small sample size in this study. Similarly for Locus 4, 32 sequences could be grouped into 4a and 4b while 4c contained a single sequence (Extended Data Figs. 4c, 5). The MHC class II *DA* genes at Locus 1 did not show such clear groupings of *DAA* and *DAB* alleles (Extended Data Fig. 4a), despite being as polymorphic as Locus 2 and 4 genes (Fig. 4a, b). Interestingly, *DAA2* sequences from allelic group 2b had a 12-bp in-frame insertion in exon 2 inserted after residue 33 encoding a tetrapeptide CTDC (Supplementary Fig. 6). The protein model with this insertion was predicted to be more stable (ΔΔG = -2.84 kcal/mol) than alleles lacking the insertion, suggesting that the additional disulfide bond likely contributes to a structural stabilization (Extended Data Fig. 6; Supplementary Figs. 7 and 8).

**Fig. 6.**
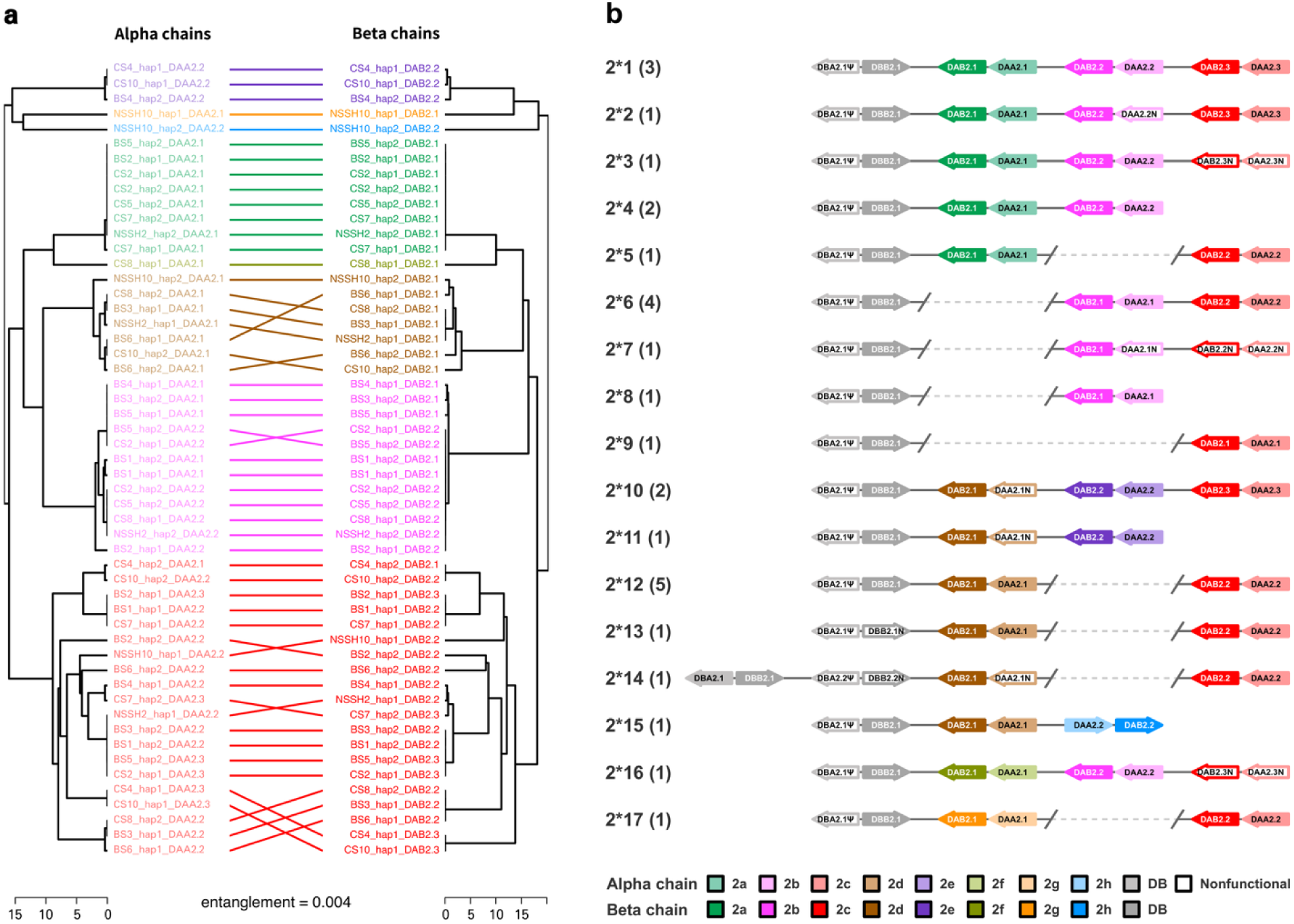
Allelic grouping and inferred structural haplotypes for MHC class II Locus 2. **a** Tanglegram illustrating association between *DAA* and *DAB* alleles in Locus 2 haplotypes. Each dendrogram represents hierarchical clustering based on pairwise percent amino acid differences. Correspondence between paired alpha and beta alleles is indicated by connecting lines. The entanglement measure, quantified using dendextend R package, is presented below the plot. Sequences are colored according to their respective allelic group. Alpha and beta sequences are colored in darker and lighter shades of the same color, respectively. The color code for the allelic groups is provided at the bottom right. **b** Seventeen haplotypes inferred from the arrangement of alpha-beta gene pairs based on their allelic groupings; numbers in parenthesis reflect the number of observations per haplotype. Gene and intergenic region lengths are shown at a fixed scale. *DBA* and *DBB* genes are displayed in light and dark gray, respectively. Nonfunctional alleles are represented by open arrows. Gene names are indicated within arrows. N, null allele; Ψ, pseudogene.

To assess whether alleles in different groups are randomly distributed across haplotypes or follow a pattern, we examined the allele groups present on each of the assembled haplotypes. This led to three interesting observations. First, each closely linked alpha-beta gene pair in the 28 haplotype-based assemblies belonged to matching alpha-beta allelic groups, suggesting a strong non-random association and co-evolution of alpha-beta gene pairs at these loci (Fig. 6a; Extended Data Fig. 5a). For instance, *DAA2.1* and *DAB2.1* on CS2_hap1 belonged to the corresponding alpha and beta allelic groups 2a. Second, each haplotype carried alpha-beta gene pairs from different allelic groups, thus maximizing the haplotype diversity (Fig. 6b; Extended Data Figs. 4b, c, 5). As example, the three alpha-beta gene pairs (*DA2.1, DA2.2, DA2.3*) in CS2_hap1 correspond to sequence groups 2a, 2b, 2e, respectively. This second observation was supported by the fact that intra-haplotype non-synonymous substitutions (*dN*) was higher than expected for a random arrangement of gene sequences (Extended Data Fig. 7). Third, specific combinations of allelic groups dominated at Locus 2 and 4. In case of Locus 2, 15 haplotypes harbored gene pairs from groups 2a, 2b, and 2c while 10 haplotypes harbored gene pairs from 2e, 2d, and 2c (Fig. 6b). Most of the Locus 4 haplotypes contained only single gene pair, of which 15 belonged to group 4a and 8 belonged to group 4b (Extended Data Fig. 5b). This suggests that MHC haplotypes in herring often consist of a specific sequence of allelic groups, which is 2a-2b-2e and 2c-2d-2e in case of Locus 2 and 4a, 4b, and 4a-4c in case of Locus 4 (Fig. 6b; Extended Data Fig. 5b; Supplementary Figs. 9, 10; Supplementary Text).

One of the most surprising findings was that several of the nonpolymorphic *DB* genes, as well as several of the highly polymorphic *DA*, contained mutations that are expected to disrupt function, and these were classified as null alleles (Supplementary Table 3). Moreover, null alleles within the same allelic group in Locus 2 carried the same inactivating mutation.

## Discussion

This study has revealed that the genetic diversity of MHC class II genes in Atlantic herring is likely more extensive than in any other vertebrate studied so far, a result consistent with pathogen-driven selection and the enormous population size of the species. To the best of our knowledge, this is the first study to explore MHC polymorphism in natural populations using long-read sequencing, an approach that facilitated the accurate determination of the number of genes per locus and of complete gene sequences. A previous attempt to take advantage of the extensive amount of whole genome, short read sequencing available in this species^11,13,21^ failed, because it was not possible to generate accurate locus-specific data even when using a data set comprising high coverage sequence from a full-sib pedigree generated to estimate the mutation rate in the Atlantic herring^10^. The extensive sequence diversity, copy number variation and the overlap between intra-locus and inter-locus sequence identities revealed in the present study explain why this previous attempt failed.

The human MHC class II region (*HLA*) is by far the most well studied in any vertebrate and includes some of the most polymorphic genes in the human genome (https://www.ebi.ac.uk/ipd/imgt/hla/). In this study, we explored how much diversity is present in a species that has population size and fecundity 1000-fold greater than that of humans, and with a much larger long-term effective population size the last million years. We now show that the class II diversity in Atlantic herring noticeably exceeds that humans. In humans, there are four highly polymorphic class II genes: *HLA-DPB*, -*DQA*, -*DQB,* and -*DRB*^22,23^, whereas the number of highly polymorphic genes are clearly higher in herring (Fig. 4). Furthermore, copy number variation occurs in human but is more prevalent in herring adding an additional layer of diversity. Sequence polymorphisms in herring, like in humans and other species, occur primarily in exon 2 encoding the peptide binding domain (Fig. 4a, b). Further, footprints of natural selection intensity are evidently more prominent in herring, where the *dN/dS* ratios for the most polymorphic genes are ∼2.0, twice as high compared with the corresponding human genes^24^ (Fig. 4e, f).

More studies using long-read sequencing are required to enhance our understanding of MHC diversity in natural populations. Previous reports based on short read data have usually underestimated the number of polymorphic class II genes. This is clearly the case for a previous study in Atlantic herring, which found only one class II gene^25^. Traditional approaches based on PCR, cloning, and targeted Illumina sequencing have been important in the past to shed light on MHC diversity, but those studies are error-prone and lead to mis-interpretations^26,27^. In these studies, the number of distinct loci is often based on maximum number of alleles observed in one individual and does not consider copy number variation and paralogs, leading to inaccurate and mostly inflated estimates of the number of loci. The report of the largest number of polymorphic loci in a teleost species is in Old world cichlids from five *O. niloticus* families (at least 17 MHC class IIB loci)^28^. Since this study lacks information on gene copy number and the presence of pseudogenes, the estimates are uncertain. In fact, none of the previous studies on teleosts have an accurate estimate of the number of MHC class II loci (Supplementary Table 2)^28–39^. We also note that the results for Atlantic herring are in stark contrast to the situation in the cod family^40^ and pipefishes^41^, in which the MHC class II genes have been lost.

The extensive class II polymorphism in Atlantic herring provides strongest support for the theory that MHC diversity is maintained by pathogen-driven natural selection^2,8^. The herring provides an ideal model for Hardy-Weinberg assumptions of random mating and infinite population size as is likely to be found for a vertebrate. A consequence of the large population size and high fecundity is that genetic drift is close to non-existent and that the frequencies of alleles affecting fitness are determined by natural selection^42^. The schooling behavior of herring and its abundance make it an attractive target for pathogens. In fact, there is some evidence that pathogen richness promotes an increase in the number of MHC genes^43^, as here noted in the Atlantic herring. Two pathogens, viral hemorrhagic septicemia virus (VHSV) and a fungal-like protist, *Ichthyophonus hoferi*, are major threats to Atlantic herring populations^44^. VHSV genotype Ib is notably prevalent among Norwegian spring-spawning herring^45^. Infection by *I. hoferi* has contributed to mass mortality of Atlantic herring from both sides of the North Atlantic Ocean^46,47^ and steep declines in the landings in the Gulf of St. Lawrence^48^. A study comparing MHC genotypes among affected and non-affected individuals as regards these major pathogens would be of considerable interest, but the extremely high polymorphism implies that such studies need to be very large to give a reasonable power to document associations.

The current study shows that a role for MHC genes in mate choice is not a requirement for maintaining extremely high levels of MHC polymorphism, because herring is a broadcast spawner lacking mate choice. MHC-based mate choice may be a secondary effect that evolves when MHC polymorphism is present due to pathogen-driven selection. Similar conclusions based on experimental data have been reported in other species^49,50^.

It has been hypothesized that the MHC diversity at the individual level is restricted to prevent T-cell receptor (TCR) depletion, as too high diversity of MHC molecules may cause too many thymocytes to fail to be positively selected during T-cell maturation in thymus^51,52^. Although how this intricate balance between MHC and TCR loci occur is still debated, a recent study in bank voles suggest that an intermediate level of MHC diversity is required to maintain optimal number of TCRs^53,54^. It is possible that herring MHC diversity is at that balancing point since we found both considerable copy number variation (Fig. 2, Table 1, Supplementary Fig. 2), higher within-haplotype diversity than expected by random chance (Extended Data Fig. 7), but we also paradoxically found null alleles at many of the highly polymorphic genes (Supplementary Table 3) causing a reduction of the diversity of antigen-presenting molecules.

We were able to categorize almost all alleles on Locus 2 and Locus 4 into a handful of major groups using sequence similarity matrices (Fig. 6; Extended Data Figs. 4b, c and 5). These could be referred to as supertypes, a concept developed for human MHC genes to group alleles sharing similar physiochemical properties at peptide binding residues^55^. We noted strong non-random association of MHC class II alpha-beta gene pairs at these loci, which is a classical feature of class II polymorphisms; alpha and beta chains together form the peptide binding site and certain combinations are more fit than others. Furthermore, the alpha-beta gene pairs in each haploid assembly carried different sets of allelic groups (supertypes) at Locus 2 and 4. Such arrangement maximizes the diversity at the haplotype level, implying a broad peptide binding repertoire per haplotype (Extended Data Fig. 7). This supports the premise that evolution favors MHC haplotypes with different sets of supertypes and addresses the questions of why certain allelic variants are maintained together in haplotypes^56^. It also supports the theoretical premise that supertypes are maintained as balanced selection under a Red Queen dynamic model which allows fluctuation in frequencies of polymorphic alleles, while alleles within the supertypes are maintained by positive selection under a Red Queen arms race model of evolution which promotes fixation of favorable alleles^57^. Studies have shown that the presence of specific supertypes instead of alleles is associated with resistance to parasites^58–60^.

Our previous short-read, whole genome sequence analyses of a large number of Atlantic herring populations have resulted in the identification of hundreds of loci underlying ecological adaptation to differences in water salinity, spawning time, temperature^11,21,61^, and light conditions^62^. It is obvious that the MHC class II regions explored in this study have been blind spots in these analyses due to the difficulty in accurately scoring genetic diversity based on short read sequencing an issue shared with other complex immune loci^63,64^. The current reference assembly is to some extent collapsed in MHC regions with duplicated gene copies and the overlap between intra- and inter-locus sequence identities make the inference of genetic polymorphism challenging. This study is based on the combined analysis of 28 haplotypes from 14 individuals which resulted in the identification of up to 28 different alleles at the most polymorphic genes. This implies that large-scale studies will be required to reveal how MHC polymorphism in Atlantic herring contributes to fitness. However, it is possible to interpret the extensive amount of short read data in the context of the results of this study, for instance as regards the segregation of major supertypes at Locus 2 and 4. Furthermore, the cost for long-read sequencing has decreased steadily in recent years, making it possible to perform large scale studies in the future and combine these with data on pathogen susceptibility in this extremely abundant marine fish.

## Materials and Methods

### Samples and sequencing

The samples included a total of 14 Atlantic herring individuals, 6 from the Celtic Sea (collected on November 11, 2019 at latitude N51°59′ and longitude W6°48′), 6 from the Baltic Sea (collected on May 18, 2020 in the Baltic Sea, Hästskär, at latitude N60°35′ and longitude E17°48′), and 2 from the Norwegian Spring spawning stock (collected on February 21, 2021 in the Norwegian Sea at latitude N67°55′ and longitude E11°18′). High molecular weight was prepared from tissue samples and subjected to PacBio HiFi long-read sequencing as previously described^65^. *De novo* genome assemblies were constructed using hifiasm^12^. The resulting high-quality 28 haploid PacBio genomes and the reference genome^13^ were used for further analysis. All PacBio contig-level assemblies were scaffolded using the chromosome-level reference assembly and the RagTag software v2.0.1^66^.

### Annotation of MHC class II genes

Due to the complex structure of the MHC genomic regions, the ENSEMBL annotation of MHC class II genes on the reference assembly was incomplete with respect to open reading frames (ORFs), intron-exon structure, distinction between alpha and beta genes, and unannotated genes. Thus, we manually annotated the genes first by BLAST^67,68^ using MHC II class genes from other species as queries (78 sequences from 16 species; Supplementary File 2) and visualized the BLAST hits on the genome using IGV^69^ to obtain the approximate location of MHC class II genes in the herring genome. To obtain the accurate gene coordinates, we used previously generated RNA-seq data from eight tissues^21^. We aligned RNA-seq data against the reference genome using STAR^70^ and visualized it using IGV^69^ to locate ORFs, intron-exon junctions, and stop codons. Sequences which had the same gene structure as other MHC class II genes, and which were not disrupted by inactivating mutations were annotated as functional genes.

After annotating MHC class II genes on the reference assembly, we lifted over these annotations to the new haplotype assemblies using a genome annotation lift-over tool, LiftOff^71^. However, due to the complexity of MHC genes, nearly 30% of the LiftOff annotated genes were either incomplete or had wrong intron-exon structure. LiftOff also failed to annotate all genes. Hence, to correct LiftOff annotations and obtain complete annotation, we used previously generated RNA-seq data from spleen^37^ and aligned these against all new assemblies using HiSat^72^. We also used BLAST using the annotated genes as query. LiftOff annotations, RNA-seq reads, and BLAST hits for each PacBio genome were visualized using IGV and all annotations were curated manually. If a gene was inactivated in all haplotypes it was classified as a pseudogene. We found only one pseudogene, *DBA2.1*, carrying a one bp deletion (nt G) at the end of exon 2. Alleles with nonsense or indel mutations disrupting the reading frame, or incomplete gene structure were referred to as ‘null ‘alleles. Supplementary Table 3 contains a list of all null alleles and their inactivating mutations. Genomic organization of the annotated genes on each assembly was plotted using gggenomes (https://github.com/thackl/gggenomes). MHC regions and the flanking sequence between haplotypes were aligned using MUMmer^73^.

From the genomic and gene organization (Figs. 1, 2; Supplementary Figs. 1, 2), we observed assembly artifacts for samples CS7 and CS4. In the case of CS7, one haplotype assembly (haplotype 2) lacked genes from one locus (Locus 1) but the alternative haplotype from the same individual carried two extra copies. This was most likely caused by incorrect phasing by hifiasm. We therefore decided to make the most conservative interpretation that both haplotypes contained Locus 1 genes than to infer that this individual carried two unique haplotypes, one with zero and the other with five copies of Locus 1 genes. In case of CS4, one haplotype (hap2) lacked Locus 4 genes. It is possible that this represents an assembly artifact but in the absence of evidence for this we did not make any manual correction of this haplotype.

Locus 1 contained two alpha-beta gene pairs, named as DA1.1 and DA1.2. However, nine new assemblies had a third pair in between DA1.1 and DA1.2 where alpha gene had an unusually long intron 1 ranging between 18-23 kb (Supplementary Table 4), indicating that these nine new assemblies share the same haplotype for Locus 1 (which in fact resulted to be true from the allele-clustering analysis (Extended Data Fig. 4a)). To distinguish this different gene structure, we named this extra DA gene pair as DA1.3. However, BS2_hap1 DA1.3 did not have long intron1, but we did not change its naming in order to keep the consistency with the rest of the naming.

### Gene sequence analysis

Sequences were extracted from the genomes using gffread^74^ for various analysis using custom scripts available on GitHub page. Phylogenetic trees of alpha and beta MHC class II coding sequences from only Atlantic herring reference assembly were constructed using the neighbor-joining method in R package ape^75^. Phylogenetic analysis of representative MHC class II sequences from Atlantic herring, along with sequences from tetrapod and fish species retrieved from public databases and a previous study, was conducted separately for the α- and β-chains using full-length amino acid sequences. A list of MHC class II α- and β-chain sequences from other species, along with their corresponding accession numbers, is provided in Supplementary Table 5. Phylogenetic trees were constructed using the maximum-likelihood method implemented in IQ-TREE^76,77^, with bootstrap support values^78^ estimated from 2,000 replicates. All trees were visualized using R package ggtree^79^. Nucleotide diversity (*π*) was calculated using R package pegas^80^ where sequences were aligned with ClustalO in the R package msa^81^ for all four exons of all genes. Analysis for the ratio of non-synonymous substitutions over synonymous substitutions (*dN/dS*) for exon 2 and 3 was carried out using the software MEGA 11^82,83^ with the Nei-Gojobori method (Proportion) with 2000 bootstrap replicates. Substitution rates were modeled with gamma-distributed rate variation across sites and gaps/missing data was treated under pairwise deletion model. Amino acid sites under positive selection were estimated by Bayes Estimation in the models for positive selection in PAML, M2 and M8^16^. Peptide binding residues (PBRs) in Atlantic herring MHC class II genes were predicted by comparison of Atlantic herring and tetrapod sequences from previous literature^18,19^.

### Protein structure modelling

The protein structure modelling was performed using AlphaFold3^84^ on two alpha-beta gene pairs, *CS4_hap1_DA2.1* and *BS1_hap1_DA2.1* to explore the location of the most polymorphic amino acid residues and the functional consequences of the four amino acid-insertion (CTDC) in the class II alpha chain, respectively. AlphaFold predicted five models for each protein, and the highest-ranking model of each protein was selected for further analysis. The transmembrane regions were predicted using DeepTMHMM 1.0^85^ (Supplementary Figs. 5a, b and 8a, b). Signal peptides of alpha and beta chains were predicted using SignalP 6.0^86^ with settings for eukarya and the slow model mode (Supplementary Figs. 5c, d and 8c, d). The signal peptide annotation for *CS4_hap1_DAA2.1* was predicted with relatively low confidence (probability of 0.67), resulting in differing cleavage site predictions between DeepTMHMM and SignalP (Supplementary Fig. 5a, c). Molecular graphics and figure rendering were performed with UCSF ChimeraX v1.9^87^ and PyMOL v2.3.0^88^.

We used Shannon entropy^89^ as a measure to quantify the degree of polymorphism at each amino acid residue using all MHC class II DA alpha and beta genes from Locus 2. The Shannon entropy values were calculated using R package bio3d^90^. The values range from 0 to 4.32 where 0 represents a site with only one amino acid (highly conserved) while 4.32 represent a site with all 20 amino acids (highly polymorphic). To represent these values on a protein structure, we used normalized values by diving all values with the maximum.

To understand the effect of the CTDC insertion on the protein structure in the alpha chain, we substituted Cys-33 in the CTDC insert of the alpha chain to an alanine (C33A) to break the disulfide bond. The stability of the resulting protein model was analyzed using DDMut^91^.

### Haplotype sequence analysis

To assess amino acid sequence similarity in our dataset of 195 *DAA*, 220 *DAB*, 53 *DBA*, and 107 *DBB* sequences, we performed hierarchical clustering and heatmap visualization. Full-length amino acid sequences were aligned within each gene category using the MAFFT multiple sequence alignment program^92^. Pairwise distance matrices were computed based on percent amino acid mismatches, with insertions and deletions counted as mismatches. These distance matrices were used to construct hierarchical clustering using the UPGMA method, and the results were visualized as heatmaps with the ComplexHeatmap R package^93^.

To evaluate intra-locus diversity and investigate haplotype structure, we performed locus-specific analyses for polymorphic genes at loci 1, 2 and 4. For these analyses, *DAA* and *DAB* sequences were clustered separately using the same alignment and distance matrix approach described above for the genome-wide analysis. The resulting dendrograms for each locus were then compared using tanglegrams, generated with the dendextend R package^94^. Dendrograms were aligned to optimize tip correspondence, and entanglement scores were calculated to quantify the similarity between *DAA* and *DAB* clustering patterns at each locus.

We conducted further haplotype analysis for loci 2 and 4, as these loci showed clear clustering patterns among sequences. We assessed homology across intronic and intergenic regions. Pairwise comparisons of the entire locus sequences, including 20 kb of flanking sequence on either side, were performed across 28 phased haploid assemblies. Alignments were generated using NUCmer from the MUMmer package^73^, and the resulting dotplots were constructed with mummerplot and visualized in R. Structural haplotypes for loci 2 and 4 were inferred based on (1) amino acid sequence clustering, as visualized with tanglegrams, and (2) pairwise alignment patterns across the entire locus, as visualized using dotplots . Null alleles, which were excluded from amino acid sequence clustering due to the absence of open reading frames (ORFs), were assigned to allelic groups solely based on homology patterns in the dotplots. Haplotype schematics, depicting gene order and orientation with fixed gene lengths and intergenic intervals, were generated in R using the gggenes package^95^.

## Author contributions

LA conceived the study. MJ and FMS performed all bioinformatic analysis. ML performed protein structure modelling and analysis. FB contributed to sample collection. BWD, JL and MEP provided advice for the bioinformatic analysis. MFC provided advice as regards MHC biology. MJ, FMS and LA wrote the paper with input from other authors. All authors approved the paper before submission.

## Data availability statement

The sequence data generated in this study is available from Bio project PRJNA1023520.

## Code availability statement

The analyses of data have been carried out with publicly available software and all are cited in the Methods section. Custom scripts used are available in https://github.com/LeifAnderssonLab/ MHC_diversity

## Acknowledgements

The study was supported by the Knut and Alice Wallenberg Foundation (KAW 2016.0361) and Vetenskapsrådet (2017-02907). The National Genomics Infrastructure (NGI)/Uppsala Genome Center provided service in massive parallel sequencing and the computational infrastructure was provided by the Swedish National Infrastructure for Computing (SNIC) at UPPMAX partially funded by the Swedish Research Council through grant agreement no. 2018-05973.

## Extended Data

**Extended Data Fig. 1.**
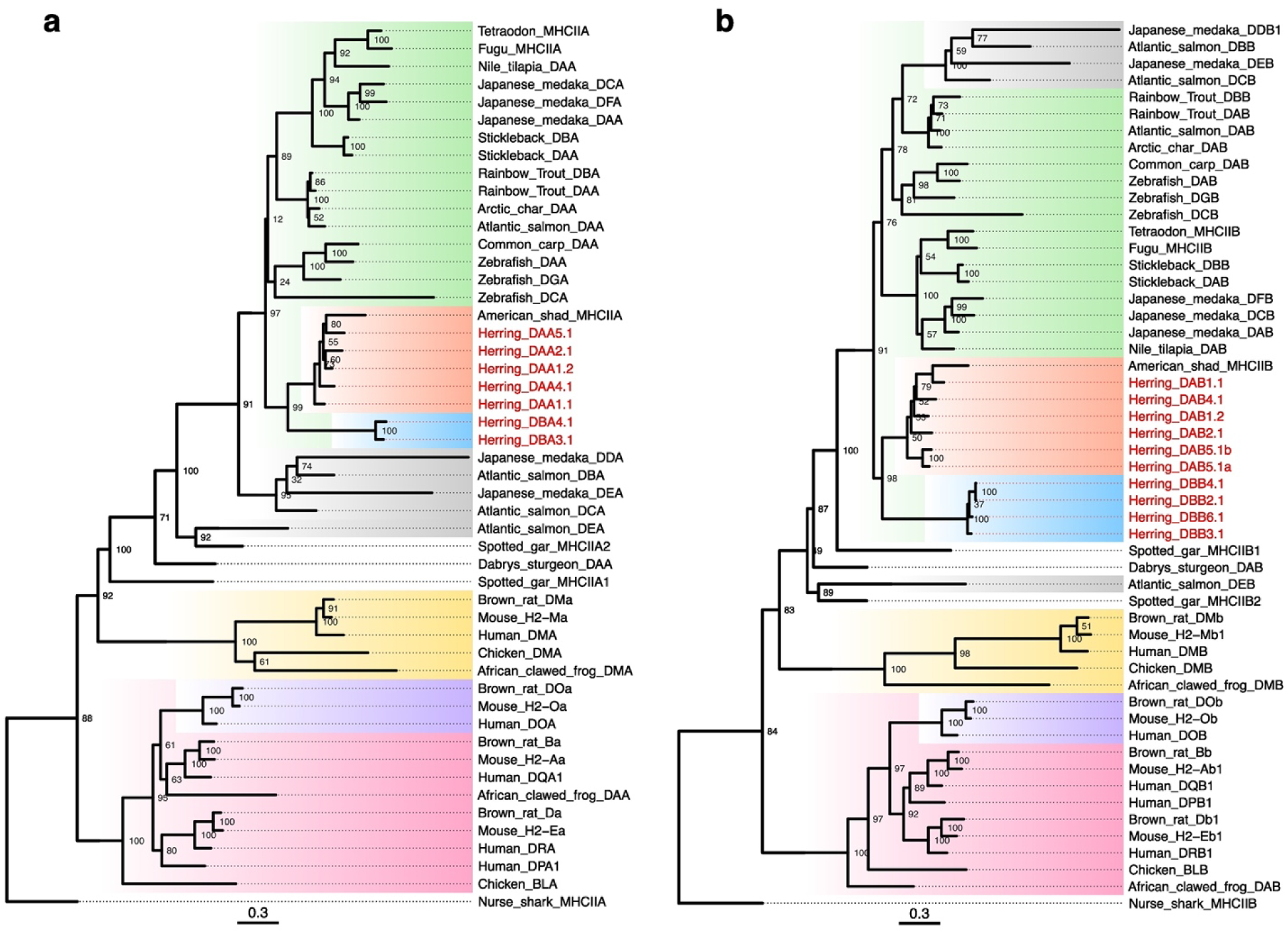
Maximum-likelihood phylogenetic tree based on full-length amino acid sequences of representative MHC class II genes from fish and tetrapod clades, along with sequences from the Atlantic herring reference genome. **a** alpha genes. **b** beta genes. Clades are highlighted as follows: green for teleost DA/DB lineages, red for Atlantic herring DA lineage, blue for Atlantic herring DB lineage, gray for teleost DE lineage, yellow for tetrapod DM lineage, purple for tetrapod DO lineage, and pink for classical tetrapod sequences. Atlantic herring sequences generated in this study are marked with red tip labels. Node labels represent bootstrap support values inferred from 2,000 replicates. The scale bar indicates branch length.

**Extended Data Fig. 2.**
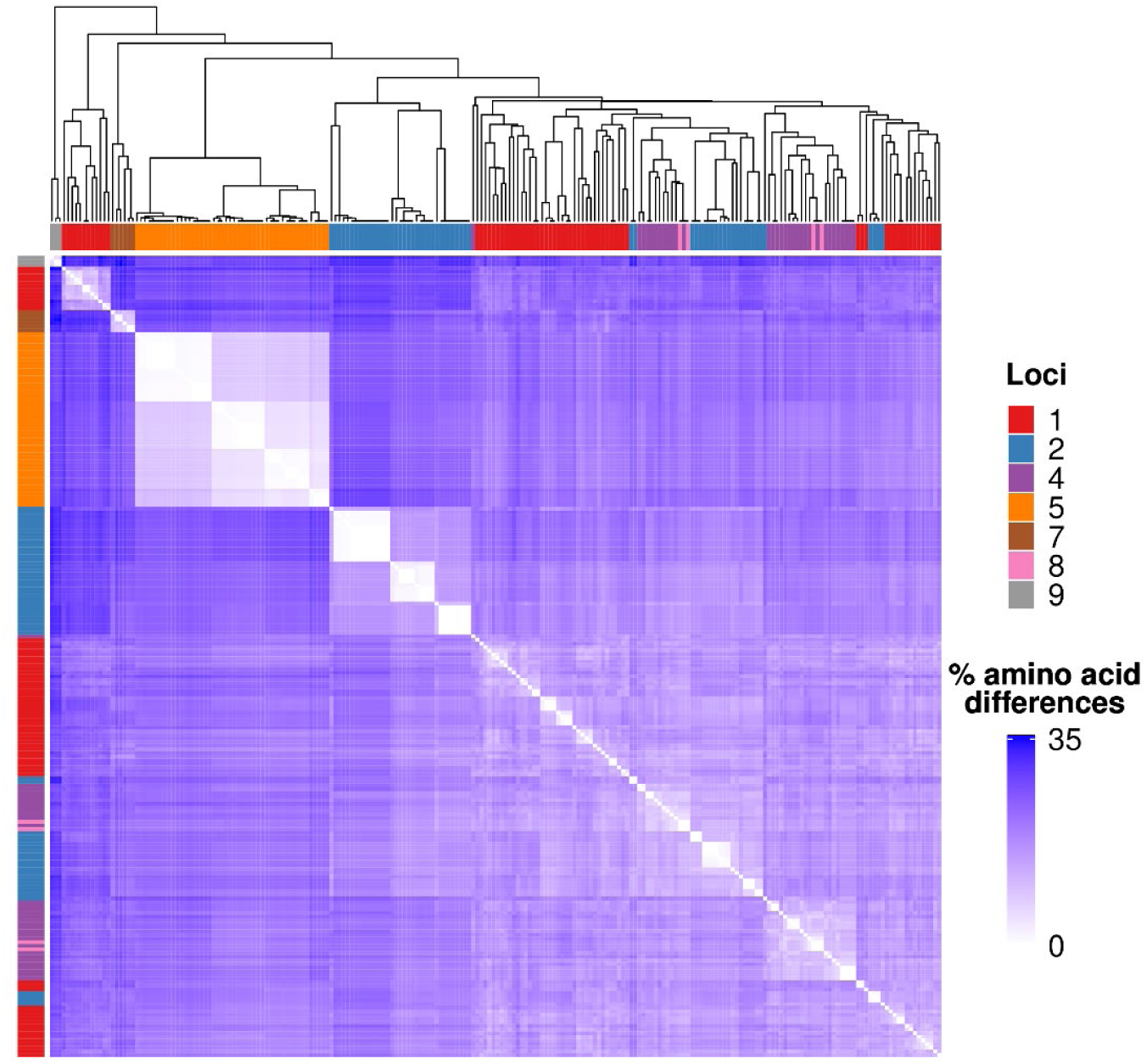
Clustering of amino acid sequences from DAB genes. **a** Heatmap and dendrogram showing hierarchical clustering of all DAA amino acid sequences across the genome. Heatmap gradients correspond to pairwise percent amino acid differences. Source loci are color-coded in the annotation bar

**Extended Data Fig. 3.**
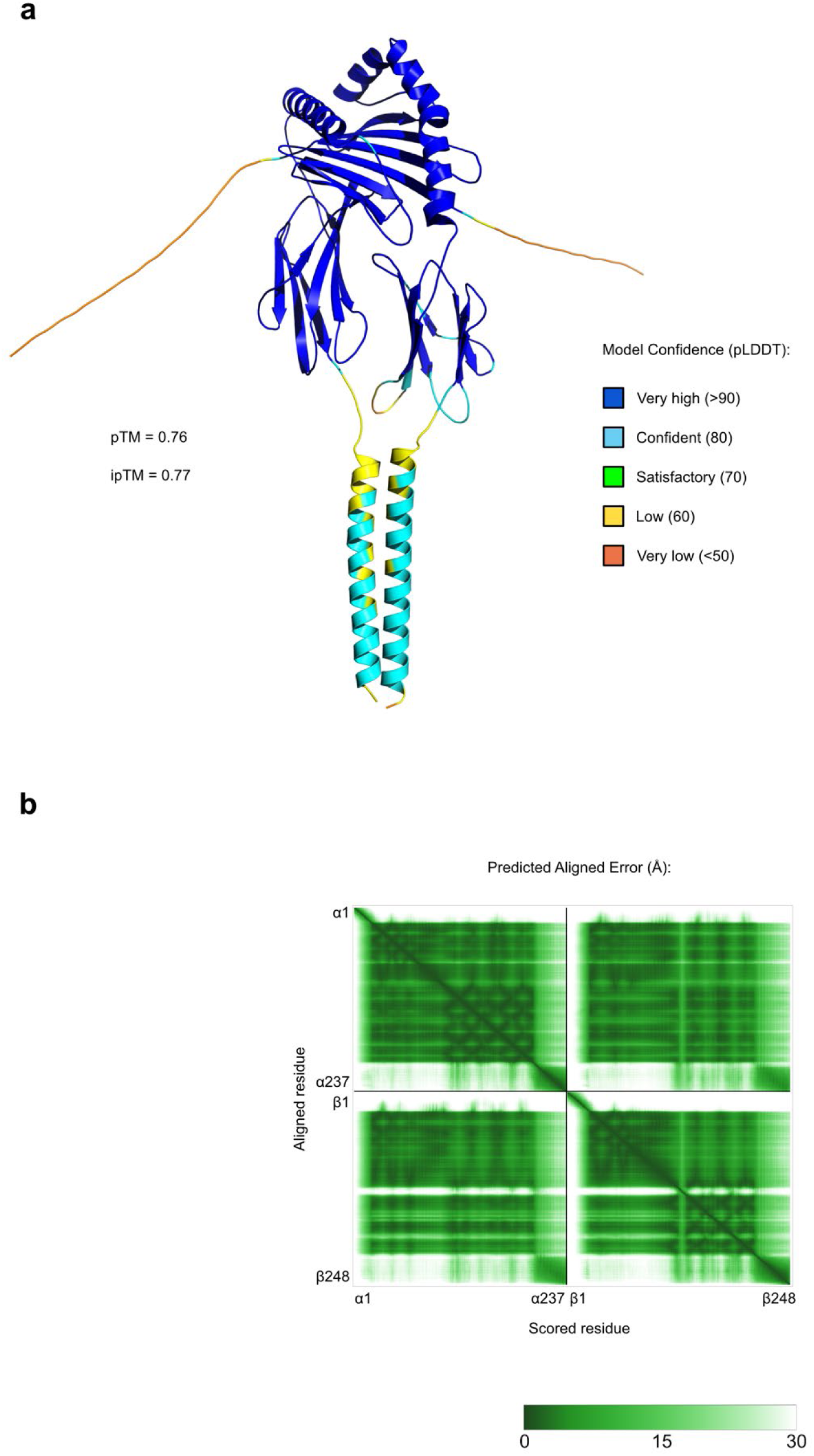
Predicted protein structure of the MHC class II heterodimer formed by *CS4_hap1_DAA2.1* (alpha chain) and *CS4_hap1_DAB2.1* (beta chain). **a** Model colored with the predicted local distance difference test values (pLDDT) of the alpha and beta chains. The predicted template modeling score (pTM) and interface predicted template modeling score (ipTM) are annotated. **b** Predicted aligned error (PAE) of the model. Amino acid residue number of alpha and beta chains are annotated by the axes of the plot.

**Extended Data Fig. 4.**
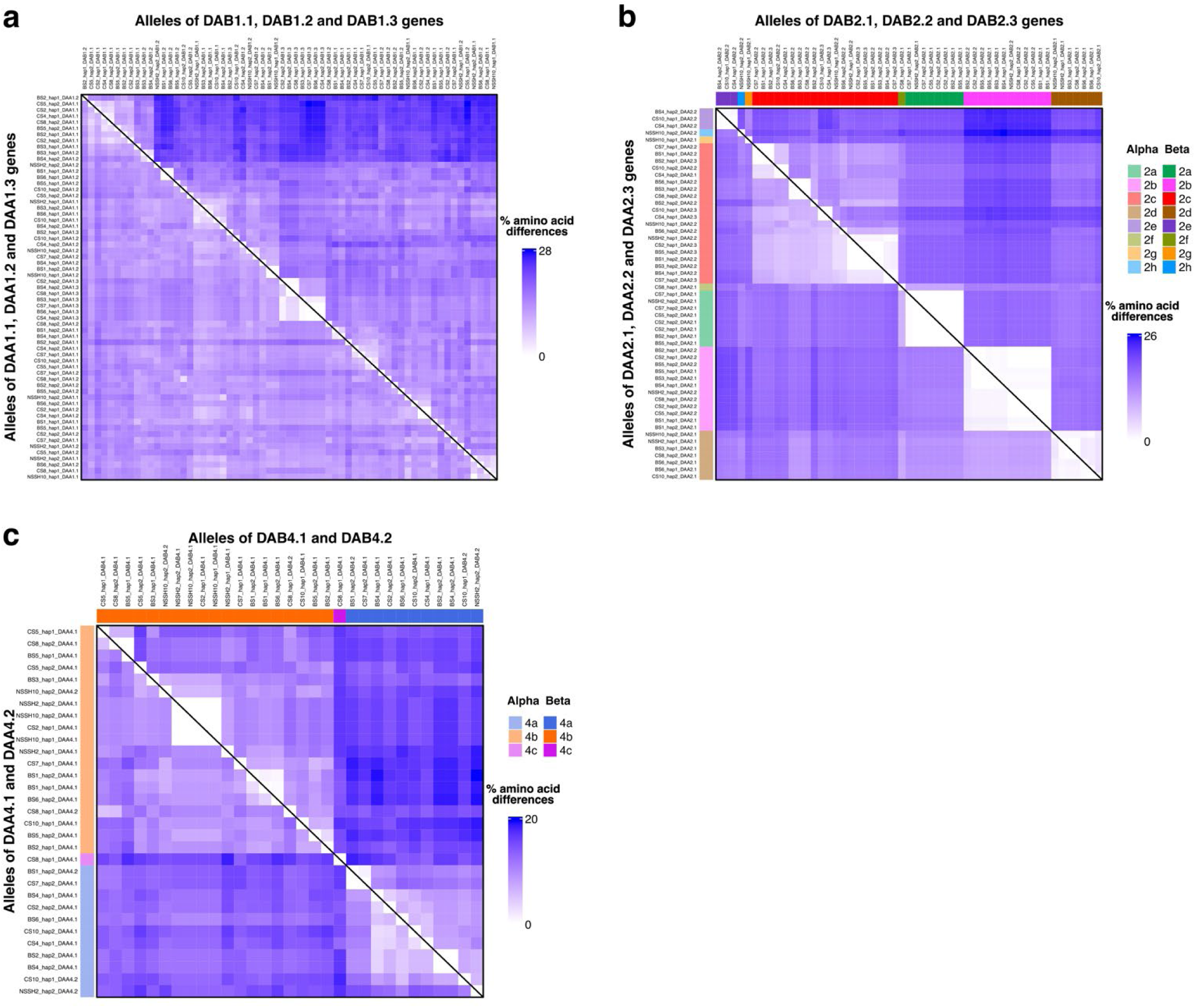
Heatmap of amino acid sequences from alleles at **a** Locus 1, **b** Locus 2, and **c** Locus 4. Upper triangle contains alleles of *DAB* genes and lower triangle contains alleles of *DAA* genes. Locus 2 and Locus 4 alleles are grouped into allelic groups indicated by different colors.

**Extended Data Fig. 5.**
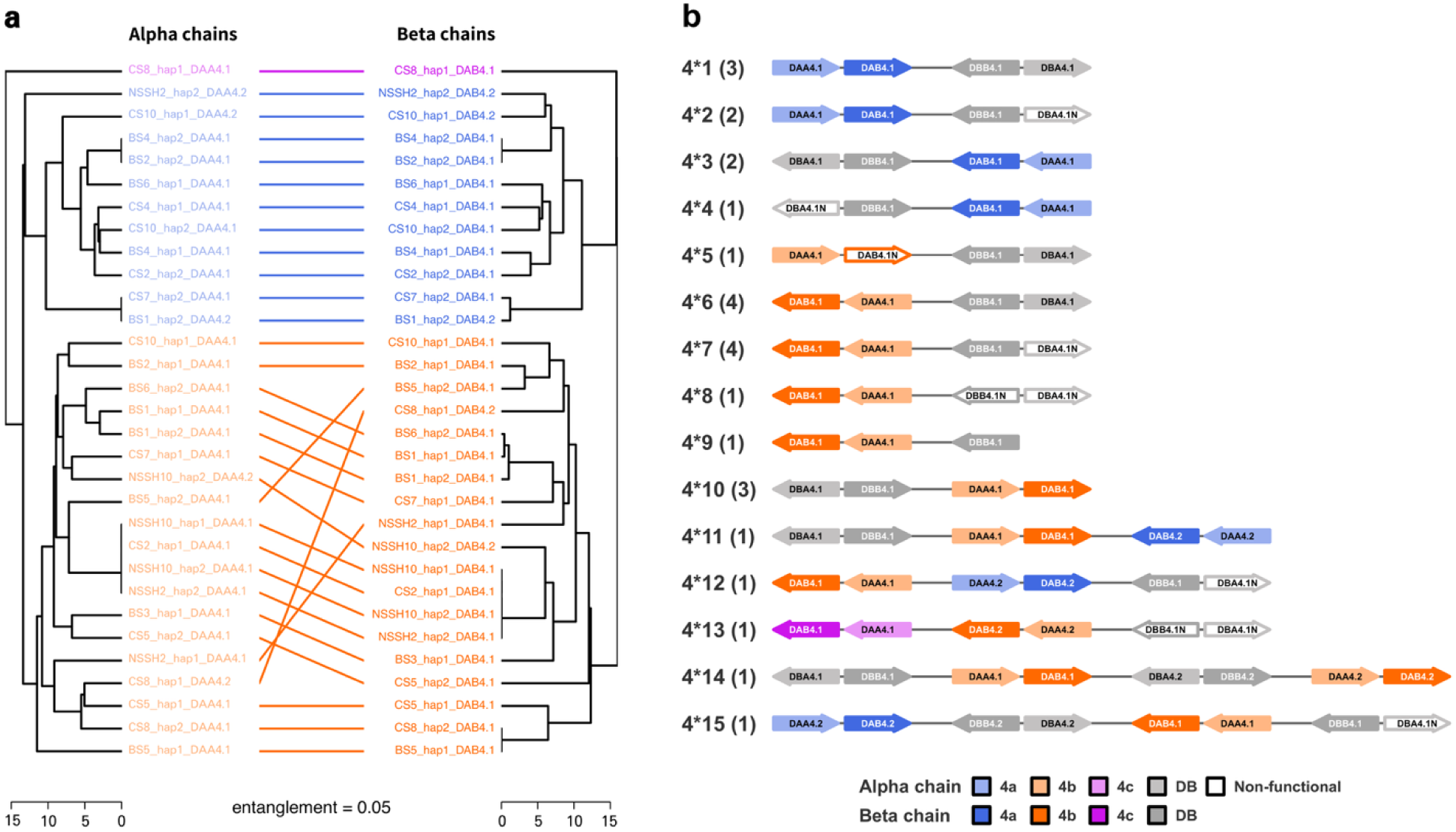
Allelic grouping and inferred structural haplotypes for MHC class II 4. **a** Tanglegram illustrating the alignment between alpha and beta amino acid sequence clustering. Each dendrogram represents hierarchical clustering based on pairwise percent amino acid differences. Correspondence between paired alpha and beta alleles is indicated by connecting lines. The entanglement measure, quantified using dendextend R package, is presented below the plot. Sequences are colored according to their respective allelic group. Alpha and beta sequences are colored in darker and lighter shades of the same color, respectively. The color code for the allelic groups is provided at the bottom right. **b** Fifteen structural haplotypes inferred from the arrangement of alpha-beta gene pairs based on their allelic groupings; numbers in parenthesis reflect the number of observations per haplotype. Gene and intergenic region lengths are shown at a fixed scale. *DBA* and *DBB* genes are displayed in light and dark gray, respectively. Nonfunctional alleles are represented by open arrows. Gene names are indicated within arrows. N, null allele.

**Extended Data Fig. 6.**
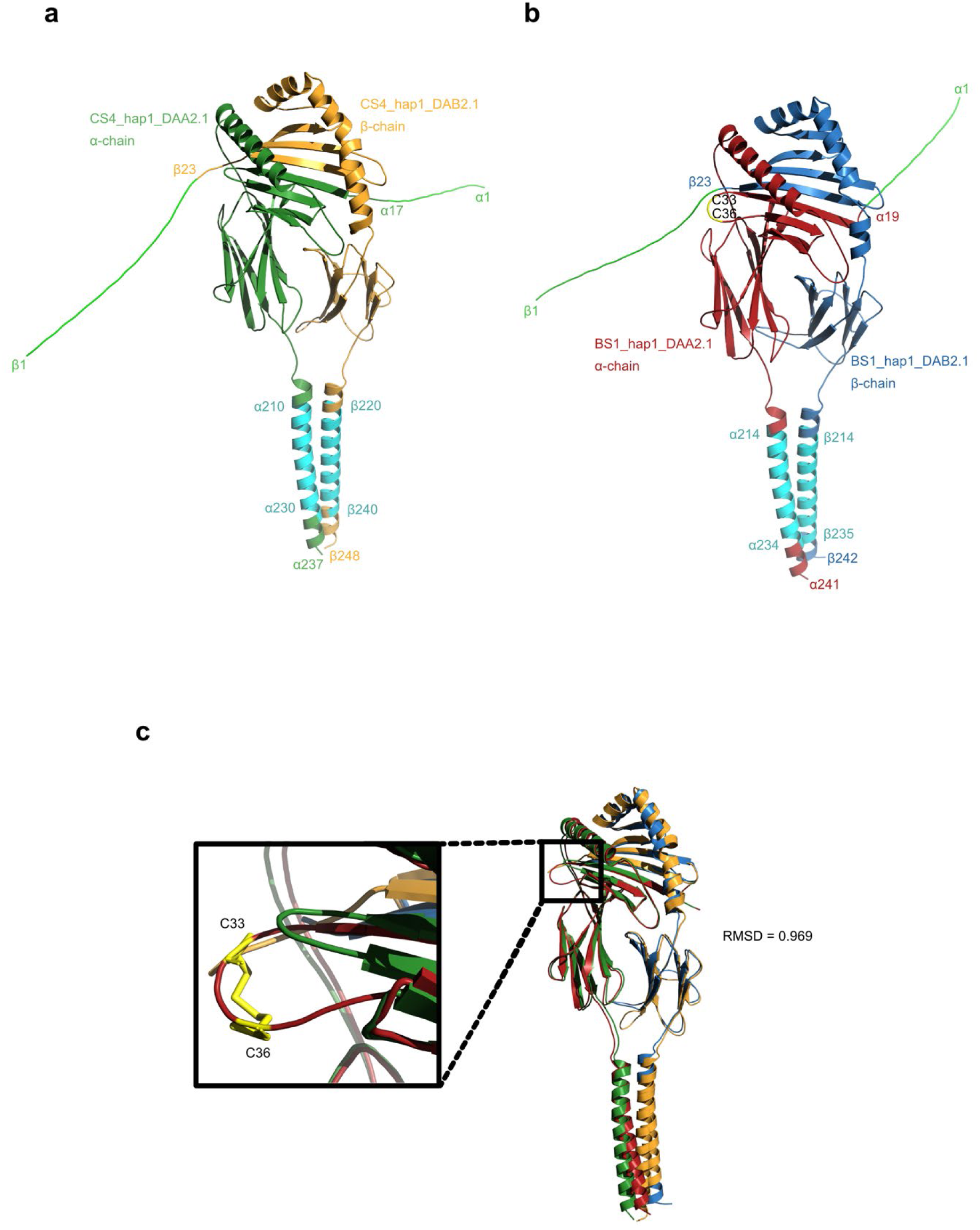
Predicted structures of two herring MHC class II proteins. **a** *CS4_hap1_DA2.1* heterodimer, where alpha and beta chains are colored in green and yellow, respectively. Predicted signal peptides and transmembrane regions are colored in light green and cyan, respectively. Borders of different regions are annotated and numbered according to the respective amino acid residue position. **b** Predicted structure of *BS1_hap1_DA2.1* heterodimer, where alpha and beta chains are colored in red and blue, respectively. Predicted signal peptides and transmembrane regions are colored same as **a**. The CTDC insert between the beta sheet 1 and beta sheet 2 of the alpha chain is colored in light yellow. Borders of different regions are annotated and numbered according to the respective amino acid residue position. **c** Structural alignment of *CS4_hap1_DA2.1* and *BS1_hap1_DA2.1*. Inset shows a magnified view of the disulfide bond formed between Cys-33 and Cys-36, highlighted in light yellow within the CTDC insert of *BS1_hap1_DAA2.1* (red), compared to the shorter loop of *CS4_hap1_DAA2.1* (green). The root mean square deviation (RMSD) of the structural alignment is annotated.

**Extended Data Fig. 7.**
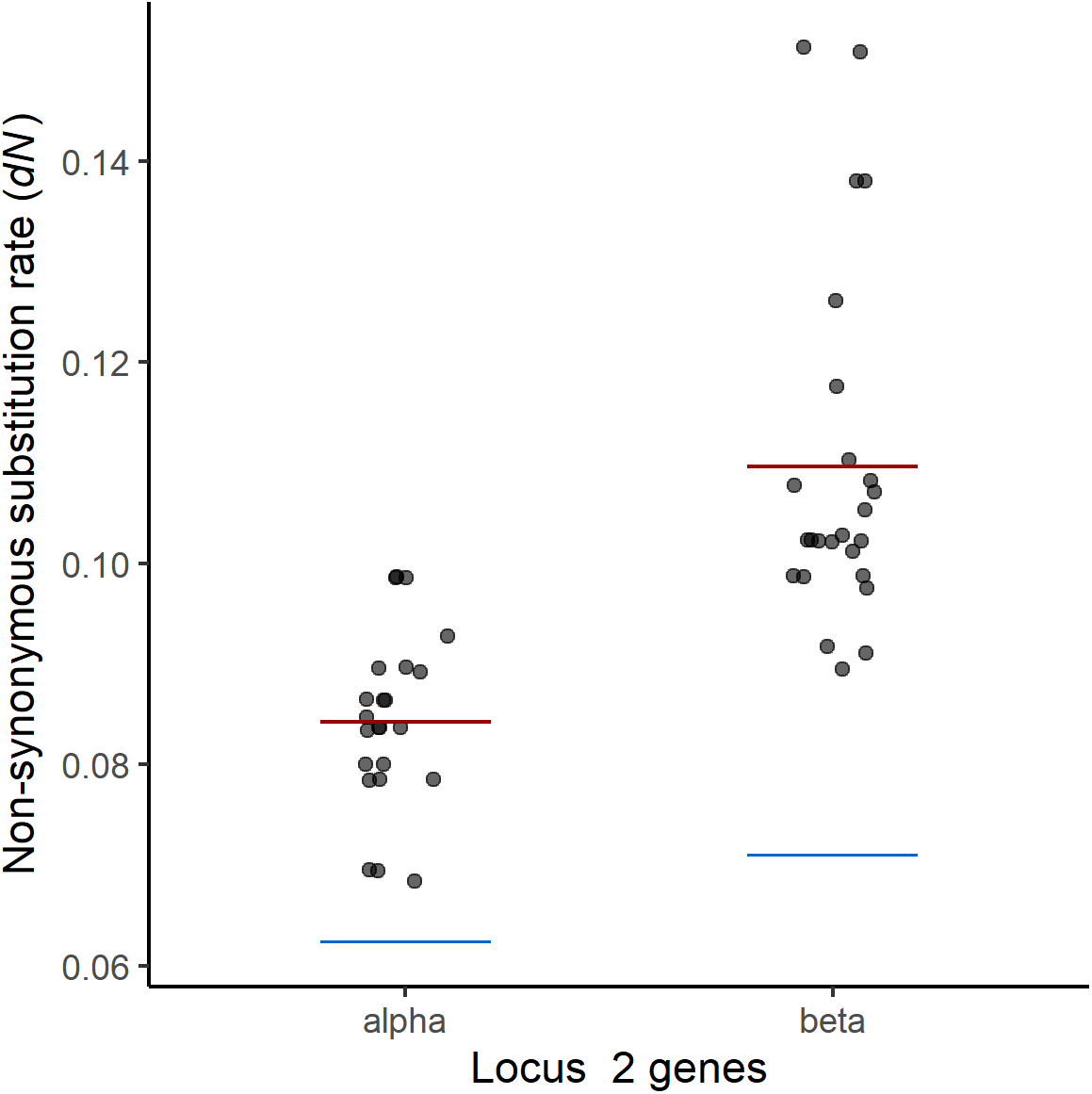
The rate of non-synonymous substitutions (*dN*) for Locus 2 haplotypes for alpha and beta genes. The average diversity is indicated by red line. The blue line indicates the mean rate for all genes irrespective of their haplotype status.

**Extended Data Table 1.** Genome statistics for Celtic Sea (CS), Baltic Sea (BS) and Norwegian spring-spawning herring (NSSH) PacBio genome assemblies using hifiasm assembler. Hap1 and hap2 are the two haplotype genomes and their statistics are shown on the top and bottom row for each sample, respectively.

| **Samples** | **Genome size**<br>**(Mb)** | **Genome**<br>**completeness (%)** | **No. contigs** | **N50 (Kb)** |
| --- | --- | --- | --- | --- |
| **CS2** | 769.1 | 91.5 | 4364 | 524.8 |
|  | 744.6 | 91.6 | 3314 | 547.9 |
| **CS4** | 776.3 | 92.8 | 3253 | 693.1 |
|  | 751.3 | 92.8 | 2539 | 731.1 |
| **CS5** | 768.1 | 91.6 | 3817 | 545.0 |
|  | 745.9 | 91.6 | 3037 | 569.8 |
| **CS7** | 783.6 | 92.0 | 3929 | 469.5 |
|  | 751.1 | 90.6 | 3117 | 512.3 |
| **CS8** | 781.7 | 91.9 | 4100 | 526.5 |
|  | 743.2 | 91.5 | 3280 | 504.5 |
| **CS10** | 773.1 | 90.7 | 3997 | 452.8 |
|  | 756.6 | 90.3 | 3228 | 472.4 |
| **BS1** | 776.2 | 91.6 | 3756 | 576.2 |
|  | 751.8 | 91.4 | 3028 | 606.5 |
| **BS2** | 788.3 | 91.8 | 3769 | 540.9 |
|  | 759.7 | 90.7 | 2952 | 539.8 |
| **BS3** | 779.2 | 92.4 | 3331 | 680.2 |
|  | 758.5 | 92.1 | 2725 | 661.8 |
| **BS4** | 792.2 | 92.8 | 3309 | 737.1 |
|  | 760.1 | 92.0 | 2570 | 735.4 |
| **BS5** | 780.3 | 90.6 | 4045 | 494.0 |
|  | 751.8 | 91.1 | 3259 | 480.0 |
| **BS6** | 787.8 | 92.2 | 3771 | 572.2 |
|  | 764.3 | 92.3 | 3083 | 557.1 |
| **NSSH2** | 782.5 | 92.6 | 3611 | 612.5 |
|  | 763.2 | 93.2 | 2961 | 634.9 |
| **NSSH10** | 792.9 | 94.2 | 2410 | 115.8 |
|  | 779.1 | 93.5 | 1806 | 127.5 |

## Supplementary Information

**Supplementary Fig. 1. Genome organization of MHC II genes on 29 haploid assemblies.** Alpha-beta gene pairs are colored based on their respective locus. \**DA1.4* and *DA1.5* genes on CS7_hap1 are misassembled and most likely present on CS7_hap2 as *DA1.1* and *DA1.2*, respectively (detail explanation in methods).

**Supplementary Fig. 2. Gene organization of MHC class II on nine loci in all 29 haplotypes**.

**Supplementary Fig. 3. Amino acid sequence alignments of α1, α2, β1, and β2 domains of DA and DB MHC class II sequences used for *dN/dS* analysis.** Colored residues represent positively selected sites predicted by M8 model of CODEML. The color shades represent the chemistry of amino acids where hydrophobic residues are in blue shades and hydrophilic residues are in red shades.

**Supplementary Fig. 4.** Amino acid sequence alignments of representative Atlantic herring sequences alongside sequences from fish and tetrapods, shown for the **a**, α1 domain and **b**, β1 domain. Sequence names are color-coded as in Extended Data Fig. 1: green for teleost DA/DB lineages, red for Atlantic herring DA lineage, blue for Atlantic herring DB lineage, gray for teleost DE lineage, yellow for tetrapod DM lineage, purple for tetrapod DO lineage, and pink for classical tetrapod sequences. Amino acid positions highlighted in green correspond to teleost-specific conserved cysteine residues in *DAA* genes, and fully conserved cysteine residues in *DAB* genes, that form disulfide bonds. Highly conserved residues involved in hydrogen bonding with the backbone of antigenic peptide are shaded in light red. Residues highlighted in yellow mark a four-amino-acid insertion (CTDC) in herring *DAA* sequences, which forms an additional disulfide bridge. Dashes represent gaps introduced for alignment. A chemistry-based coloring scheme is used to indicate amino acid chemical properties. The scale above the alignment indicates amino acid positions, beginning with the first residue of exon 2 in the full-length Atlantic herring protein. The black and red squares below the plots represent human^18^ and chicken PBRs^19^, respectively.

**Supplementary Fig. 5.** Metrics of predicted MHC class II protein model formed by *CS4_hap1_DAA2.1* (alpha chain) and *CS4_hap1_DAB2.1* (beta chain). **a, b** Results from SignalP 6.0 signal peptide prediction of the alpha chain and beta chain, respectively. **c** and **d**, Results from DeepTMHMM transmembrane region prediction of the alpha chain and beta chain, respectively.

**Supplementary Fig. 6.** Amino acid alignment of α1 domain sequences with an insertion of ‘CTDC ’domain, highlighted in a black box.

**Supplementary Fig. 7.** Predicted protein structure of the MHC class II heterodimer formed by *BS1_hap1_DAA2.1* (alpha chain) and *BS1_hap1_DAB2.1* (beta chain). **a** Model colored with the predicted local distance difference test values (pLDDT) of the alpha and beta chains. The predicted template modeling score (pTM) and interface predicted template modeling score (ipTM) are annotated. **b** Predicted aligned error (PAE) of the model. Amino acid residue number of alpha and beta chains are annotated by the axes of the plot.

**Supplementary Fig. 8.** Metrics of predicted MHC class II protein model *BS1_hap1_DAA2.1* (alpha chain) and *BS1_hap1_DAB2.1* (beta chain). **a, b** Results from SignalP 6.0 signal peptide prediction of the alpha chain and beta chain, respectively. **c, d** Results from DeepTMHMM transmembrane region prediction of the alpha chain and beta chain, respectively.

**Supplementary Table 1.** Non-synonymous substitutions (*dN*) and synonymous substitutions (*dS*) values computed by codon selection test in MEGA for exon2 and exon3 sequences of alpha and beta genes from DA and DB lineages along with their respective standard errors (S.E.).

**Supplementary Table 2.** Summary of studies on MHC II genes on teleost

**Supplementary Table 3**. Summary of null allele occurrences and mutations

**Supplementary Table 4.** Length (bp) of intron 1 in the *DAA1.3* gene from eight haplotypes

**Supplementary Table 5.** List of MHC class II α and β chain sequences from other species used in phylogenetic analyses, along with their corresponding accession numbers or gene IDs.

**Additional supplementary files (not included in the main Supplementary Information file) Supplementary Fig. 9.** Dotplots showing pairwise alignments of MHC class II haplotypes at Locus 2 from 28 phased haploid genome assemblies. Each page compares one assembly (y-axis) to all others (x-axes). The aligned sequence includes the full MHC class II region spanning the DA and DB lineage α-and β-chain genes, intergenic regions, and 20 kb of flanking sequence on each side. Red and blue lines indicate alignments in the same and opposite orientations, respectively. Gene positions are indicated by vertical and horizontal colored bars: green for *DAA*, blue for *DAB*, purple for *DBA*, and pink for *DBB* genes.

**Supplementary Fig. 10.** Dotplots showing pairwise alignments of MHC class II haplotypes at Locus 4 from 27 phased haploid genome assemblies. Each page compares one assembly (y-axis) to all others (x-axes). The aligned sequence includes the full MHC class II region spanning the DA and DB lineage α- and β-chain genes, intergenic regions, and 20 kb of flanking sequence on each side. Red and blue lines indicate alignments in the same and opposite orientations, respectively. Gene positions are indicated by vertical and horizontal colored bars: green for *DAA*, blue for *DAB*, purple for *DBA*, and pink for *DBB* genes.

**Supplementary File 1. Sites under positive selection identified by the M8 model in PAML.** Each row shows the codon position, corresponding amino acid, posterior probability of ω > 1 (Pr(w > 1)), and the posterior mean ± standard error (SE) of ω at that site. Sites with posterior probability ≥ 0.95 are marked with a single asterisk (*) and those with posterior probability ≥ 0.99 are marked with two asterisks (**), indicating high confidence of positive selection. Estimates are based on the Bayes Empirical Bayes (BEB) approach.

**Supplementary File 2**. Teleost MHC II amino acid sequences (*n* = 78) used as BLAST query to find MHC II sequences in Atlantic herring.

**Supplementary Video 1.** Animated visualization of the predicted quaternary protein structure of *CS4_hap1_DA2.1* modeled using AlphaFold3. The color gradient from red to white to blue represents normalized Shannon entropy values, indicating the degree of polymorphism from highest (red) to the lowest (blue).

