## Supplementary Text, Supplementary Figures 1-8, Supplementary tables for "Unheralded high MHC Class II polymorphism in the abundant Atlantic herring resolved by long-read sequencing"

**Non-random association of allelic groups in Locus 2 and 4 haplotypes**

In case of Locus 2, further careful observation about distribution of five major allelic groups (2a-e) in each of the 25 haplotypes revealed that haplotypes carry one of the two major sets of allelic groups. One set consisted of 2a, 2b, and 2e found in 15 haplotypes (2*1 – 2*9 in Fig 6b); while another set consists of 2c, 2d, and 2e found in the remaining 10 haplotypes (2*10 – 2*14 in Fig 6b). This indicated that allelic groups on each haplotype are not randomly arranged, and a set of these allelic group combinations could be a major MHC haplotype. By this convention, Locus 2 would contain two major haplotypes, one with allelic combination 2a|2b|2c and another with 2d|2e|2c. Since many of the 25 haplotypes carried only two gene pairs, it was difficult to say if the MHC haplotype structure is conserved irrespective of number of gene copies at a locus. To study this aspect, we used the non-genic sequence of each haplotype in addition to the genic sequence. We compared the sequences of an entire MHC II locus containing whole-length genes, intergenic sequences, and 20 kb flanking sequence among 29 haplotypes (Supplementary Fig. 9). This analysis categorized the same 15 and 10 haplotypes from the previous analysis under 2a|2b|2c and 2d|2e|2c, respectively (Fig. 6b). Furthermore, it categorized null alleles into allelic groups and interestingly, the null alleles also followed the expected pattern for functional genes (Fig. 6b). The most important observation was that in cases of haplotypes where only two gene pairs are present, we also observed absence of non-genic sequence corresponding to the third gene pair (Supplementary Fig. 9, represented by dashed lines in Fig. 6b).

In case of Locus 4, most of the haplotypes contained only single gene pair, of which 15 belonged to group 4a and 8 belonged to group 4b. Three haplotypes that contained two gene pairs had MHC haplotype structure as 4a|4b. Only one haplotype (NSSH10_hap2) with two gene pairs carried genes from the same allelic group 4a, deviating from the earlier observation from Locus 2 that each haplotype maintains high diversity by harboring different set of allelic groups. Interestingly, Locus 4 harbors inversions, which affects the order in which gene pairs are arranged in each haplotype (Extended Data Fig. 5b; Supplementary Fig. 10). Similar to Locus 2, the analysis with non-genic sequence of Locus 4, revealed that there are three major MHC haplotypes with allelic groups 4a, 4b, and 4a|4c (Extended Data Fig. 5b).

These major haplotypes (2a|2b|2c, 2d|2e|2c, 4a, 4b, and 4a|4c) resemble the conserved ancestral haploblocks (CEH) observed in humans, which are blocks of minimum recombination maintained over many generations and containing specific combinations of HLA and non-HLA alleles[1–3]. Polymorphism at sequence and copy number level for HLA and non-HLA alleles is observed in CEH[1–3].

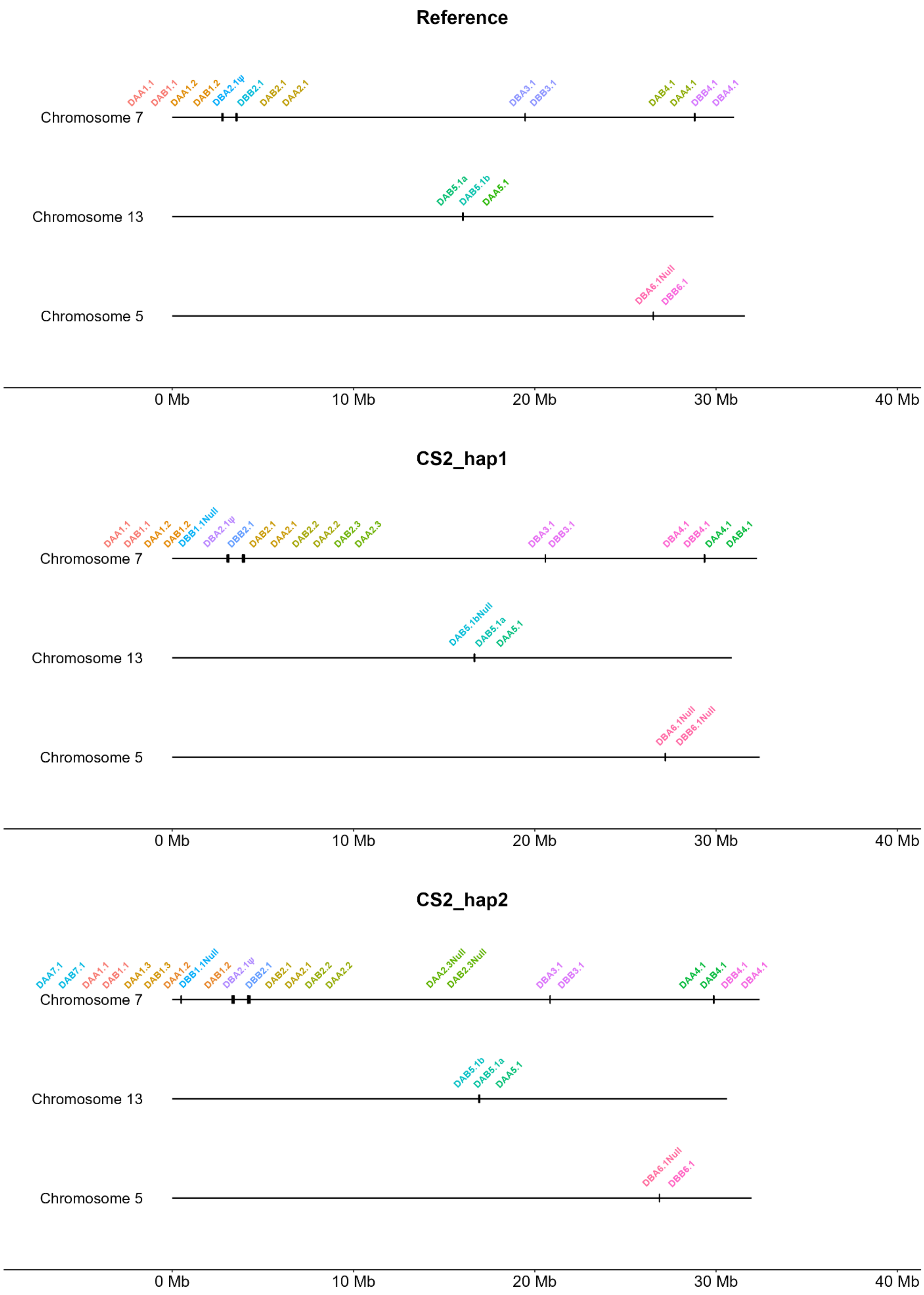
Reference
DAA1.1
DAB1.1
DAA1.2
DAB1.2
DBA2.1ψ
DBB2.1
DAB2.1
DAA2.1
DBA3.1
DBB3.1
DAB4.1
DAA4.1
DBB4.1
DBA4.1
Chromosome 7
DAB5.1a
DAB5.1b
DAA5.1
Chromosome 13
DBA6.1Null
DBB6.1
Chromosome 5
0 Mb
10 Mb
20 Mb
30 Mb
40 Mb
CS2_hap1
DAA1.1
DAB1.1
DAA1.2
DAB1.2
DBB1.1Null
DBA2.1ψ
DBB2.1
DAB2.1
DAA2.1
DAB2.2
DAA2.2
DAB2.3
DAA2.3
DBA3.1
DBB3.1
DBA4.1
DBB4.1
DAA4.1
DAB4.1
Chromosome 7
DAB5.1bNull
DAB5.1a
DAA5.1
Chromosome 13
DBA6.1Null
DBB6.1Null
Chromosome 5
0 Mb
10 Mb
20 Mb
30 Mb
40 Mb
CS2_hap2
DAA7.1
DAB7.1
DAA1.1
DAB1.1
DAA1.3
DAB1.3
DAA1.2
DBB1.1Null
DAB1.2
DBA2.1ψ
DBB2.1
DAB2.1
DAA2.1
DAB2.2
DAA2.2
DAA2.3Null
DAB2.3Null
DBA3.1
DBB3.1
DAA4.1
DAB4.1
DBB4.1
DBA4.1
Chromosome 7
DAB5.1b
DAB5.1a
DAA5.1
Chromosome 13
DBA6.1Null
DBB6.1
Chromosome 5
0 Mb
10 Mb
20 Mb
30 Mb
40 Mb

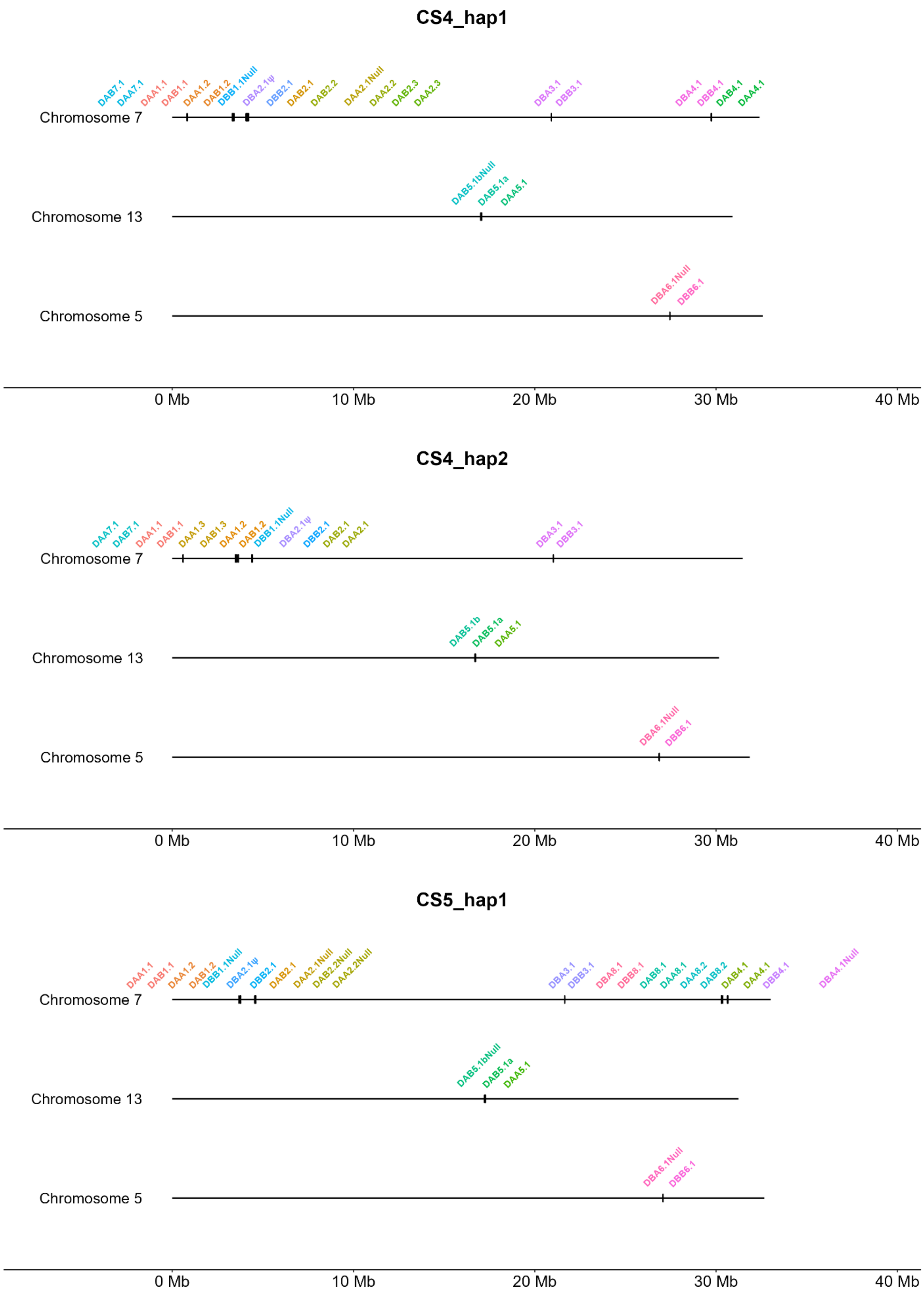
CS4_hap1
DAB7.1
DAA7.1
DAA1.1
DAB1.1
DAA1.2
DAB1.2
DBB1.1Null
DBA2.1ψ
DBB2.1
DAB2.1
DAB2.2
DAA2.1Null
DAA2.2
DAB2.3
DAA2.3
DBA3.1
DBB3.1
DBA4.1
DBB4.1
DAB4.1
DAA4.1
Chromosome 7
DAB5.1bNull
DAB5.1a
DAA5.1
Chromosome 13
DBA6.1Null
DBB6.1
Chromosome 5
0 Mb
10 Mb
20 Mb
30 Mb
40 Mb
CS4_hap2
DAA7.1
DAB7.1
DAA1.1
DAB1.1
DAA1.3
DAB1.3
DAA1.2
DAB1.2
DBB1.1Null
DBA2.1ψ
DBB2.1
DAB2.1
DAA2.1
DBA3.1
DBB3.1
Chromosome 7
DAB5.1b
DAB5.1a
DAA5.1
Chromosome 13
DBA6.1Null
DBB6.1
Chromosome 5
0 Mb
10 Mb
20 Mb
30 Mb
40 Mb
CS5_hap1
DAA1.1
DAB1.1
DAA1.2
DAB1.2
DBB1.1Null
DBA2.1ψ
DBB2.1
DAB2.1
DAA2.1Null
DAB2.2Null
DAA2.2Null
DBA3.1
DBB3.1
DBA8.1
DBB8.1
DAB8.1
DAA8.1
DAA8.2
DAB8.2
DAB4.1
DAA4.1
DBB4.1
DBA4.1Null
Chromosome 7
DAB5.1bNull
DAB5.1a
DAA5.1
Chromosome 13
DBA6.1Null
DBB6.1
Chromosome 5
0 Mb
10 Mb
20 Mb
30 Mb
40 Mb

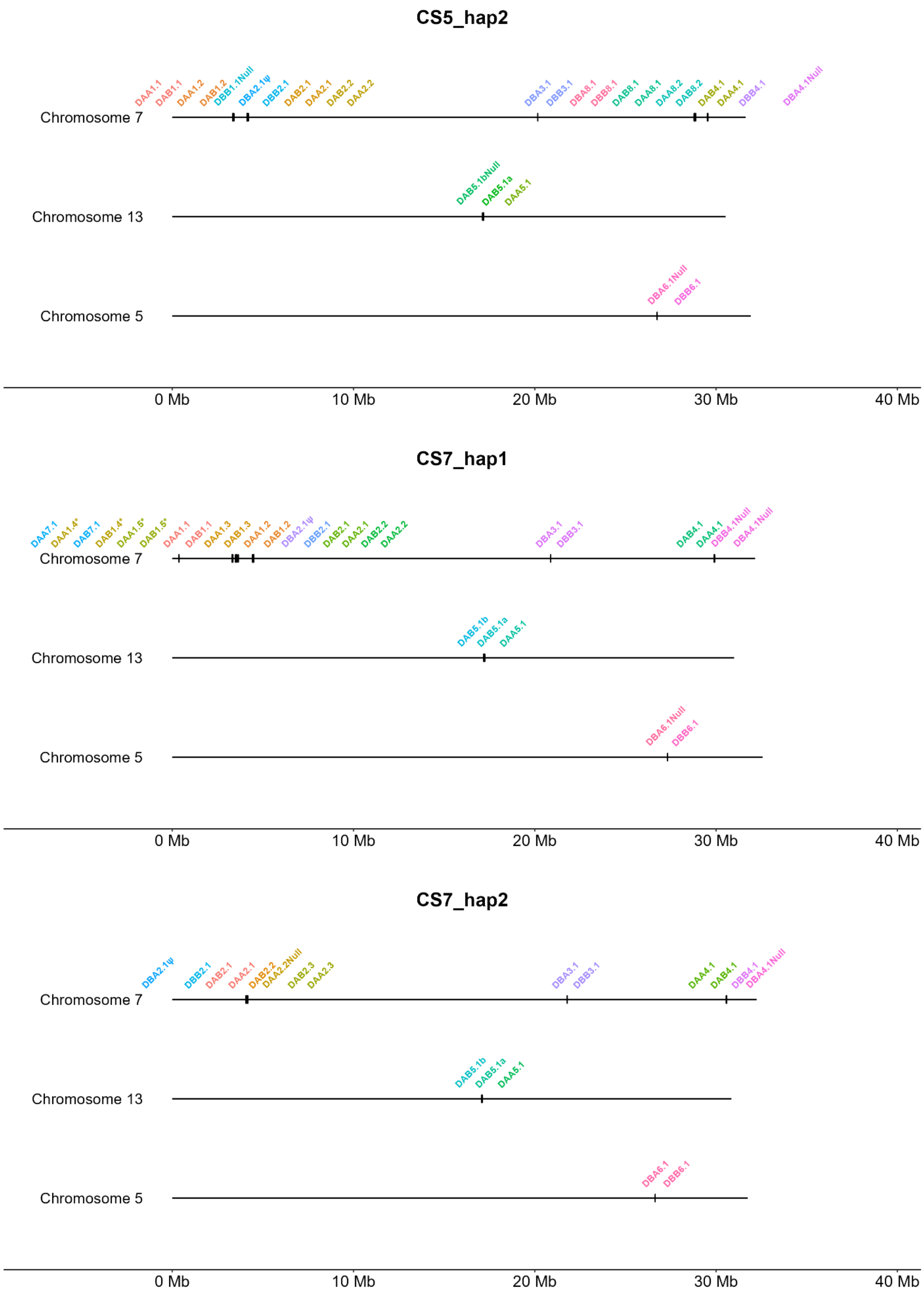
CS5_hap2
DAA1.1
DAB1.1
DAA1.2
DAB1.2
DBB1.1Null
DBA2.1ψ
DBB2.1
DAB2.1
DAA2.1
DAB2.2
DAA2.2
DBA3.1
DBB3.1
DBA8.1
DBB8.1
DAB8.1
DAA8.1
DAA8.2
DAB8.2
DAB4.1
DAA4.1
DBB4.1
DBA4.1Null
Chromosome 7
DAB5.1bNull
DAB5.1a
DAA5.1
Chromosome 13
DBA6.1Null
DBB6.1
Chromosome 5
0 Mb
10 Mb
20 Mb
30 Mb
40 Mb
CS7_hap1
DAA7.1
DAA1.4*
DAB7.1
DAB1.4*
DAA1.5*
DAB1.5*
DAA1.1
DAB1.1
DAA1.3
DAB1.3
DAA1.2
DAB1.2
DBA2.1ψ
DBB2.1
DAB2.1
DAA2.1
DAB2.2
DAA2.2
DBA3.1
DBB3.1
DAB4.1
DAA4.1
DBB4.1Null
DBA4.1Null
Chromosome 7
DAB5.1b
DAB5.1a
DAA5.1
Chromosome 13
DBA6.1Null
DBB6.1
Chromosome 5
0 Mb
10 Mb
20 Mb
30 Mb
40 Mb
CS7_hap2
DBA2.1ψ
DBB2.1
DAB2.1
DAA2.1
DAB2.2
DAA2.2Null
DAB2.3
DAA2.3
DBA3.1
DBB3.1
DAA4.1
DAB4.1
DBB4.1
DBA4.1Null
Chromosome 7
DAB5.1b
DAB5.1a
DAA5.1
Chromosome 13
DBA6.1
DBB6.1
Chromosome 5
0 Mb
10 Mb
20 Mb
30 Mb
40 Mb

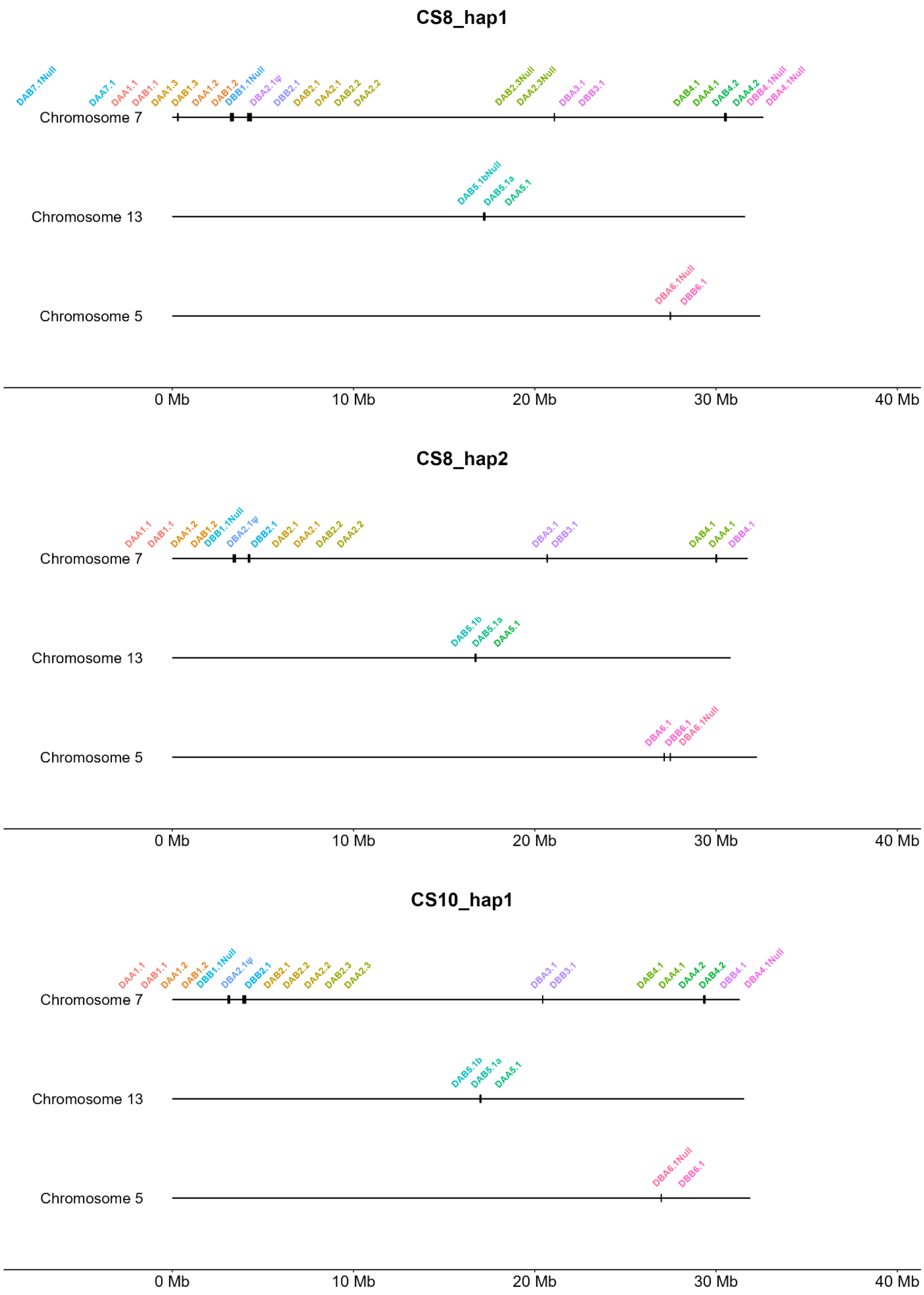
CS8_hap1
DAB7.1Null
DAA7.1
DAA1.1
DAB1.1
DAA1.3
DAB1.3
DAA1.2
DAB1.2
DBB1.1Null
DBA2.1ψ
DBB2.1
DAB2.1
DAA2.1
DAB2.2
DAA2.2
DAB2.3Null
DAA2.3Null
DBA3.1
DBB3.1
DAB4.1
DAA4.1
DAB4.2
DAA4.2
DBB4.1Null
DBA4.1Null
Chromosome 7
DAB5.1bNull
DAB5.1a
DAA5.1
Chromosome 13
DBA6.1Null
DBB6.1
Chromosome 5
0 Mb
10 Mb
20 Mb
30 Mb
40 Mb
CS8_hap2
DAA1.1
DAB1.1
DAA1.2
DAB1.2
DBB1.1Null
DBA2.1ψ
DBB2.1
DAB2.1
DAA2.1
DAB2.2
DAA2.2
DBA3.1
DBB3.1
DAB4.1
DAA4.1
DBB4.1
Chromosome 7
DAB5.1b
DAB5.1a
DAA5.1
Chromosome 13
DBA6.1
DBB6.1
DBA6.1Null
Chromosome 5
0 Mb
10 Mb
20 Mb
30 Mb
40 Mb
CS10_hap1
DAA1.1
DAB1.1
DAA1.2
DAB1.2
DBB1.1Null
DBA2.1ψ
DBB2.1
DAB2.1
DAB2.2
DAA2.2
DAB2.3
DAA2.3
DBA3.1
DBB3.1
DAB4.1
DAA4.1
DAA4.2
DAB4.2
DBB4.1
DBA4.1Null
Chromosome 7
DAB5.1b
DAB5.1a
DAA5.1
Chromosome 13
DBA6.1Null
DBB6.1
Chromosome 5
0 Mb
10 Mb
20 Mb
30 Mb
40 Mb

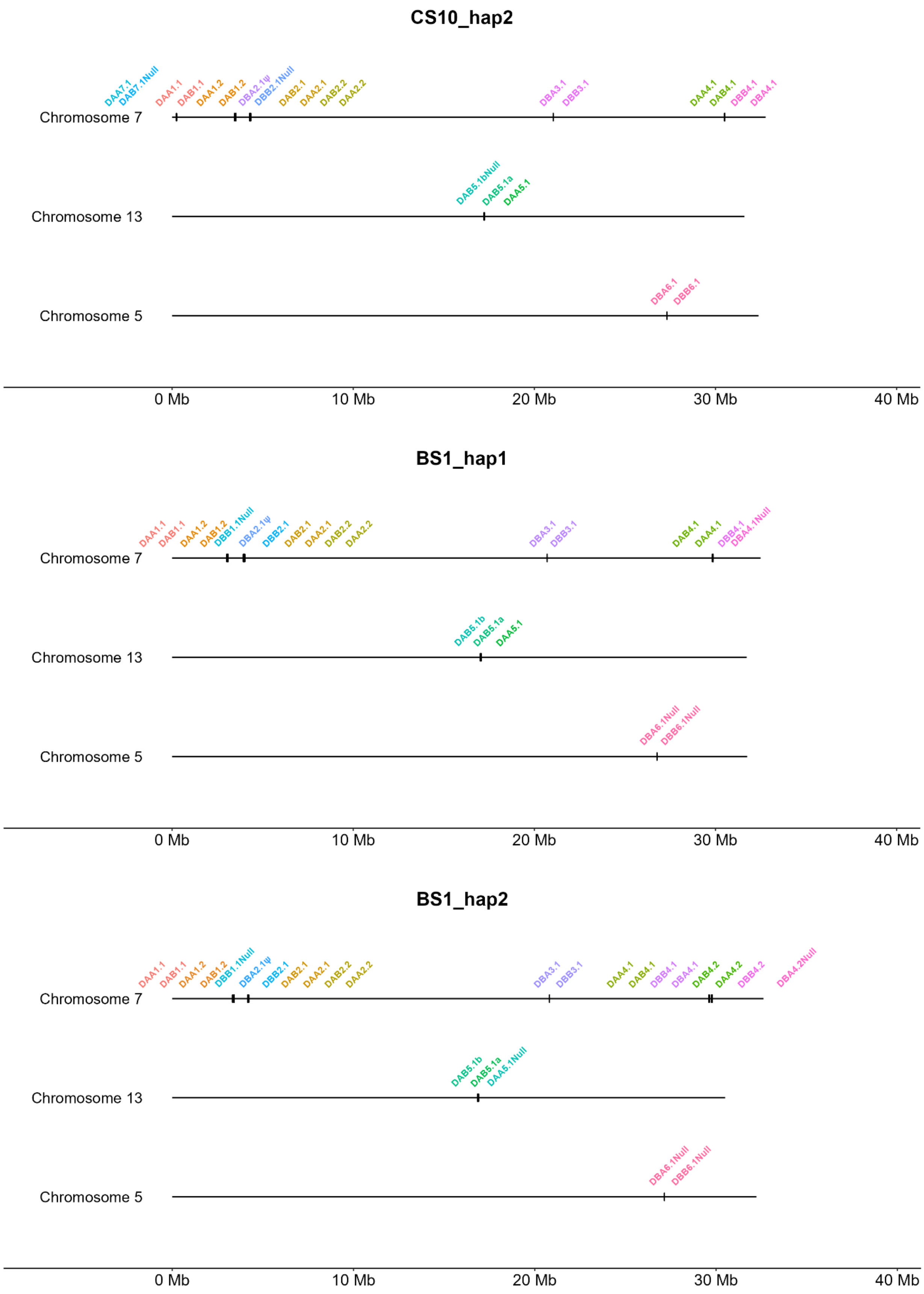

CS10_hap2
DAA7.1
DAB7.1Null
DAA1.1
DAB1.1
DAA1.2
DAB1.2
DBA2.1ψ
DBB2.1Null
DAB2.1
DAA2.1
DAB2.2
DAA2.2
DBA3.1
DBB3.1
DAA4.1
DAB4.1
DBB4.1
DBA4.1
Chromosome 7
DAB5.1bNull
DAB5.1a
DAA5.1
Chromosome 13
DBA6.1
DBB6.1
Chromosome 5
0 Mb
10 Mb
20 Mb
30 Mb
40 Mb
BS1_hap1
DAA1.1
DAB1.1
DAA1.2
DAB1.2
DBB1.1Null
DBA2.1ψ
DBB2.1
DAB2.1
DAA2.1
DAB2.2
DAA2.2
DBA3.1
DBB3.1
DAB4.1
DAA4.1
DBB4.1
DBA4.1Null
Chromosome 7
DAB5.1b
DAB5.1a
DAA5.1
Chromosome 13
DBA6.1Null
DBB6.1Null
Chromosome 5
0 Mb
10 Mb
20 Mb
30 Mb
40 Mb
BS1_hap2
DAA1.1
DAB1.1
DAA1.2
DAB1.2
DBB1.1Null
DBA2.1ψ
DBB2.1
DAB2.1
DAA2.1
DAB2.2
DAA2.2
DBA3.1
DBB3.1
DAA4.1
DAB4.1
DBB4.1
DBA4.1
DAB4.2
DAA4.2
DBB4.2
DBA4.2Null
Chromosome 7
DAB5.1b
DAB5.1a
DAA5.1Null
Chromosome 13
DBA6.1Null
DBB6.1Null
Chromosome 5
0 Mb
10 Mb
20 Mb
30 Mb
40 Mb

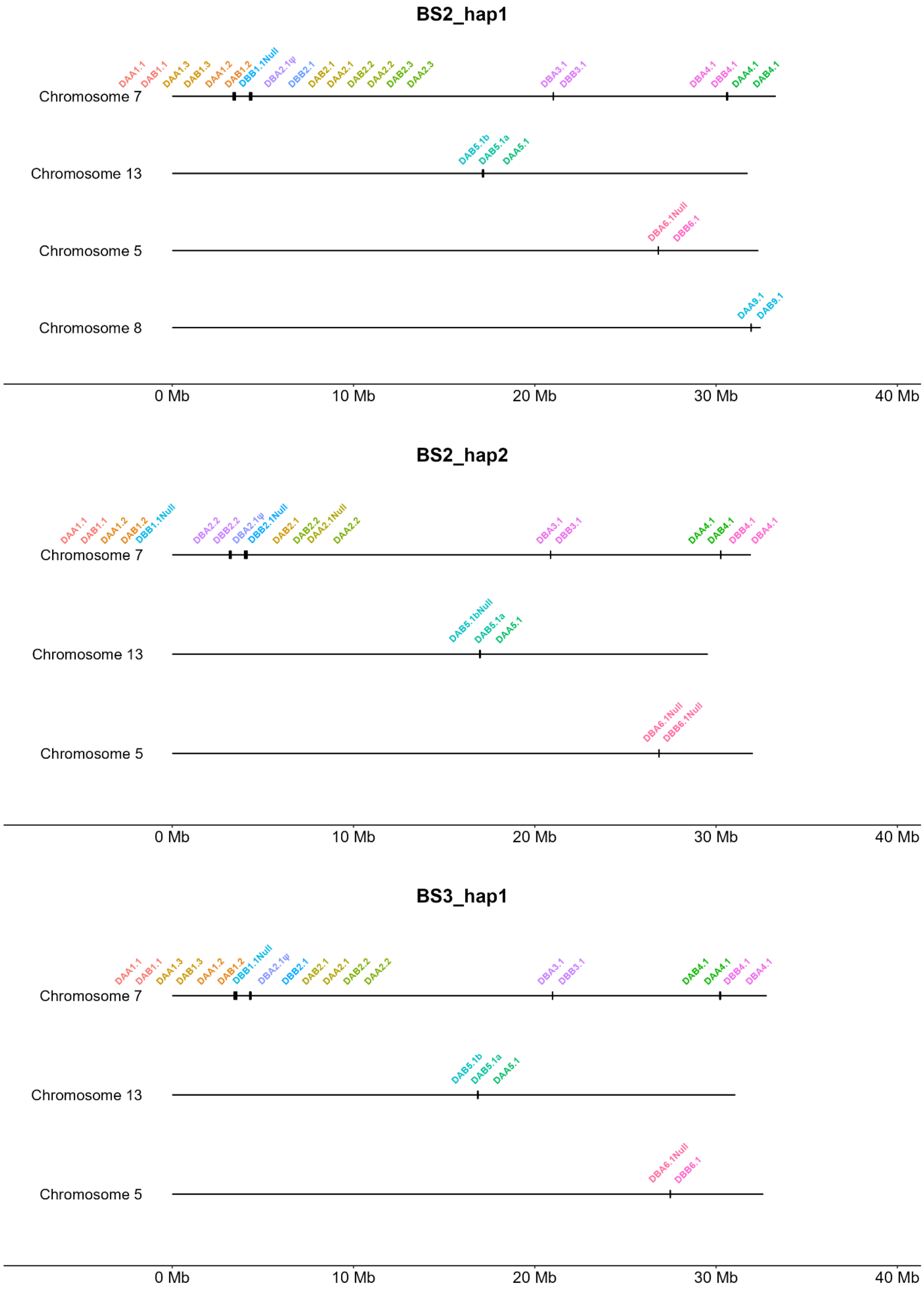

BS2_hap1
DAA1.1
DAB1.1
DAA1.3
DAB1.3
DAA1.2
DAB1.2
DBB1.1Null
DBA2.1ψ
DBB2.1
DAB2.1
DAA2.1
DAB2.2
DAA2.2
DAB2.3
DAA2.3
DBA3.1
DBB3.1
DBA4.1
DBB4.1
DAA4.1
DAB4.1
Chromosome 7
DAB5.1b
DAB5.1a
DAA5.1
Chromosome 13
DBA6.1Null
DBB6.1
Chromosome 5
DAA9.1
DAB9.1
Chromosome 8
0 Mb
10 Mb
20 Mb
30 Mb
40 Mb
BS2_hap2
DAA1.1
DAB1.1
DAA1.2
DAB1.2
DBB1.1Null
DBA2.2
DBB2.2
DBA2.1ψ
DBB2.1Null
DAB2.1
DAB2.2
DAA2.1Null
DAA2.2
DBA3.1
DBB3.1
DAA4.1
DAB4.1
DBB4.1
DBA4.1
Chromosome 7
DAB5.1bNull
DAB5.1a
DAA5.1
Chromosome 13
DBA6.1Null
DBB6.1Null
Chromosome 5
0 Mb
10 Mb
20 Mb
30 Mb
40 Mb
BS3_hap1
DAA1.1
DAB1.1
DAA1.3
DAB1.3
DAA1.2
DAB1.2
DBB1.1Null
DBA2.1ψ
DBB2.1
DAB2.1
DAA2.1
DAB2.2
DAA2.2
DBA3.1
DBB3.1
DAB4.1
DAA4.1
DBB4.1
DBA4.1
Chromosome 7
DAB5.1b
DAB5.1a
DAA5.1
Chromosome 13
DBA6.1Null
DBB6.1
Chromosome 5
0 Mb
10 Mb
20 Mb
30 Mb
40 Mb

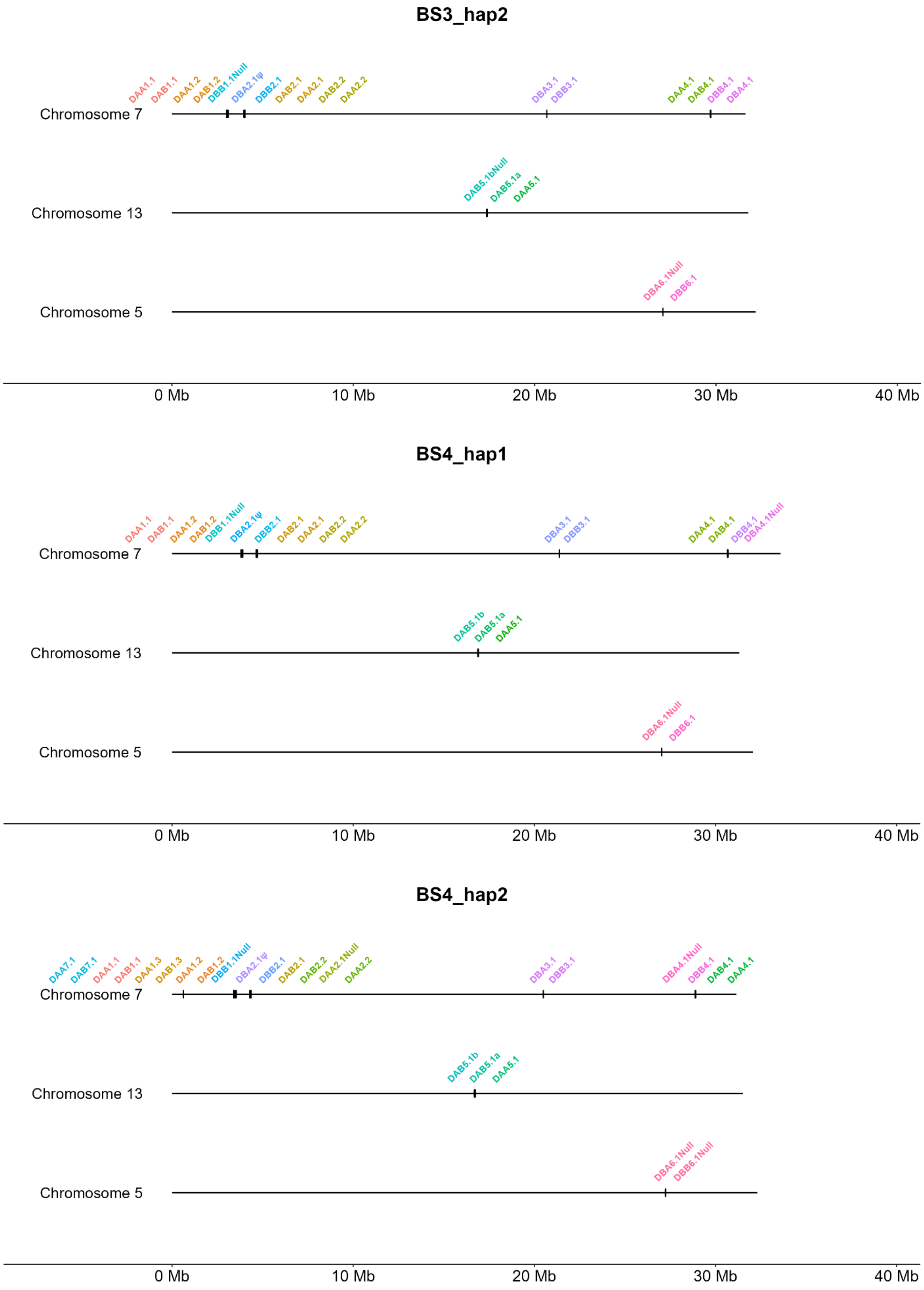
BS3_hap2
DAA1.1
DAB1.1
DAA1.2
DAB1.2
DBB1.1Null
DBA2.1ψ
DBB2.1
DAB2.1
DAA2.1
DAB2.2
DAA2.2
DBA3.1
DBB3.1
DAA4.1
DAB4.1
DBB4.1
DBA4.1
Chromosome 7
DAB5.1bNull
DAB5.1a
DAA5.1
Chromosome 13
DBA6.1Null
DBB6.1
Chromosome 5
0 Mb
10 Mb
20 Mb
30 Mb
40 Mb
BS4_hap1
DAA1.1
DAB1.1
DAA1.2
DAB1.2
DBB1.1Null
DBA2.1ψ
DBB2.1
DAB2.1
DAA2.1
DAB2.2
DAA2.2
DBA3.1
DBB3.1
DAA4.1
DAB4.1
DBB4.1
DBA4.1Null
Chromosome 7
DAB5.1b
DAB5.1a
DAA5.1
Chromosome 13
DBA6.1Null
DBB6.1
Chromosome 5
0 Mb
10 Mb
20 Mb
30 Mb
40 Mb
BS4_hap2
DAA7.1
DAB7.1
DAA1.1
DAB1.1
DAA1.3
DAB1.3
DAA1.2
DAB1.2
DBB1.1Null
DBA2.1ψ
DBB2.1
DAB2.1
DAB2.2
DAA2.1Null
DAA2.2
DBA3.1
DBB3.1
DBA4.1Null
DBB4.1
DAB4.1
DAA4.1
Chromosome 7
DAB5.1b
DAB5.1a
DAA5.1
Chromosome 13
DBA6.1Null
DBB6.1Null
Chromosome 5
0 Mb
10 Mb
20 Mb
30 Mb
40 Mb

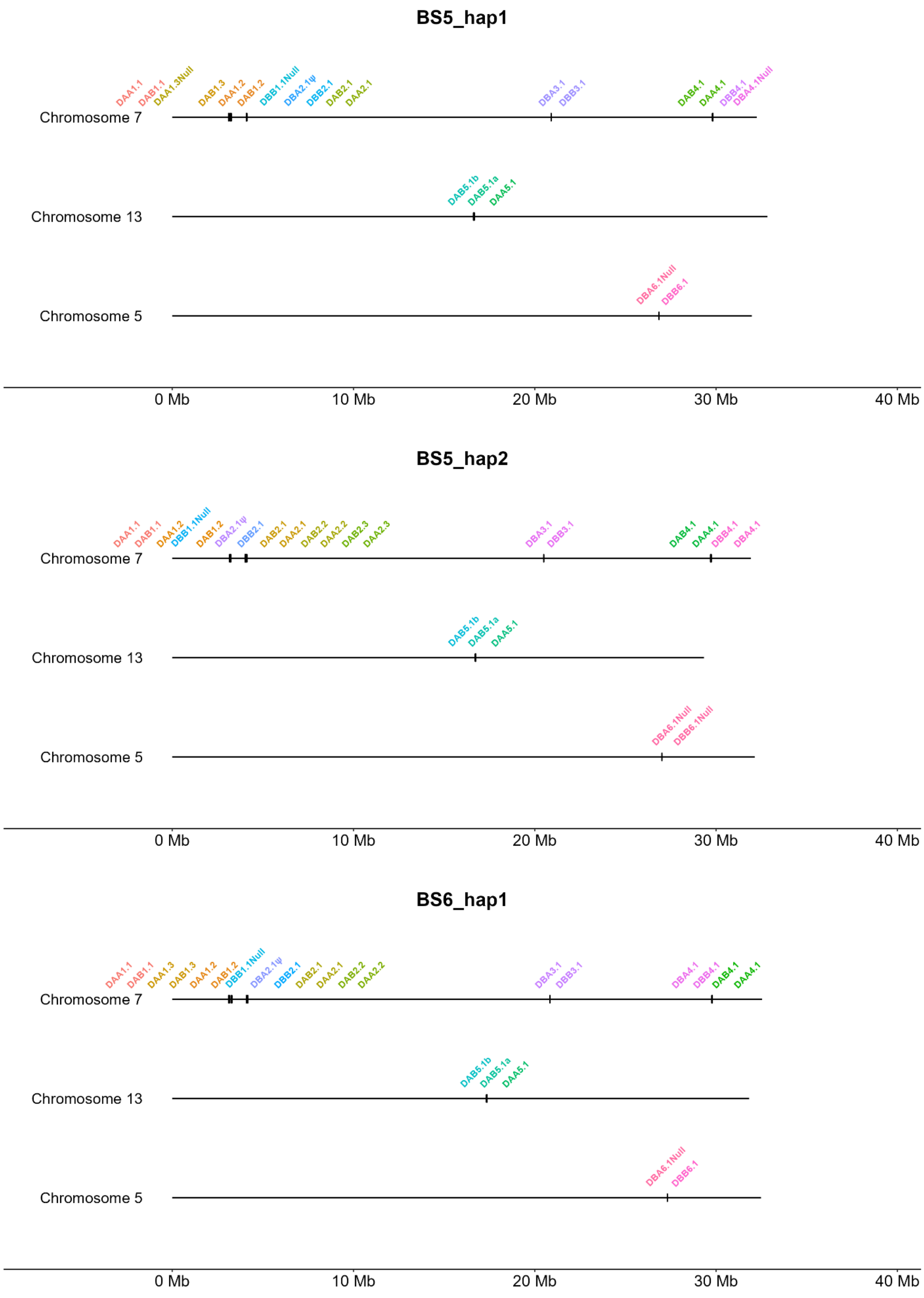
BS5_hap1
DAA1.1
DAB1.1
DAA1.3Null
DAB1.3
DAA1.2
DAB1.2
DBB1.1Null
DBA2.1ψ
DBB2.1
DAB2.1
DAA2.1
DBA3.1
DBB3.1
DAB4.1
DAA4.1
DBB4.1
DBA4.1Null
Chromosome 7
DAB5.1b
DAB5.1a
DAA5.1
Chromosome 13
DBA6.1Null
DBB6.1
Chromosome 5
0 Mb
10 Mb
20 Mb
30 Mb
40 Mb
BS5_hap2
DAA1.1
DAB1.1
DAA1.2
DBB1.1Null
DAB1.2
DBA2.1ψ
DBB2.1
DAB2.1
DAA2.1
DAB2.2
DAA2.2
DAB2.3
DAA2.3
DBA3.1
DBB3.1
DAB4.1
DAA4.1
DBB4.1
DBA4.1
Chromosome 7
DAB5.1b
DAB5.1a
DAA5.1
Chromosome 13
DBA6.1Null
DBB6.1Null
Chromosome 5
0 Mb
10 Mb
20 Mb
30 Mb
40 Mb
BS6_hap1
DAA1.1
DAB1.1
DAA1.3
DAB1.3
DAA1.2
DAB1.2
DBB1.1Null
DBA2.1ψ
DBB2.1
DAB2.1
DAA2.1
DAB2.2
DAA2.2
DBA3.1
DBB3.1
DBA4.1
DBB4.1
DAB4.1
DAA4.1
Chromosome 7
DAB5.1b
DAB5.1a
DAA5.1
Chromosome 13
DBA6.1Null
DBB6.1
Chromosome 5
0 Mb
10 Mb
20 Mb
30 Mb
40 Mb

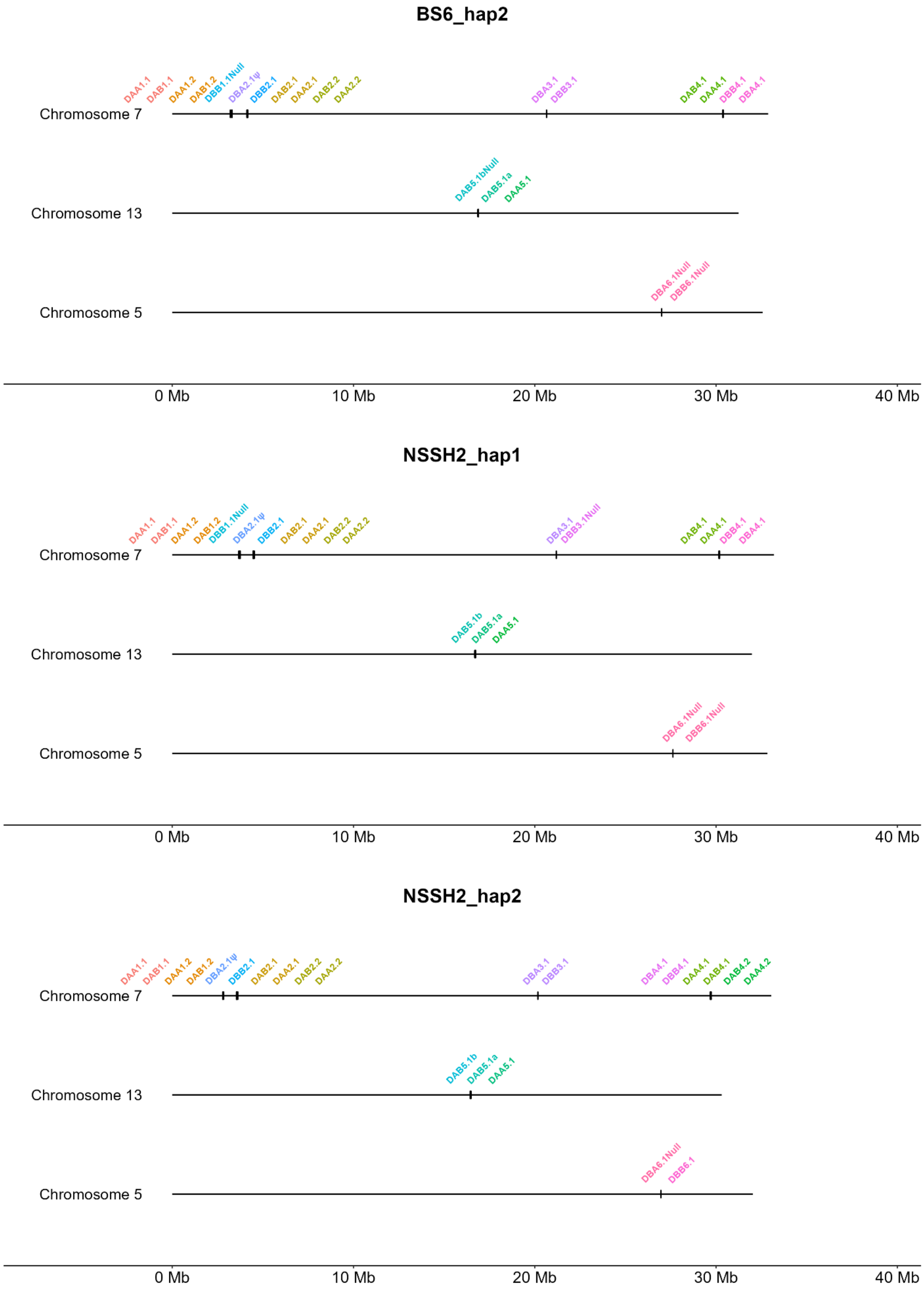
BS6_hap2
DAA1.1
DAB1.1
DAA1.2
DAB1.2
DBB1.1Null
DBA2.1ψ
DBB2.1
DAB2.1
DAA2.1
DAB2.2
DAA2.2
DBA3.1
DBB3.1
DAB4.1
DAA4.1
DBB4.1
DBA4.1
Chromosome 7
DAB5.1bNull
DAB5.1a
DAA5.1
Chromosome 13
DBA6.1Null
DBB6.1Null
Chromosome 5
0 Mb
10 Mb
20 Mb
30 Mb
40 Mb
NSSH2_hap1
DAA1.1
DAB1.1
DAA1.2
DAB1.2
DBB1.1Null
DBA2.1ψ
DBB2.1
DAB2.1
DAA2.1
DAB2.2
DAA2.2
DBA3.1
DBB3.1Null
DAB4.1
DAA4.1
DBB4.1
DBA4.1
Chromosome 7
DAB5.1b
DAB5.1a
DAA5.1
Chromosome 13
DBA6.1Null
DBB6.1Null
Chromosome 5
0 Mb
10 Mb
20 Mb
30 Mb
40 Mb
NSSH2_hap2
DAA1.1
DAB1.1
DAA1.2
DAB1.2
DBA2.1ψ
DBB2.1
DAB2.1
DAA2.1
DAB2.2
DAA2.2
DBA3.1
DBB3.1
DBA4.1
DBB4.1
DAA4.1
DAB4.1
DAB4.2
DAA4.2
Chromosome 7
DAB5.1b
DAB5.1a
DAA5.1
Chromosome 13
DBA6.1Null
DBB6.1
Chromosome 5
0 Mb
10 Mb
20 Mb
30 Mb
40 Mb

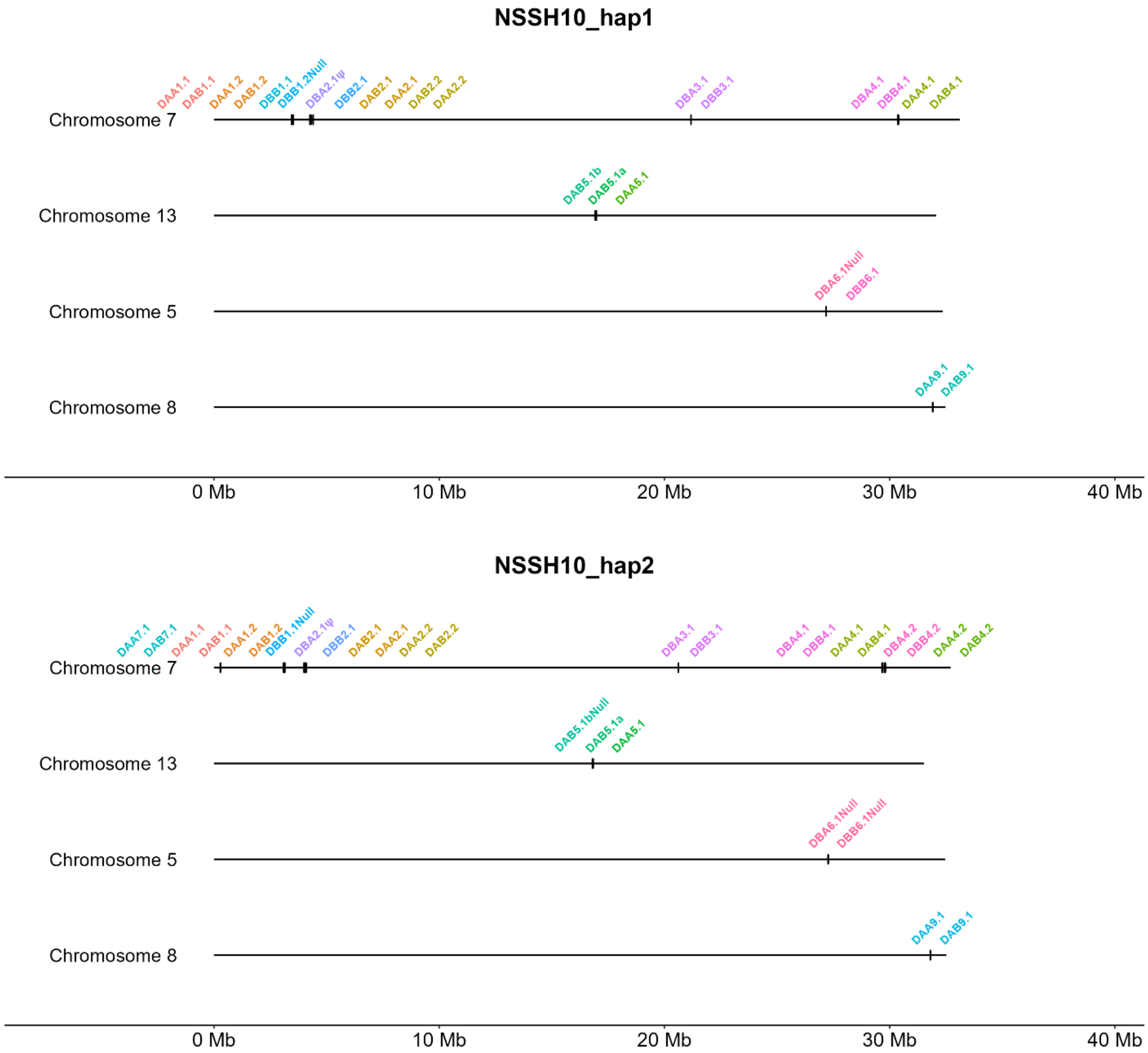


**Supplementary Fig. 1.** Genome organization of MHC II genes on 29 haploid assemblies. Alpha-beta gene pairs are colored based on their respective locus. **DA1.4* and *DA1.5* genes on CS7_hap1 are misassembled and most likely present on CS7_hap2 as *DA1.1* and *DA1.2*, respectively (detail explanation in methods).

### Locus1

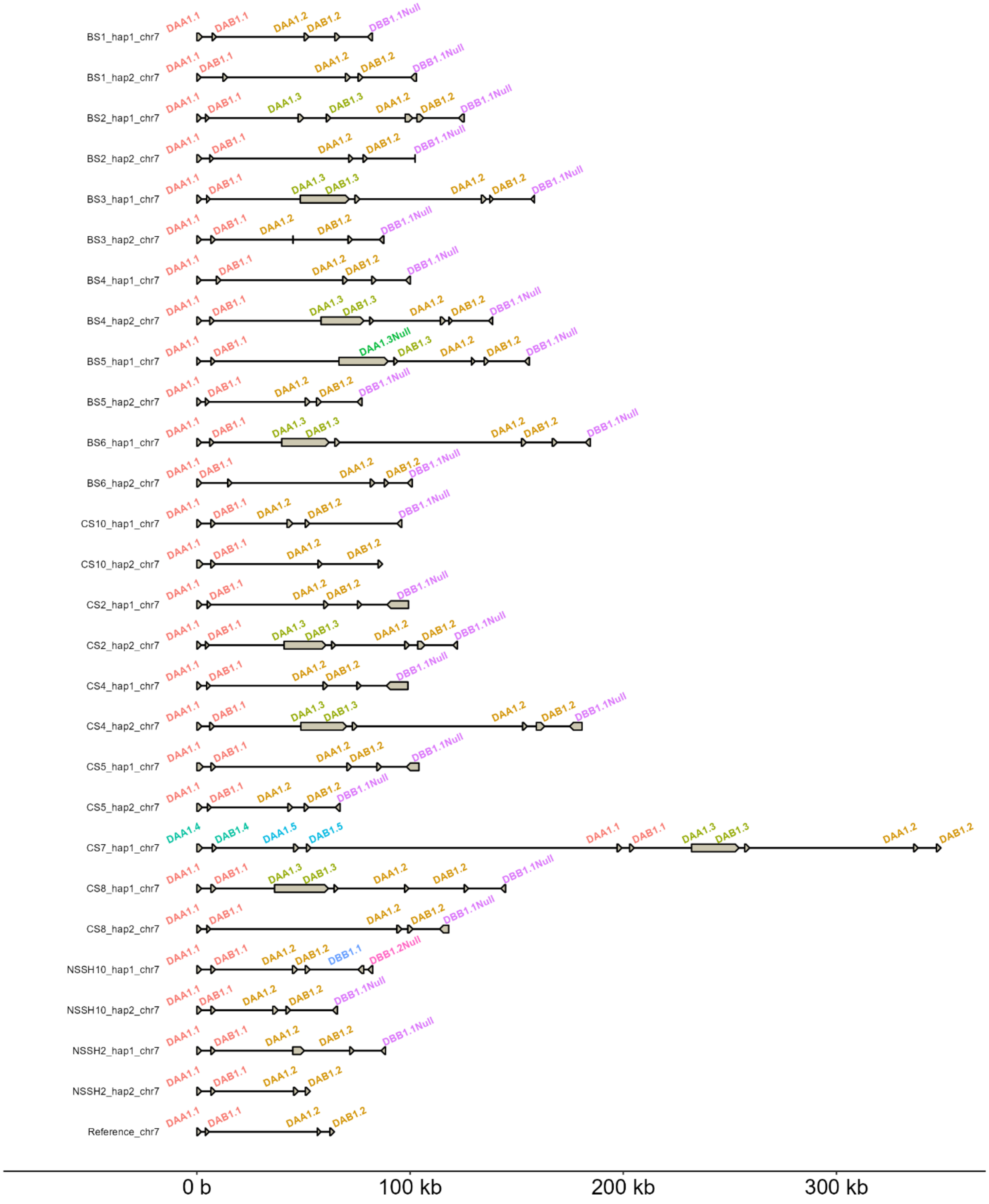

BS1_hap1_chr7 DAA1.1 DAB1.1 DAA1.2 DAB1.2 DBB1.1Null
BS1_hap2_chr7 DAA1.1 DAB1.1 DAA1.2 DAB1.2 DBB1.1Null
BS2_hap1_chr7 DAA1.1 DAB1.1 DAA1.3 DAB1.3 DAA1.2 DAB1.2 DBB1.1Null
BS2_hap2_chr7 DAA1.1 DAB1.1 DAA1.2 DAB1.2 DBB1.1Null
BS3_hap1_chr7 DAA1.1 DAB1.1 DAA1.3 DAB1.3 DAA1.2 DAB1.2 DBB1.1Null
BS3_hap2_chr7 DAA1.1 DAB1.1 DAA1.2 DAB1.2 DBB1.1Null
BS4_hap1_chr7 DAA1.1 DAB1.1 DAA1.2 DAB1.2 DBB1.1Null
BS4_hap2_chr7 DAA1.1 DAB1.1 DAA1.3 DAB1.3 DAA1.2 DAB1.2 DBB1.1Null
BS5_hap1_chr7 DAA1.1 DAB1.1 DAA1.3Null DAB1.3 DAA1.2 DAB1.2 DBB1.1Null
BS5_hap2_chr7 DAA1.1 DAB1.1 DAA1.2 DAB1.2 DBB1.1Null
BS6_hap1_chr7 DAA1.1 DAB1.1 DAA1.3 DAB1.3 DAA1.2 DAB1.2 DBB1.1Null
BS6_hap2_chr7 DAA1.1 DAB1.1 DAA1.2 DAB1.2 DBB1.1Null
CS10_hap1_chr7 DAA1.1 DAB1.1 DAA1.2 DAB1.2 DBB1.1Null
CS10_hap2_chr7 DAA1.1 DAB1.1 DAA1.2 DAB1.2
CS2_hap1_chr7 DAA1.1 DAB1.1 DAA1.2 DAB1.2 DBB1.1Null
CS2_hap2_chr7 DAA1.1 DAB1.1 DAA1.3 DAB1.3 DAA1.2 DAB1.2 DBB1.1Null
CS4_hap1_chr7 DAA1.1 DAB1.1 DAA1.2 DAB1.2 DBB1.1Null
CS4_hap2_chr7 DAA1.1 DAB1.1 DAA1.3 DAB1.3 DAA1.2 DAB1.2 DBB1.1Null
CS5_hap1_chr7 DAA1.1 DAB1.1 DAA1.2 DAB1.2 DBB1.1Null
CS5_hap2_chr7 DAA1.1 DAB1.1 DAA1.2 DAB1.2 DBB1.1Null
CS7_hap1_chr7 DAA1.4 DAB1.4 DAA1.5 DAB1.5 DAA1.1 DAB1.1 DAA1.3 DAB1.3 DAA1.2 DAB1.2
CS8_hap1_chr7 DAA1.1 DAB1.1 DAA1.3 DAB1.3 DAA1.2 DAB1.2 DBB1.1Null
CS8_hap2_chr7 DAA1.1 DAB1.1 DAA1.2 DAB1.2 DBB1.1Null
NSSH10_hap1_chr7 DAA1.1 DAB1.1 DAA1.2 DAB1.2 DBB1.1 DBB1.2Null
NSSH10_hap2_chr7 DAA1.1 DAB1.1 DAA1.2 DAB1.2 DBB1.1Null
NSSH2_hap1_chr7 DAA1.1 DAB1.1 DAA1.2 DAB1.2 DBB1.1Null
NSSH2_hap2_chr7 DAA1.1 DAB1.1 DAA1.2 DAB1.2
Reference_chr7 DAA1.1 DAB1.1 DAA1.2 DAB1.2
0 b
100 kb
200 kb
300 kb

### Locus2

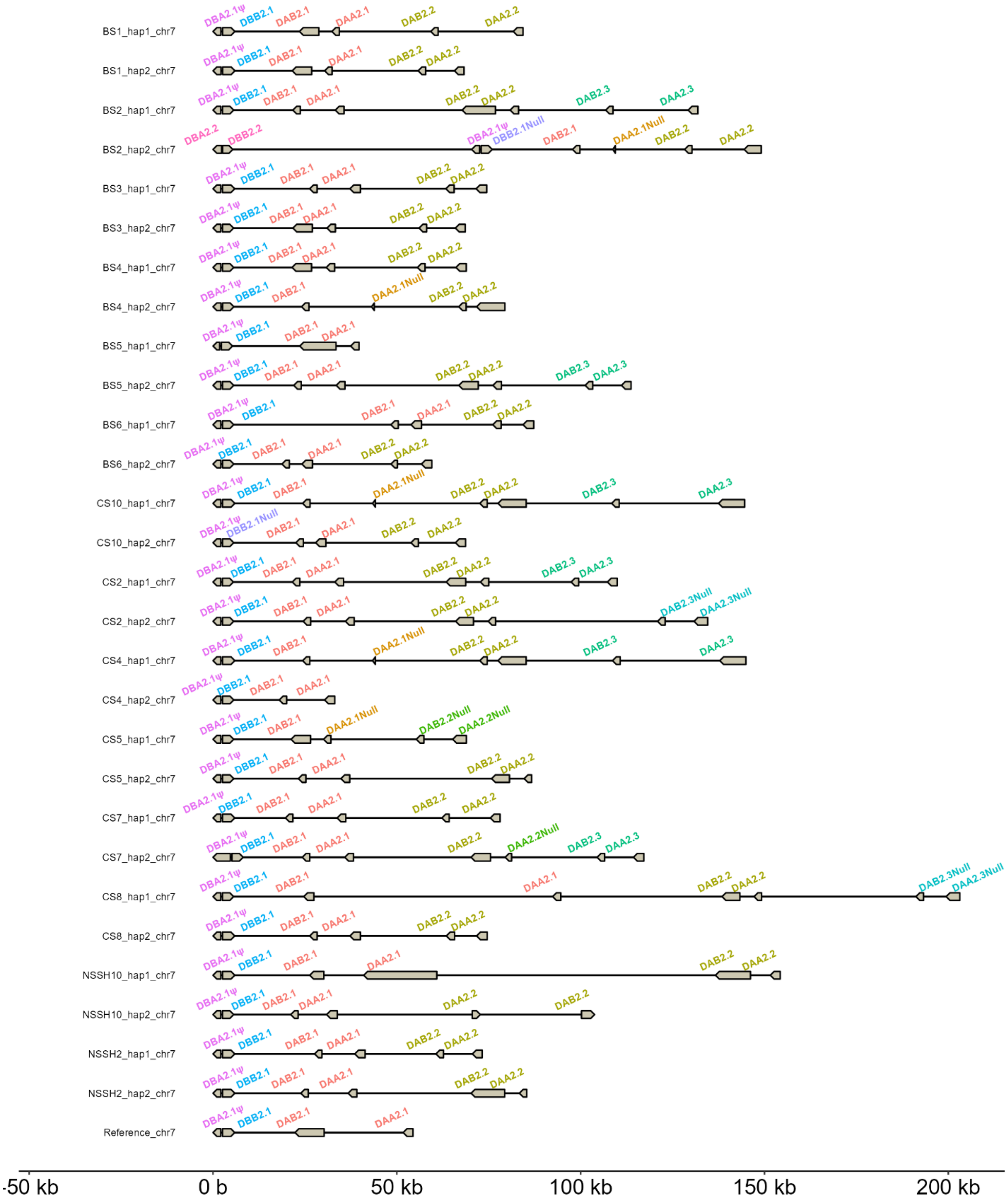

BS1_hap1_chr7
BS1_hap2_chr7
BS2_hap1_chr7
BS2_hap2_chr7
BS3_hap1_chr7
BS3_hap2_chr7
BS4_hap1_chr7
BS4_hap2_chr7
BS5_hap1_chr7
BS5_hap2_chr7
BS6_hap1_chr7
BS6_hap2_chr7
CS10_hap1_chr7
CS10_hap2_chr7
CS2_hap1_chr7
CS2_hap2_chr7
CS4_hap1_chr7
CS4_hap2_chr7
CS5_hap1_chr7
CS5_hap2_chr7
CS7_hap1_chr7
CS7_hap2_chr7
CS8_hap1_chr7
CS8_hap2_chr7
NSSH10_hap1_chr7
NSSH10_hap2_chr7
NSSH2_hap1_chr7
NSSH2_hap2_chr7
Reference_chr7
DBA2.1ψ
DBB2.1
DAB2.1
DAA2.1
DAB2.2
DAA2.2
DAB2.3
DAA2.3
DBA2.2
DBB2.2
DBB2.1Null
DAA2.1Null
DAB2.2Null
DAA2.2Null
DAB2.3Null
DAA2.3Null
-50 kb
0 b
50 kb
100 kb
150 kb
200 kb

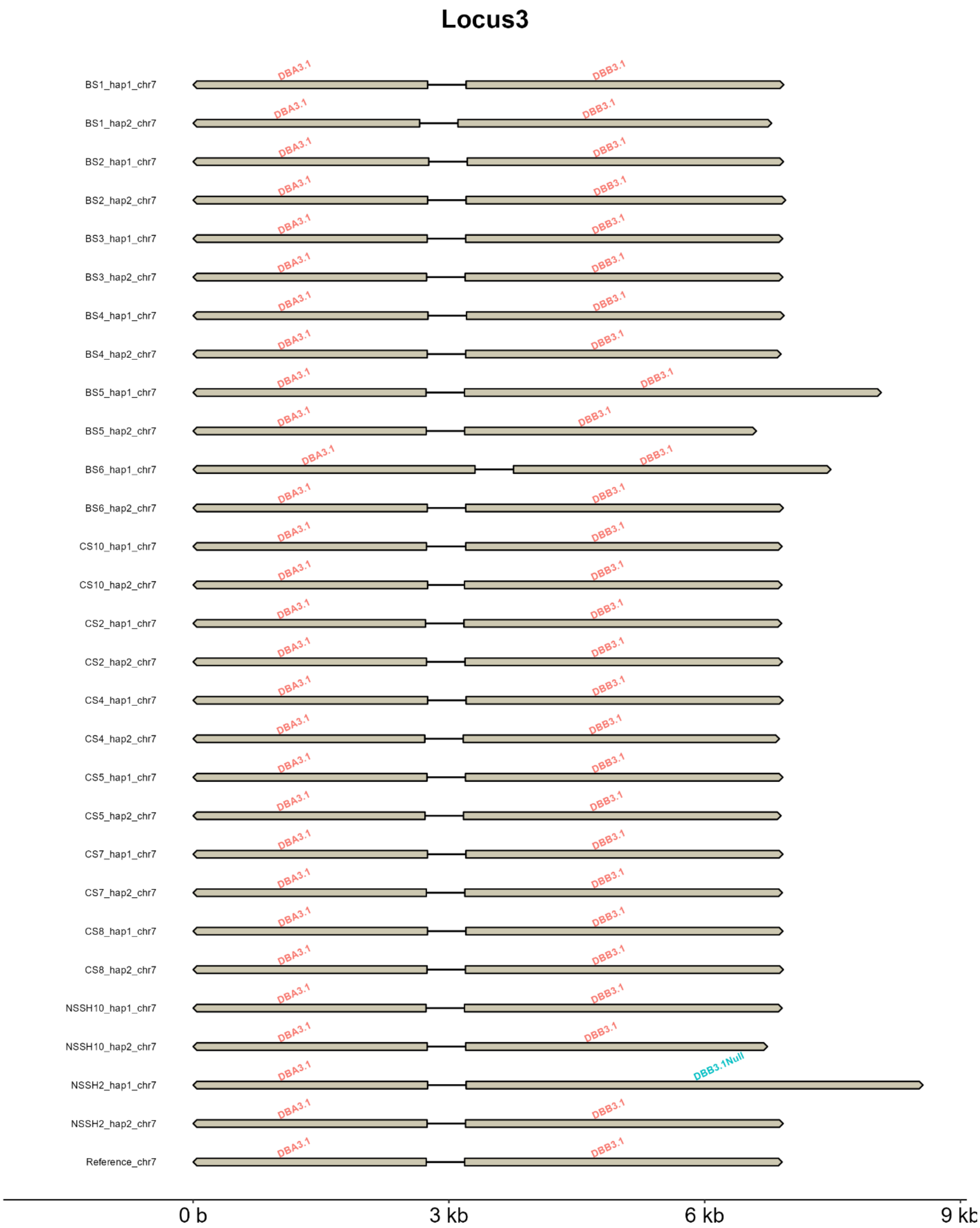
Locus3
BS1_hap1_chr7
BS1_hap2_chr7
BS2_hap1_chr7
BS2_hap2_chr7
BS3_hap1_chr7
BS3_hap2_chr7
BS4_hap1_chr7
BS4_hap2_chr7
BS5_hap1_chr7
BS5_hap2_chr7
BS6_hap1_chr7
BS6_hap2_chr7
CS10_hap1_chr7
CS10_hap2_chr7
CS2_hap1_chr7
CS2_hap2_chr7
CS4_hap1_chr7
CS4_hap2_chr7
CS5_hap1_chr7
CS5_hap2_chr7
CS7_hap1_chr7
CS7_hap2_chr7
CS8_hap1_chr7
CS8_hap2_chr7
NSSH10_hap1_chr7
NSSH10_hap2_chr7
NSSH2_hap1_chr7
NSSH2_hap2_chr7
Reference_chr7
DBA3.1
DBB3.1
DBB3.1Null
0 b
3 kb
6 kb
9 kb

### Locus4

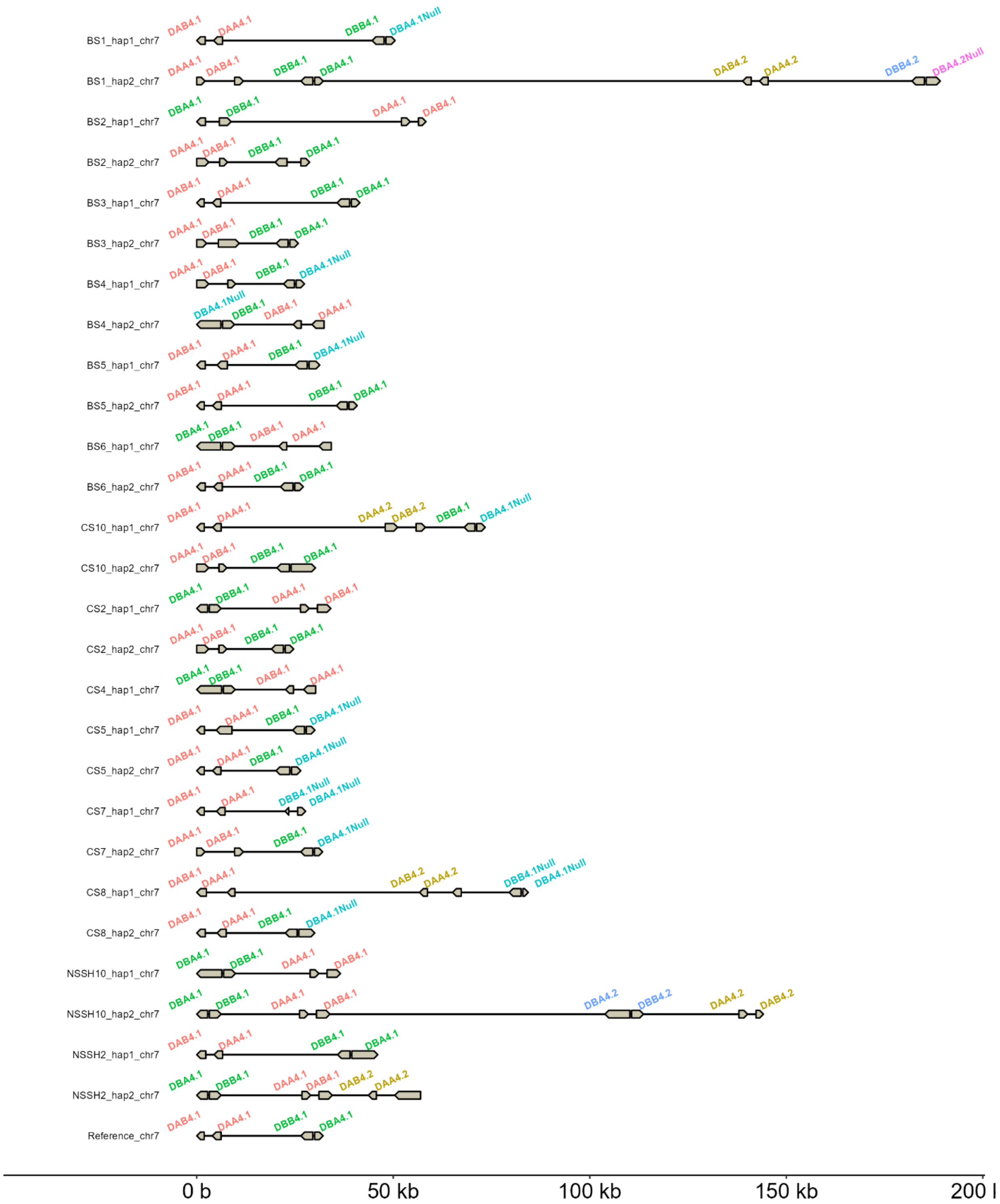

BS1_hap1_chr7
DAB4.1 DAA4.1 DBB4.1 DBA4.1Null
BS1_hap2_chr7
DAA4.1 DAB4.1 DBB4.1 DBA4.1 DAB4.2 DAA4.2 DBB4.2 DBA4.2Null
BS2_hap1_chr7
DBA4.1 DBB4.1 DAA4.1 DAB4.1
BS2_hap2_chr7
DAA4.1 DAB4.1 DBB4.1 DBA4.1
BS3_hap1_chr7
DAB4.1 DAA4.1 DBB4.1 DBA4.1
BS3_hap2_chr7
DAA4.1 DAB4.1 DBB4.1 DBA4.1
BS4_hap1_chr7
DAA4.1 DAB4.1 DBB4.1 DBA4.1Null
BS4_hap2_chr7
DBA4.1Null DBB4.1 DAB4.1 DAA4.1
BS5_hap1_chr7
DAB4.1 DAA4.1 DBB4.1 DBA4.1Null
BS5_hap2_chr7
DAB4.1 DAA4.1 DBB4.1 DBA4.1
BS6_hap1_chr7
DBA4.1 DBB4.1 DAB4.1 DAA4.1
BS6_hap2_chr7
DAB4.1 DAA4.1 DBB4.1 DBA4.1
CS10_hap1_chr7
DAB4.1 DAA4.1 DAA4.2 DAB4.2 DBB4.1 DBA4.1Null
CS10_hap2_chr7
DAA4.1 DAB4.1 DBB4.1 DBA4.1
CS2_hap1_chr7
DBA4.1 DBB4.1 DAA4.1 DAB4.1
CS2_hap2_chr7
DAA4.1 DAB4.1 DBB4.1 DBA4.1
CS4_hap1_chr7
DBA4.1 DBB4.1 DAB4.1 DAA4.1
CS5_hap1_chr7
DAB4.1 DAA4.1 DBB4.1 DBA4.1Null
CS5_hap2_chr7
DAB4.1 DAA4.1 DBB4.1 DBA4.1Null
CS7_hap1_chr7
DAB4.1 DAA4.1 DBB4.1Null DBA4.1Null
CS7_hap2_chr7
DAA4.1 DAB4.1 DBB4.1 DBA4.1Null
CS8_hap1_chr7
DAB4.1 DAA4.1 DAB4.2 DAA4.2 DBB4.1Null DBA4.1Null
CS8_hap2_chr7
DAB4.1 DAA4.1 DBB4.1 DBA4.1Null
NSSH10_hap1_chr7
DBA4.1 DBB4.1 DAA4.1 DAB4.1
NSSH10_hap2_chr7
DBA4.1 DBB4.1 DAA4.1 DAB4.1 DBA4.2 DBB4.2 DAA4.2 DAB4.2
NSSH2_hap1_chr7
DAB4.1 DAA4.1 DBB4.1 DBA4.1
NSSH2_hap2_chr7
DBA4.1 DBB4.1 DAA4.1 DAB4.1 DAB4.2 DAA4.2
Reference_chr7
DAB4.1 DAA4.1 DBB4.1 DBA4.1
0 b
50 kb
100 kb
150 kb
200 l

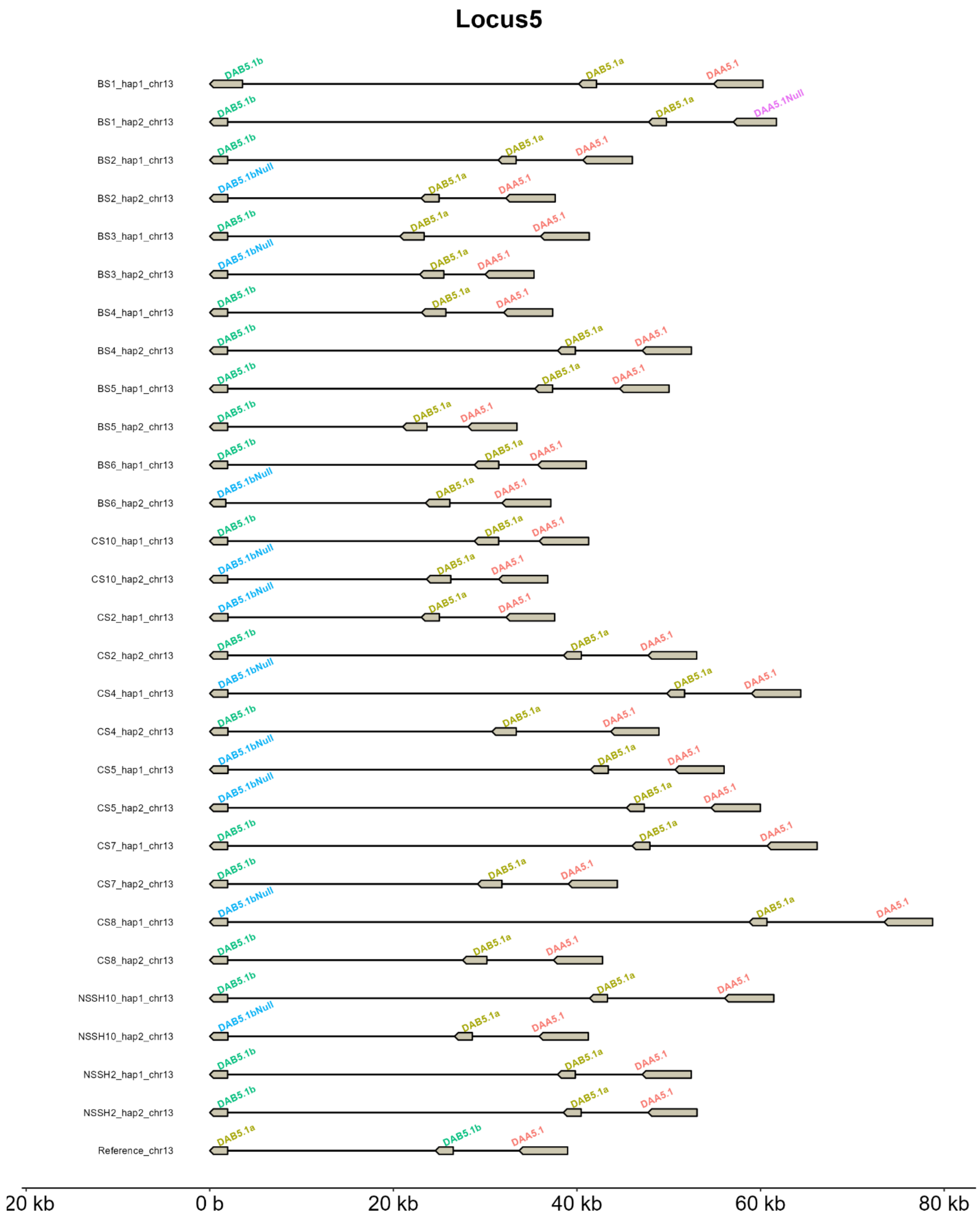
Locus5
BS1_hap1_chr13
BS1_hap2_chr13
BS2_hap1_chr13
BS2_hap2_chr13
BS3_hap1_chr13
BS3_hap2_chr13
BS4_hap1_chr13
BS4_hap2_chr13
BS5_hap1_chr13
BS5_hap2_chr13
BS6_hap1_chr13
BS6_hap2_chr13
CS10_hap1_chr13
CS10_hap2_chr13
CS2_hap1_chr13
CS2_hap2_chr13
CS4_hap1_chr13
CS4_hap2_chr13
CS5_hap1_chr13
CS5_hap2_chr13
CS7_hap1_chr13
CS7_hap2_chr13
CS8_hap1_chr13
CS8_hap2_chr13
NSSH10_hap1_chr13
NSSH10_hap2_chr13
NSSH2_hap1_chr13
NSSH2_hap2_chr13
Reference_chr13
DAB5.1b
DAB5.1bNull
DAB5.1a
DAA5.1
DAA5.1Null
20 kb
0 b
20 kb
40 kb
60 kb
80 kb

### Locus6

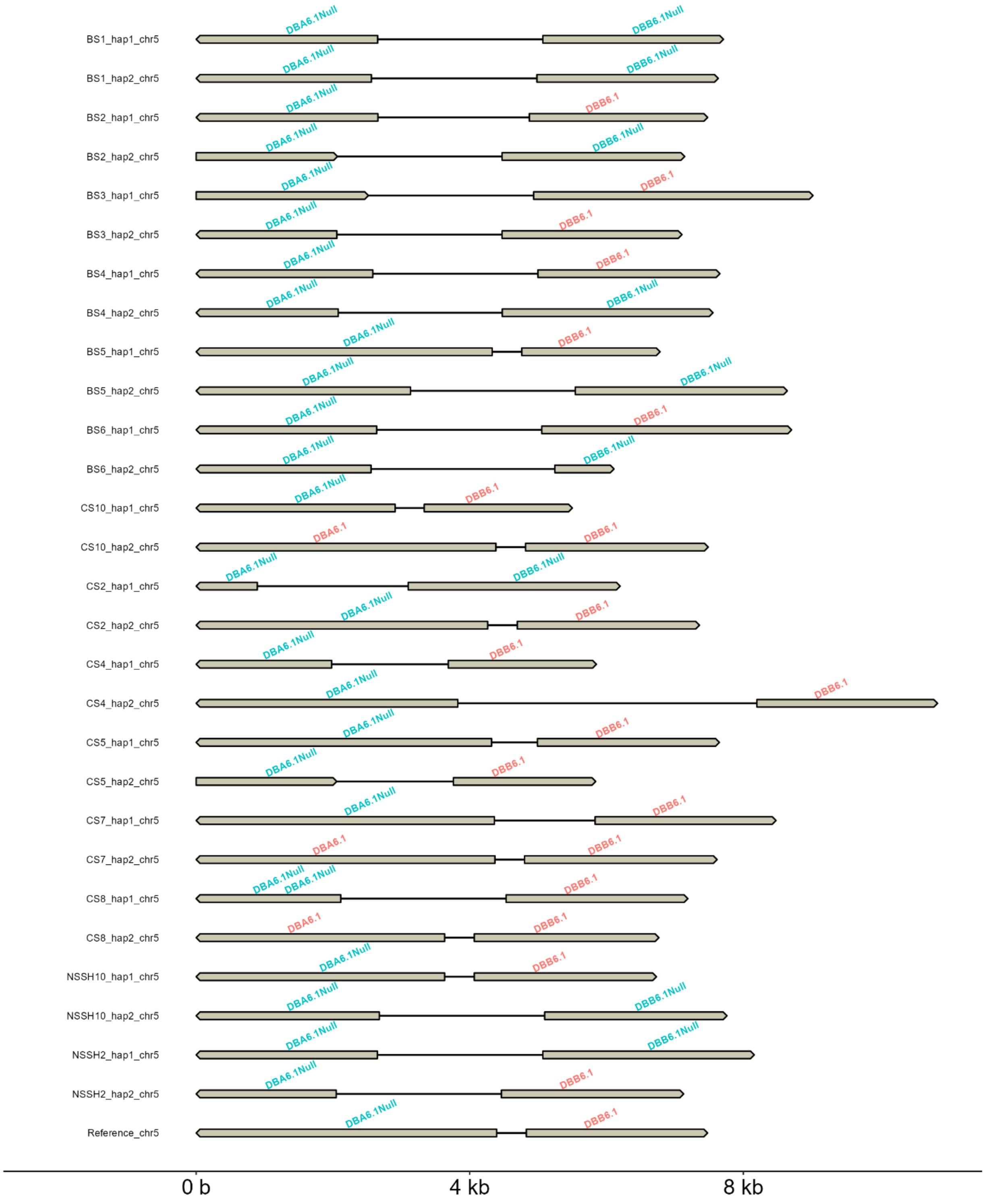
BS1_hap1_chr5
DBA6.1Null
DBB6.1Null
BS1_hap2_chr5
DBA6.1Null
DBB6.1Null
BS2_hap1_chr5
DBA6.1Null
DBB6.1
BS2_hap2_chr5
DBA6.1Null
DBB6.1Null
BS3_hap1_chr5
DBA6.1Null
DBB6.1
BS3_hap2_chr5
DBA6.1Null
DBB6.1
BS4_hap1_chr5
DBA6.1Null
DBB6.1
BS4_hap2_chr5
DBA6.1Null
DBB6.1Null
BS5_hap1_chr5
DBA6.1Null
DBB6.1
BS5_hap2_chr5
DBA6.1Null
DBB6.1Null
BS6_hap1_chr5
DBA6.1Null
DBB6.1
BS6_hap2_chr5
DBA6.1Null
DBB6.1Null
CS10_hap1_chr5
DBA6.1Null
DBB6.1
CS10_hap2_chr5
DBA6.1
DBB6.1
CS2_hap1_chr5
DBA6.1Null
DBB6.1Null
CS2_hap2_chr5
DBA6.1Null
DBB6.1
CS4_hap1_chr5
DBA6.1Null
DBB6.1
CS4_hap2_chr5
DBA6.1Null
DBB6.1
CS5_hap1_chr5
DBA6.1Null
DBB6.1
CS5_hap2_chr5
DBA6.1Null
DBB6.1
CS7_hap1_chr5
DBA6.1Null
DBB6.1
CS7_hap2_chr5
DBA6.1
DBB6.1
CS8_hap1_chr5
DBA6.1Null
DBA6.1Null
DBB6.1
CS8_hap2_chr5
DBA6.1
DBB6.1
NSSH10_hap1_chr5
DBA6.1Null
DBB6.1
NSSH10_hap2_chr5
DBA6.1Null
DBB6.1Null
NSSH2_hap1_chr5
DBA6.1Null
DBB6.1Null
NSSH2_hap2_chr5
DBA6.1Null
DBB6.1
Reference_chr5
DBA6.1Null
DBB6.1
0 b
4 kb
8 kb

### Locus7

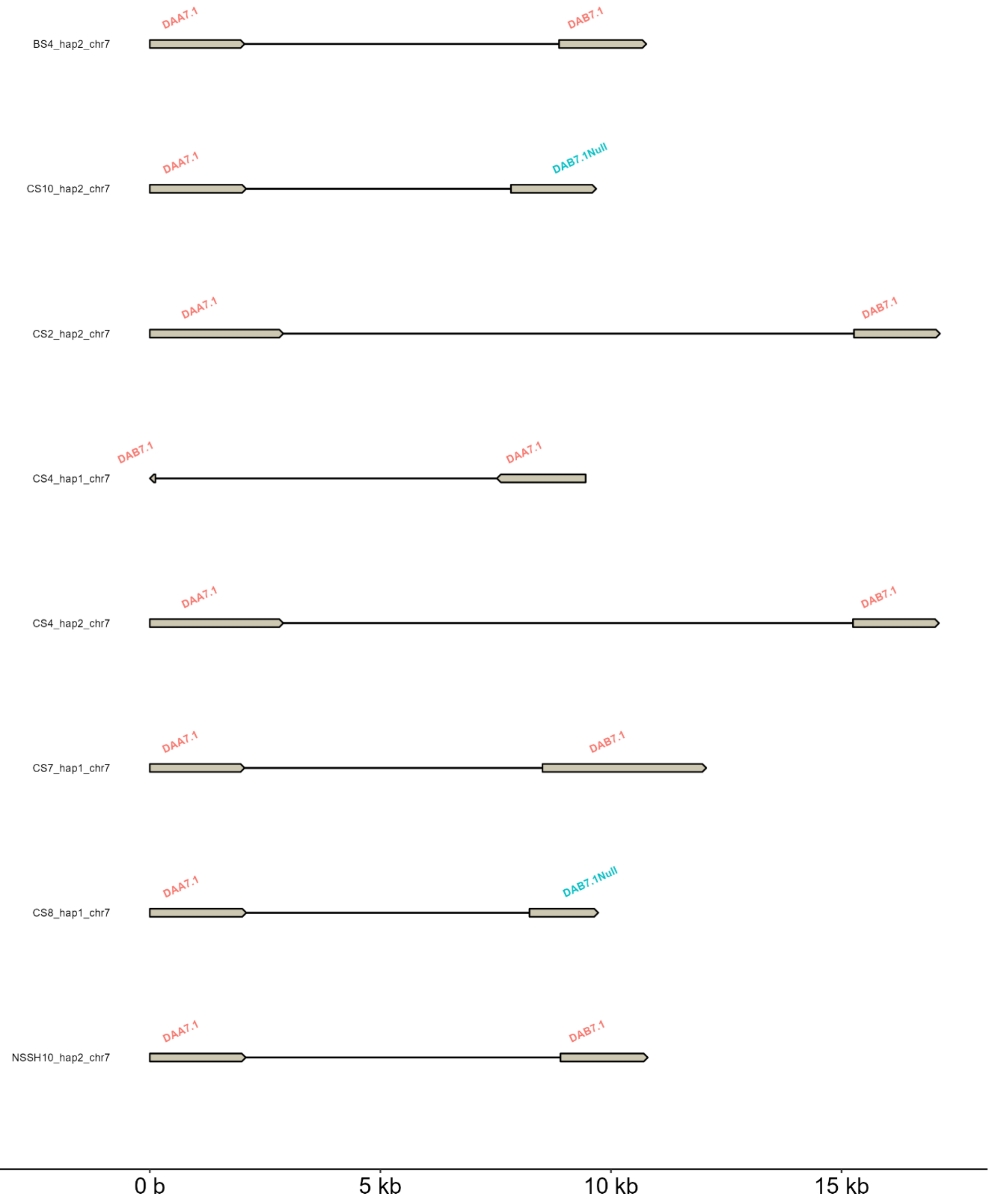

BS4_hap2_chr7
DAA7.1
DAB7.1
CS10_hap2_chr7
DAA7.1
DAB7.1Null
CS2_hap2_chr7
DAA7.1
DAB7.1
CS4_hap1_chr7
DAB7.1
DAA7.1
CS4_hap2_chr7
DAA7.1
DAB7.1
CS7_hap1_chr7
DAA7.1
DAB7.1
CS8_hap1_chr7
DAA7.1
DAB7.1Null
NSSH10_hap2_chr7
DAA7.1
DAB7.1
0 b
5 kb
10 kb
15 kb

### Locus8

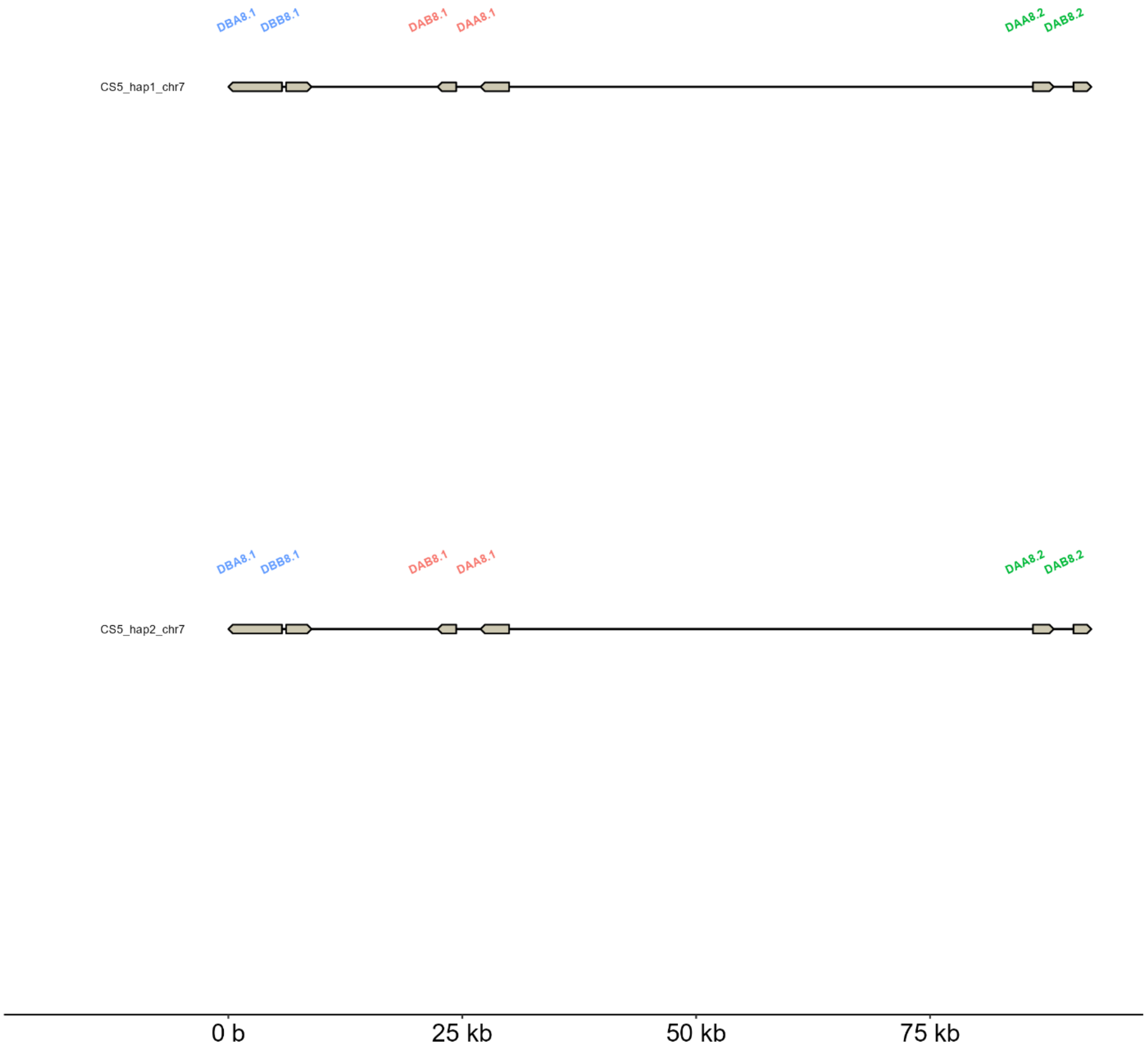
DBA8.1
DBB8.1
DAB8.1
DAA8.1
DAA8.2
DAB8.2
CS5_hap1_chr7
DBA8.1
DBB8.1
DAB8.1
DAA8.1
DAA8.2
DAB8.2
CS5_hap2_chr7
0 b
25 kb
50 kb
75 kb

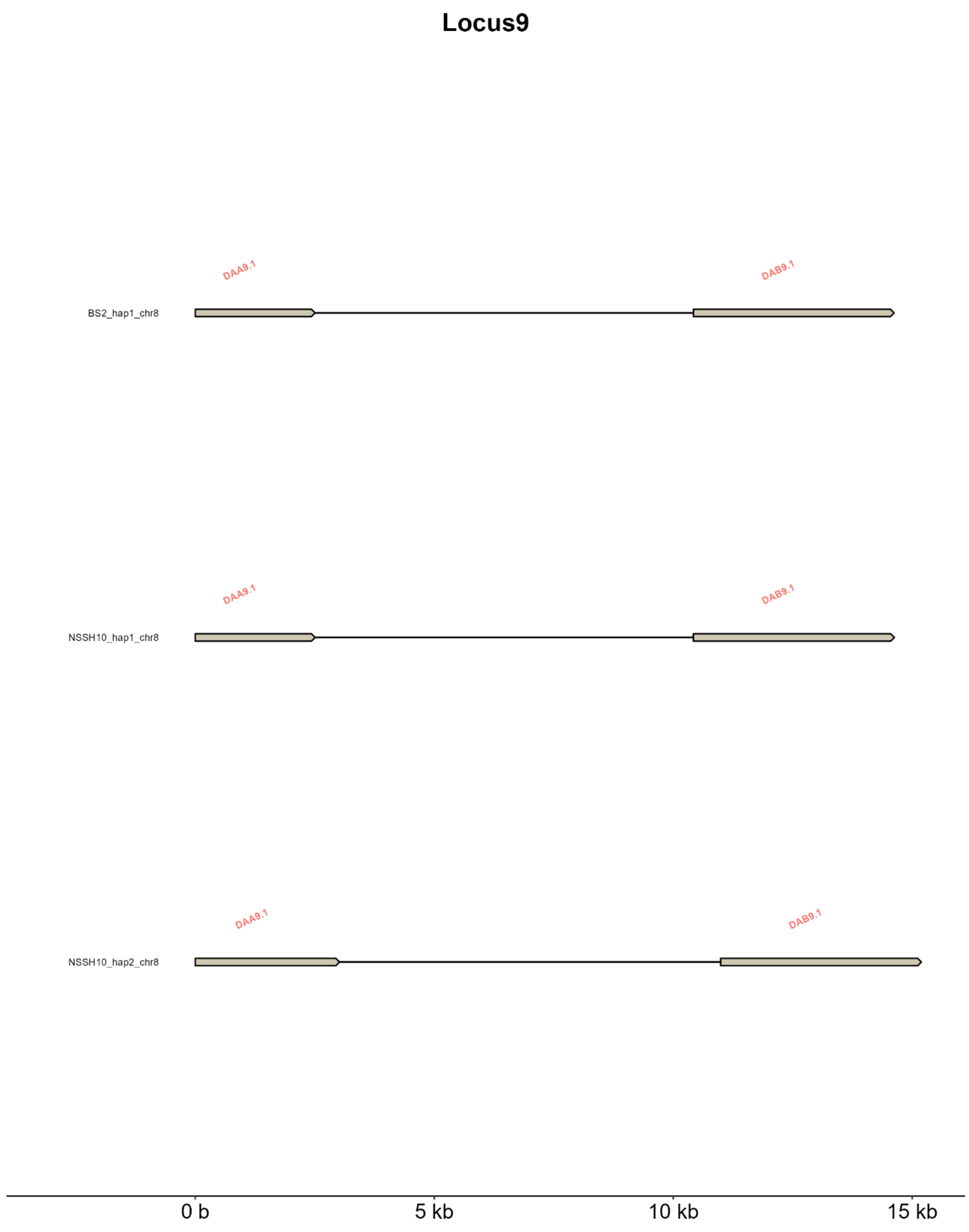


**Supplementary Fig. 2**. Gene organization of MHC class II on nine loci in all 29 haplotypes.

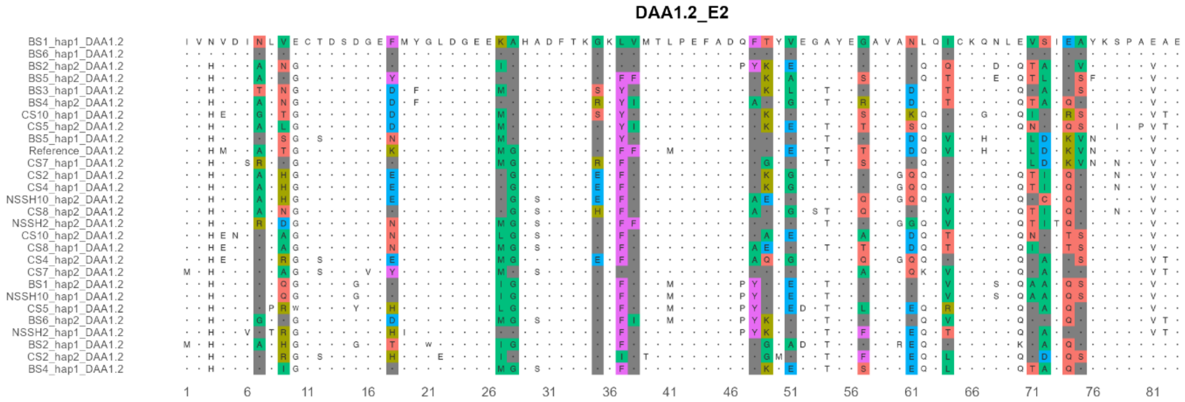
DAA1.2_E2

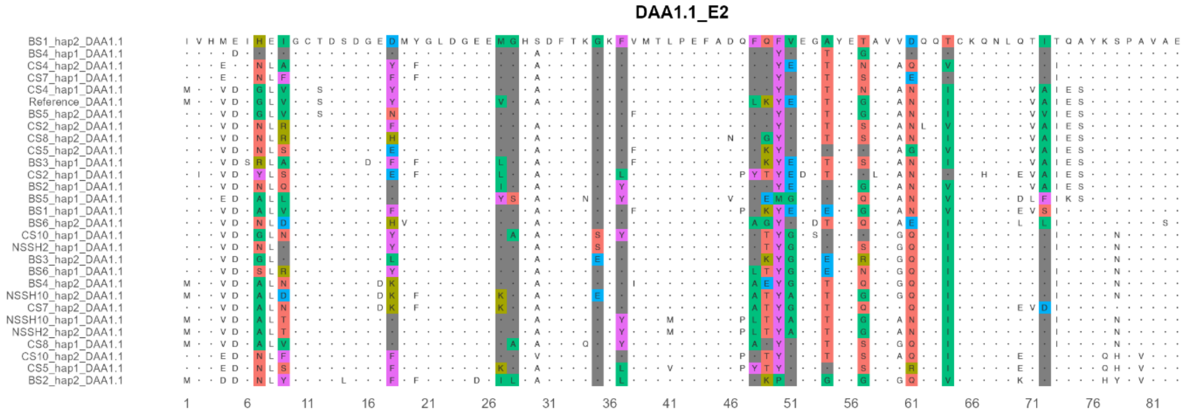
DAA1.1_E2

#### DAA1.3_E2

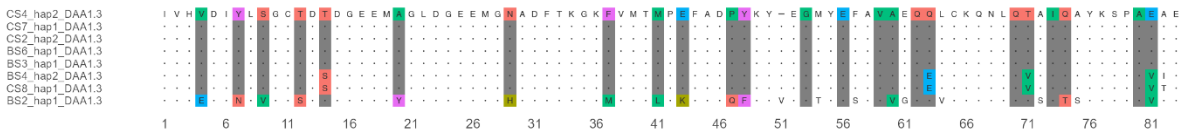

#### DAA2.1_E2

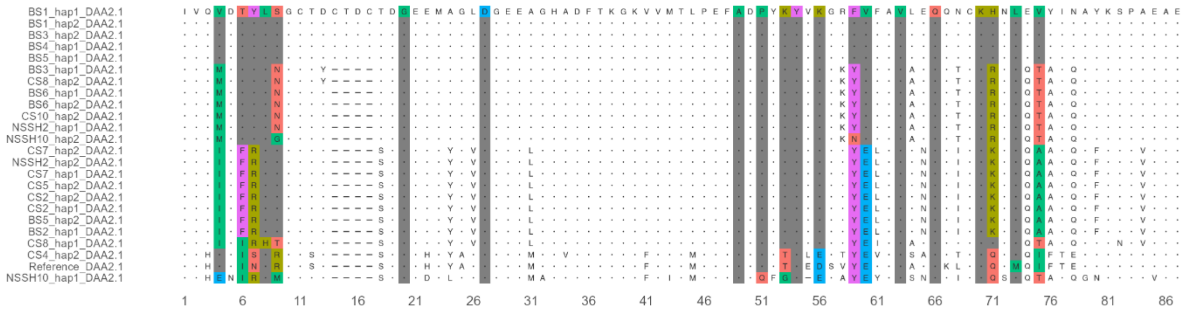

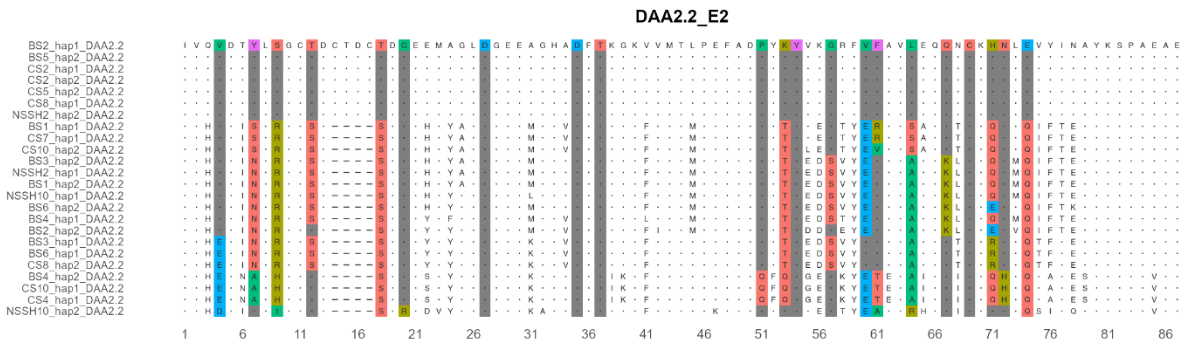
DAA2.2_E2
BS2_hap1_DAA2.2
BS5_hap2_DAA2.2
CS2_hap1_DAA2.2
CS2_hap2_DAA2.2
CS5_hap2_DAA2.2
CS8_hap1_DAA2.2
NSSH2_hap2_DAA2.2
BS1_hap1_DAA2.2
CS7_hap1_DAA2.2
CS10_hap2_DAA2.2
BS3_hap2_DAA2.2
NSSH2_hap1_DAA2.2
BS1_hap2_DAA2.2
NSSH10_hap1_DAA2.2
BS6_hap2_DAA2.2
BS4_hap1_DAA2.2
BS2_hap2_DAA2.2
BS3_hap1_DAA2.2
BS6_hap1_DAA2.2
CS8_hap2_DAA2.2
BS4_hap2_DAA2.2
CS10_hap1_DAA2.2
CS4_hap1_DAA2.2
NSSH10_hap2_DAA2.2

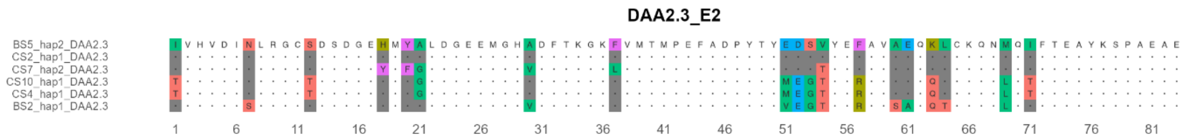
DAA2.3_E2
BS5_hap2_DAA2.3
CS2_hap1_DAA2.3
CS7_hap2_DAA2.3
CS10_hap1_DAA2.3
CS4_hap1_DAA2.3
BS2_hap1_DAA2.3

**DAA4.1_E2**

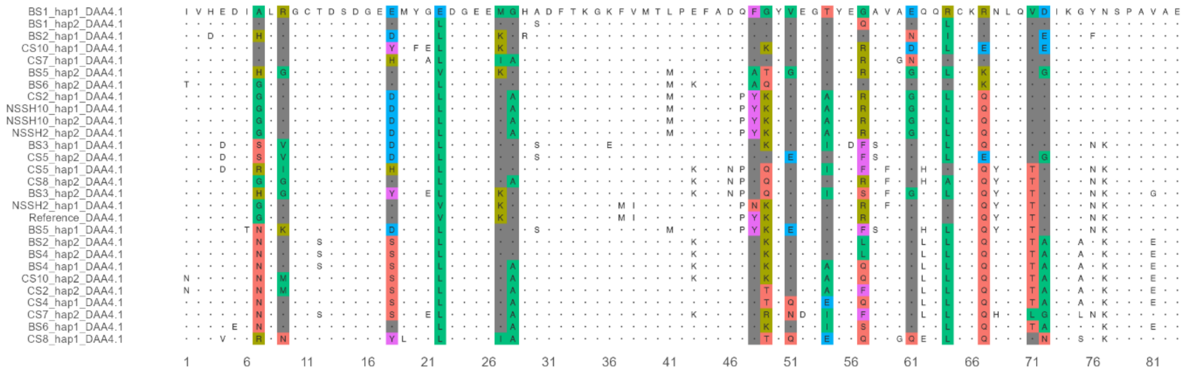


**DAA4.2_E2**

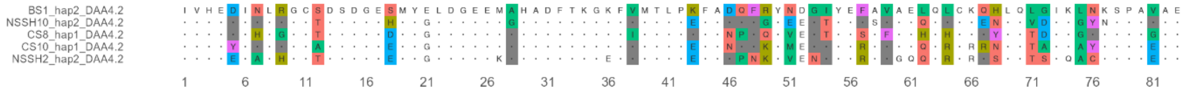

**DAA7.1_E2**

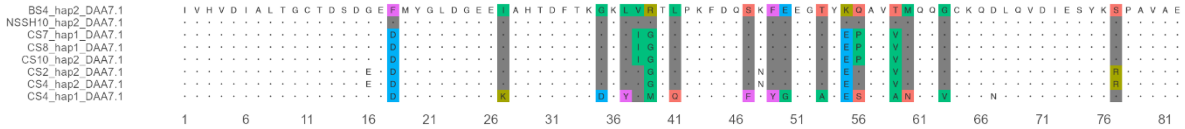


**DAB1.1_E2**

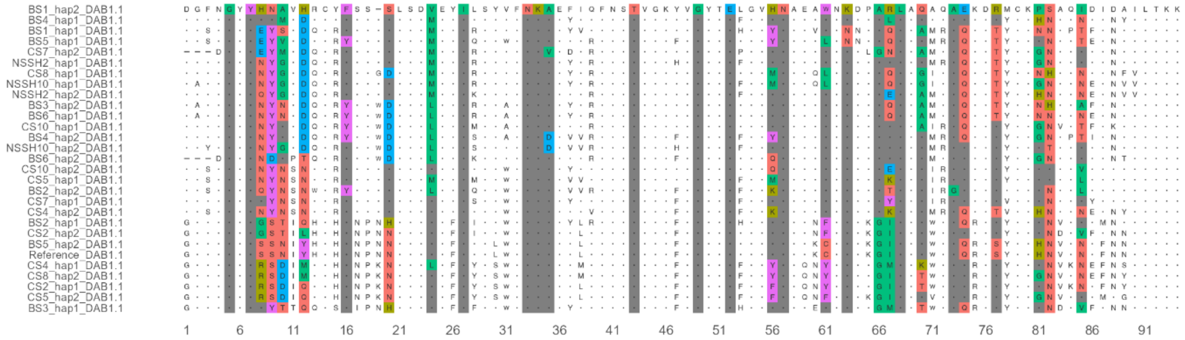

**DAB1.2_E2**

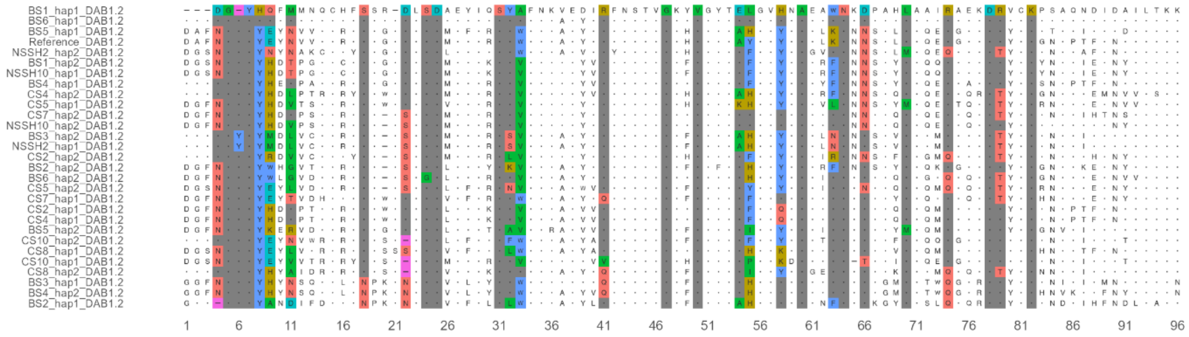


**DAB1.3_E2**

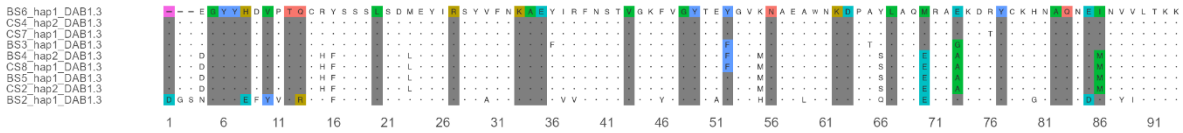

**DAB2.1_E2**

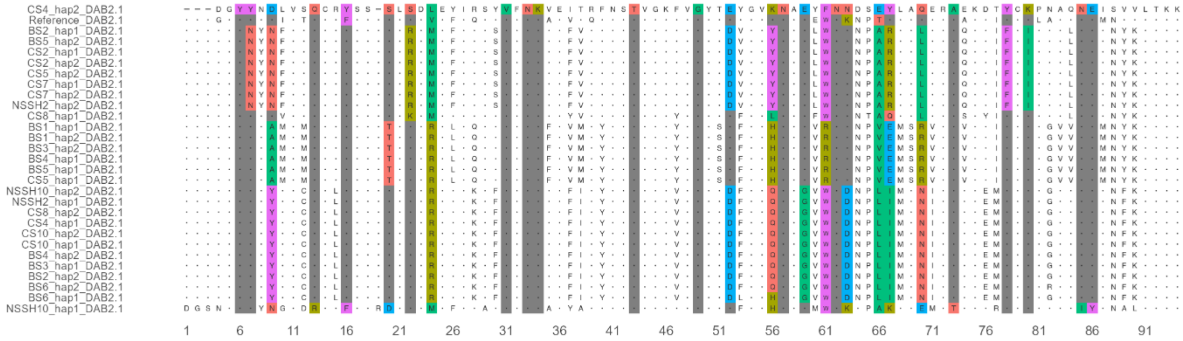

**DAB2.3_E2**

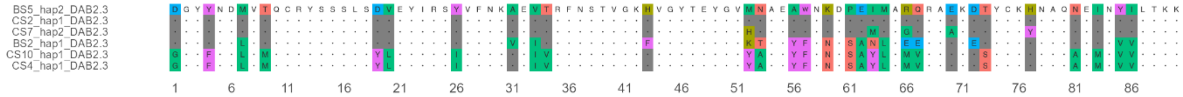

DAB2.2_E2

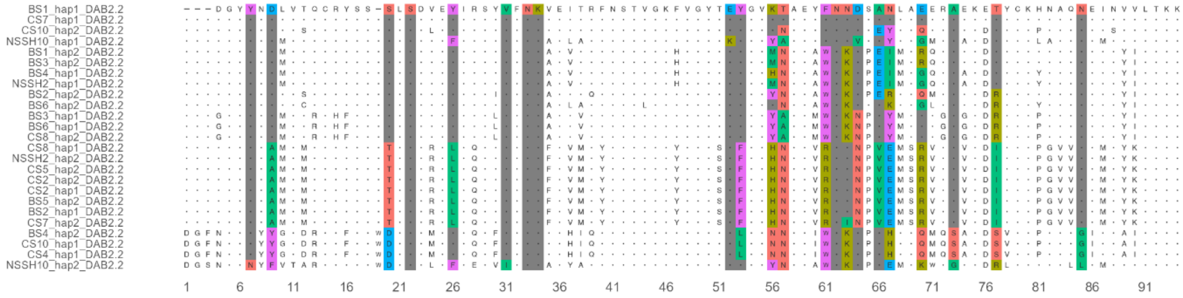


DAB4.1_E2

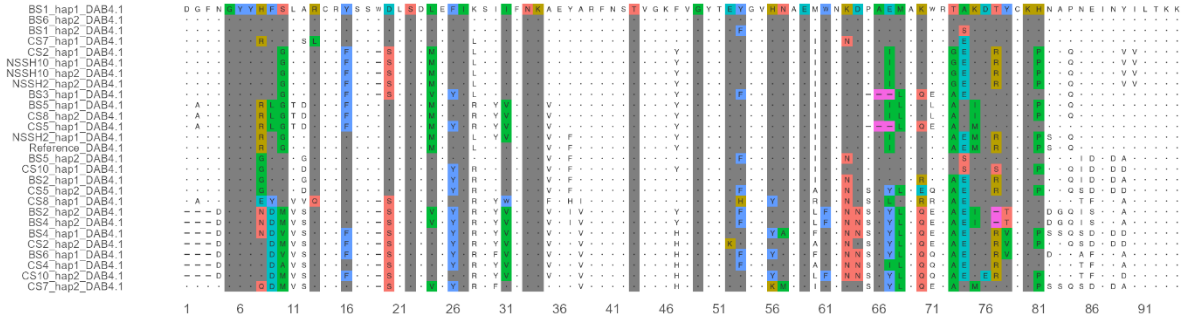

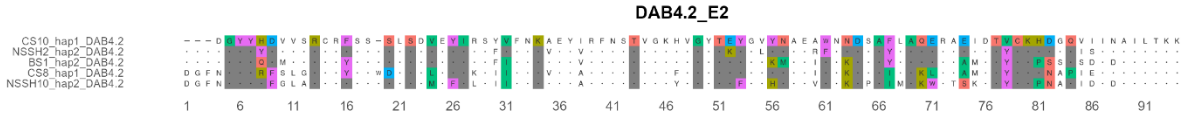
DAB4.2_E2

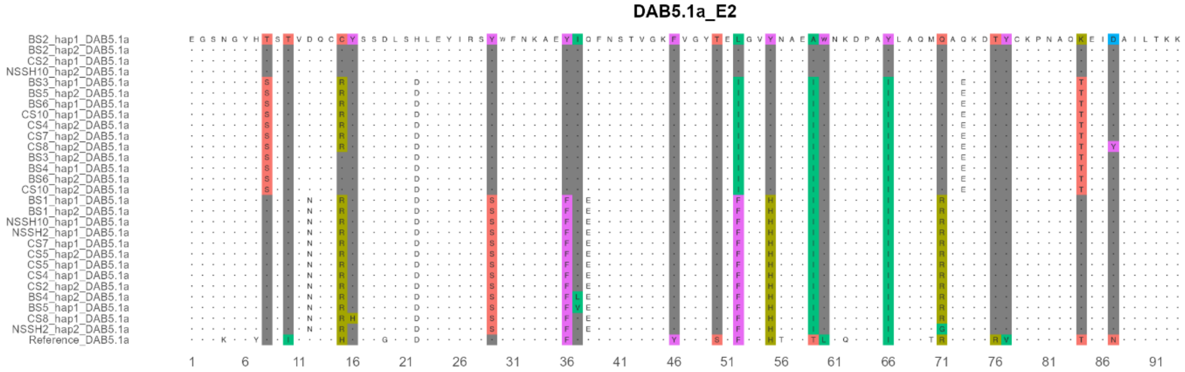
DAB5.1a_E2

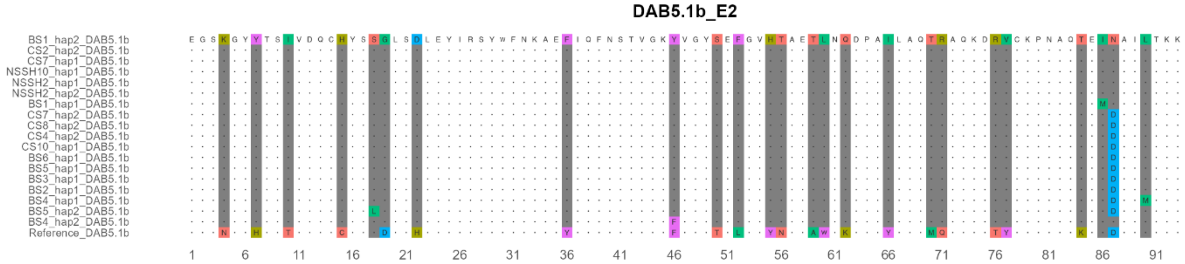
DAB5.1b_E2
BS1_hap2_DAB5.1b
CS2_hap2_DAB5.1b
CS7_hap1_DAB5.1b
NSSH10_hap1_DAB5.1b
NSSH2_hap1_DAB5.1b
NSSH2_hap2_DAB5.1b
BS1_hap1_DAB5.1b
CS7_hap2_DAB5.1b
CS8_hap2_DAB5.1b
CS4_hap2_DAB5.1b
CS10_hap1_DAB5.1b
BS6_hap1_DAB5.1b
BS5_hap1_DAB5.1b
BS3_hap1_DAB5.1b
BS2_hap1_DAB5.1b
BS4_hap1_DAB5.1b
BS5_hap2_DAB5.1b
BS4_hap2_DAB5.1b
Reference_DAB5.1b

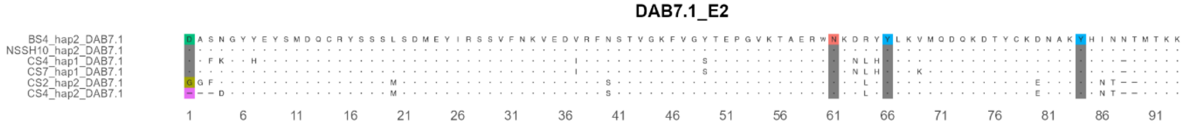
DAB7.1_E2
BS4_hap2_DAB7.1
NSSH10_hap2_DAB7.1
CS4_hap1_DAB7.1
CS7_hap1_DAB7.1
CS2_hap2_DAB7.1
CS4_hap2_DAB7.1

**DBA3.1_E2**

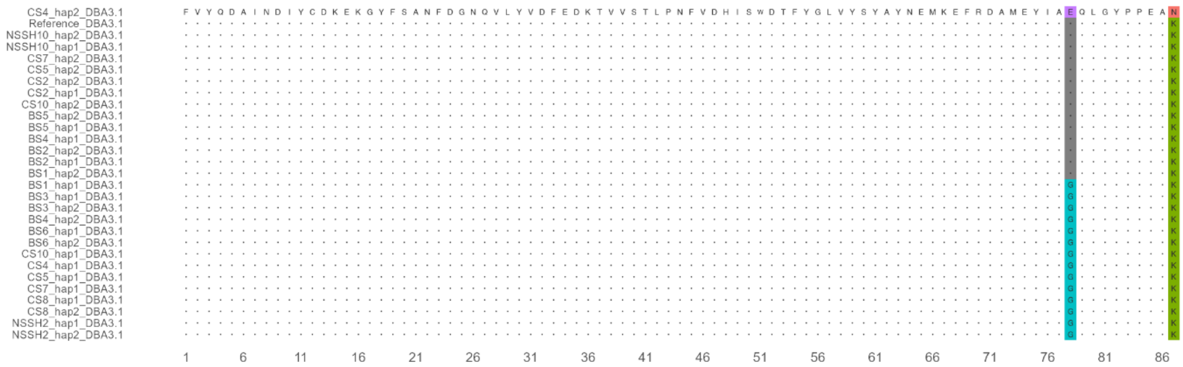


**DBA4.1_E2**

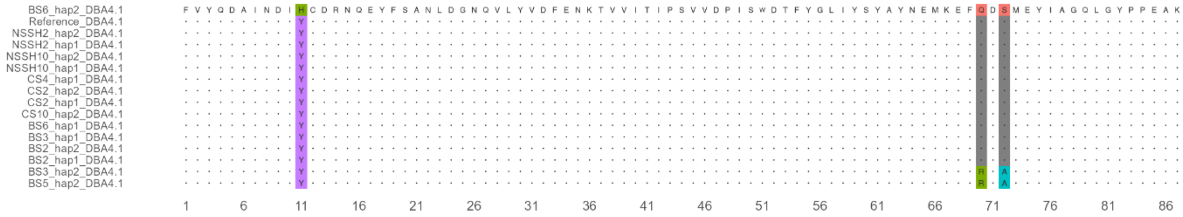

**DAA1.1_E3**

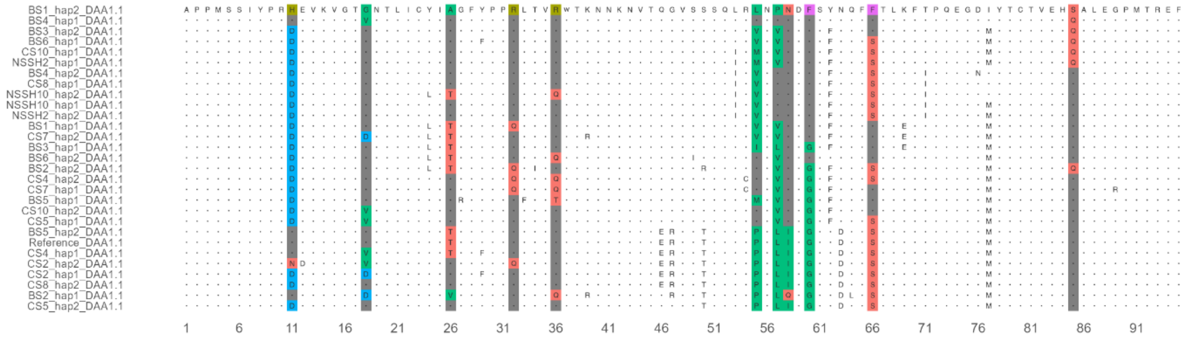


**DAA1.2_E3**

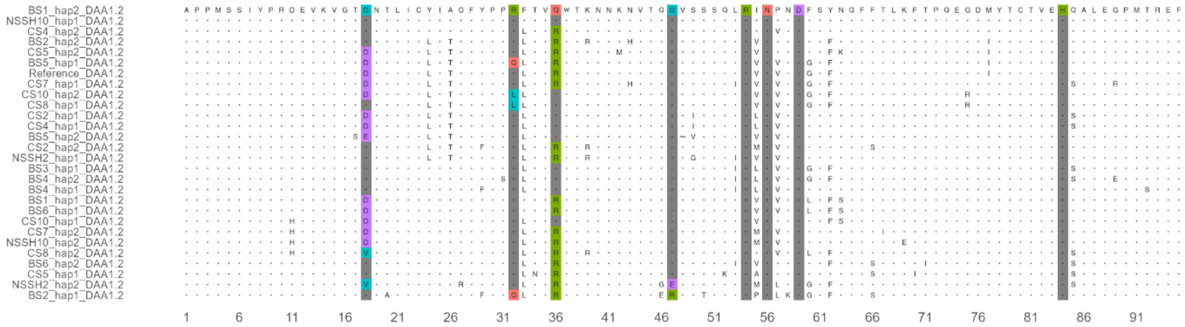

#### DAA1.3_E3

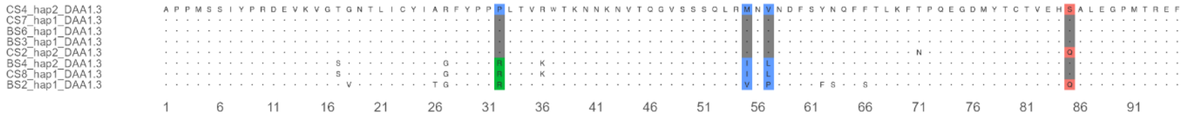


#### DAA2.1_E3

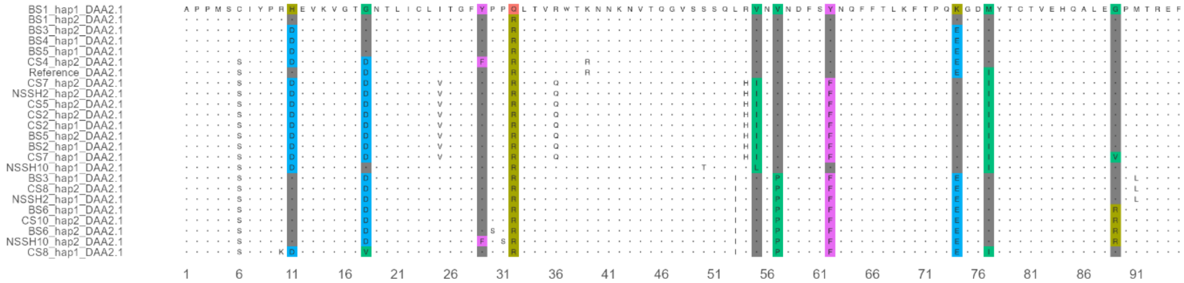

DAA2.2_E3

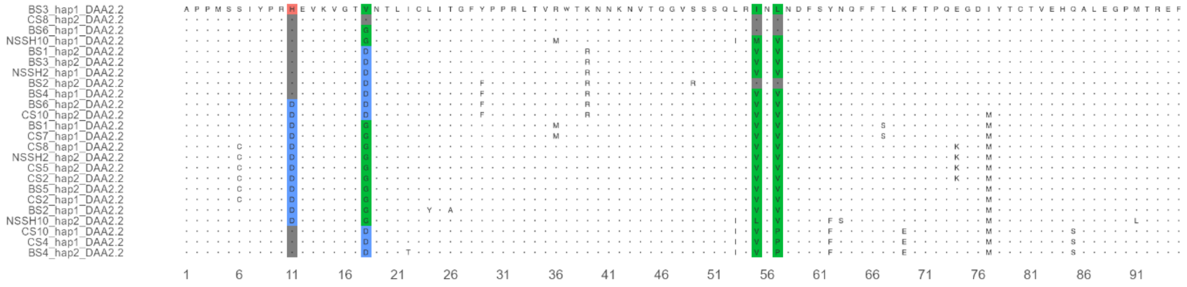

DAA2.3_E3

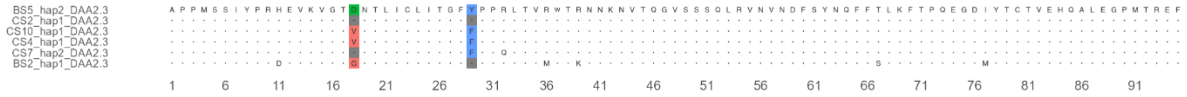

DAA4.1_E3

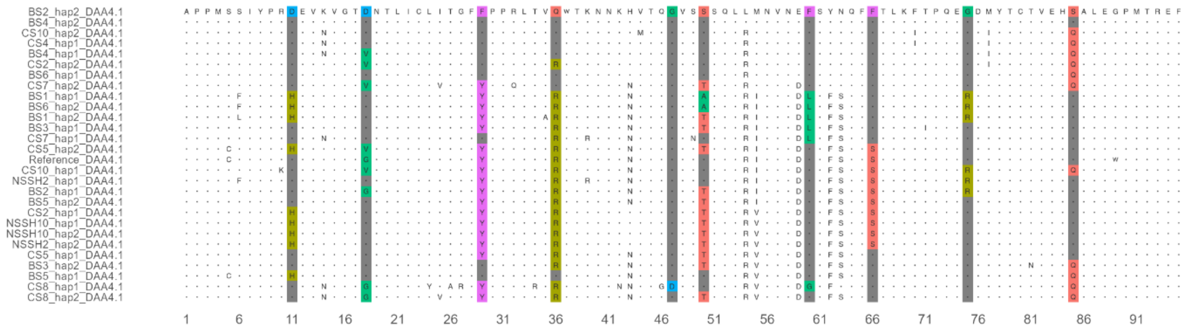


DAA4.2_E3

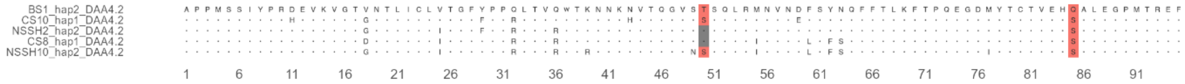

**DAA7.1_E3**

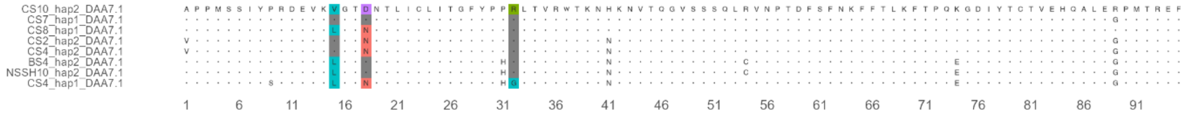


**DAB1.1_E3**

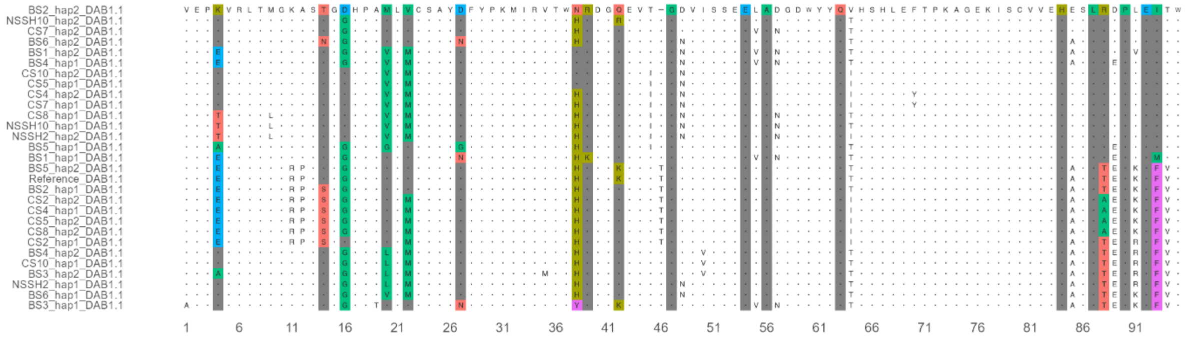

#### DAB1.2_E3

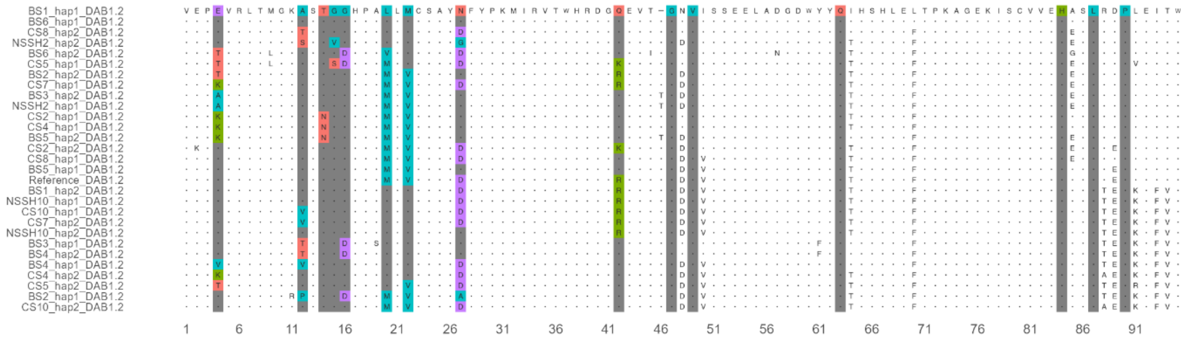

#### DAB1.3_E3

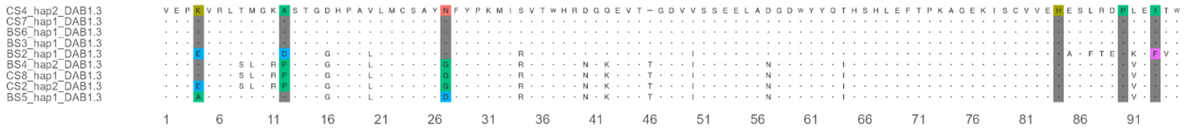

DAB2.1_E3

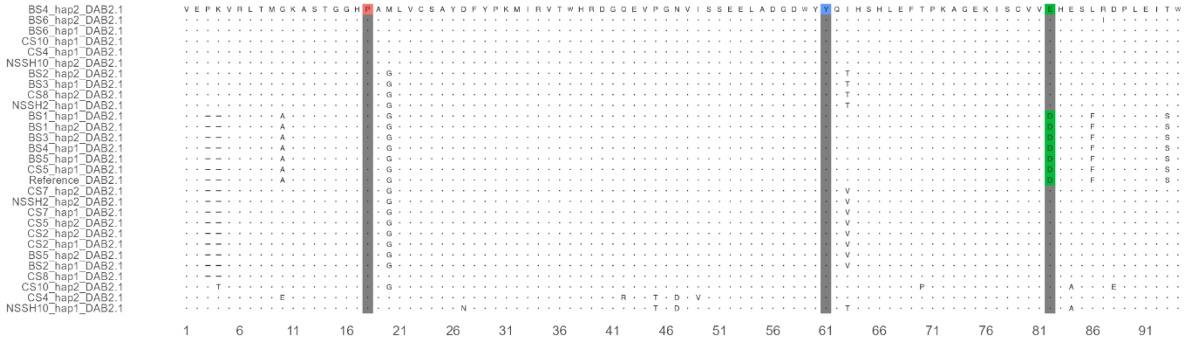

DAB2.2_E3

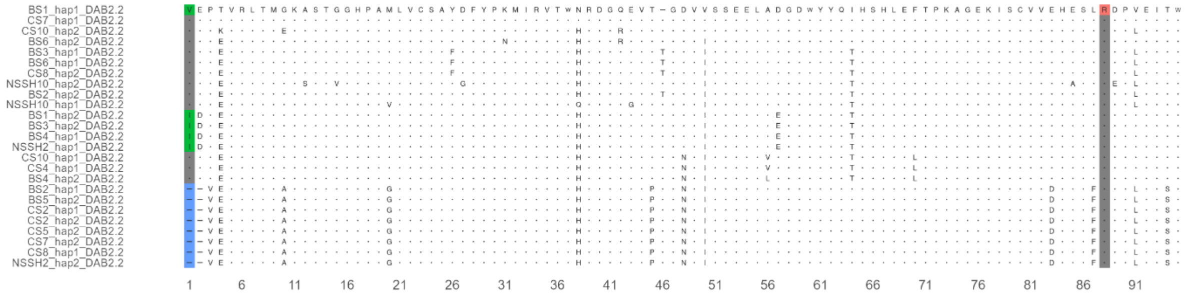

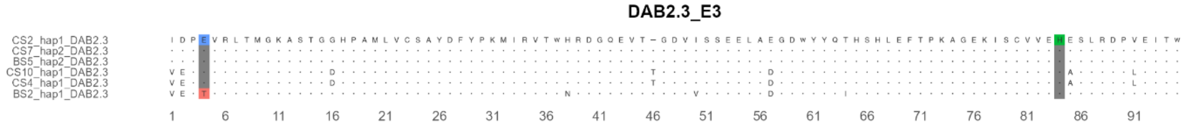
DAB2.3_E3

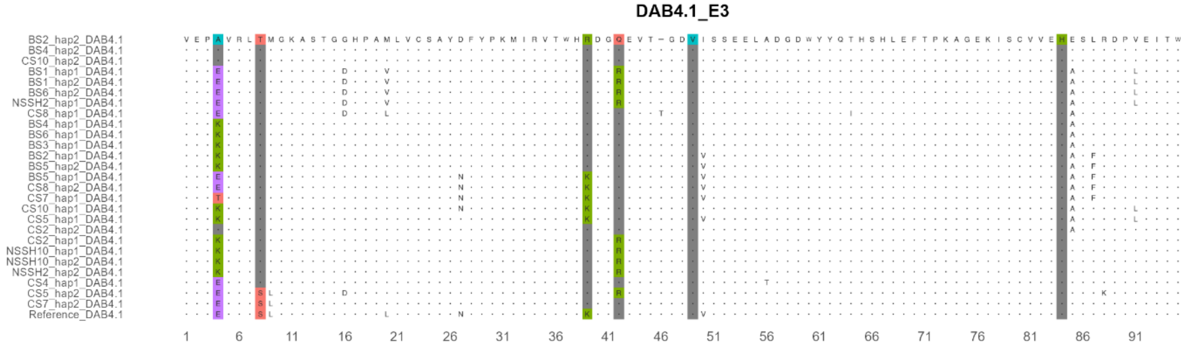
DAB4.1_E3

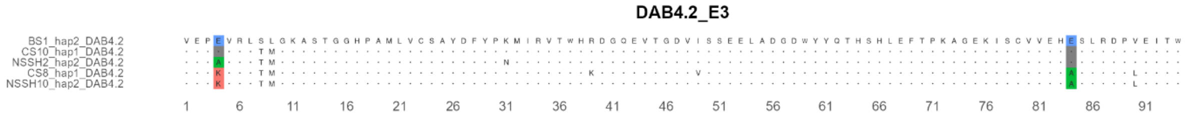
DAB4.2_E3
BS1_hap2_DAB4.2
CS10_hap1_DAB4.2
NSSH2_hap2_DAB4.2
CS8_hap1_DAB4.2
NSSH10_hap2_DAB4.2
1 6 11 16 21 26 31 36 41 46 51 56 61 66 71 76 81 86 91

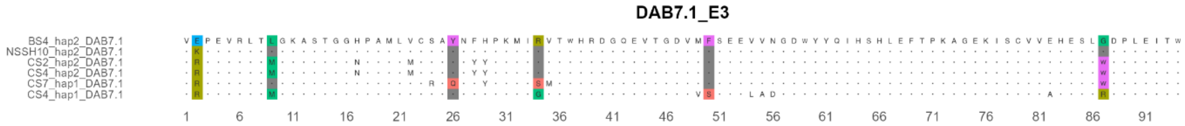
DAB7.1_E3
BS4_hap2_DAB7.1
NSSH10_hap2_DAB7.1
CS2_hap2_DAB7.1
CS4_hap2_DAB7.1
CS7_hap1_DAB7.1
CS4_hap1_DAB7.1
1 6 11 16 21 26 31 36 41 46 51 56 61 66 71 76 81 86 91

DBB2.1_E3

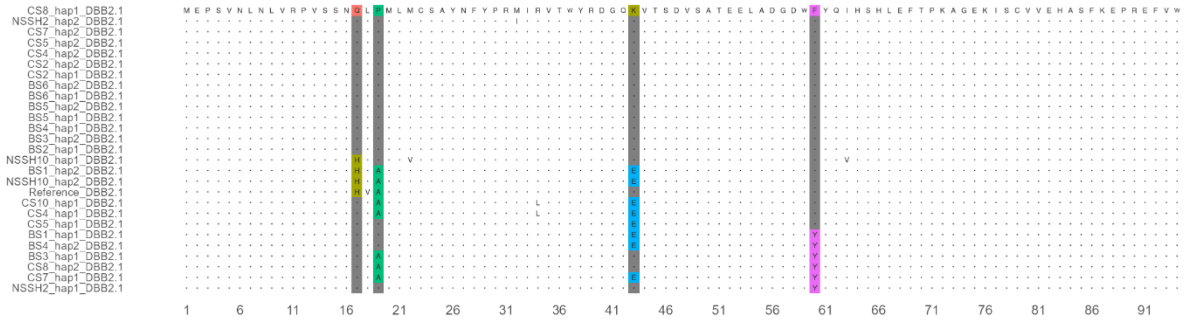


DBB4.1_E3

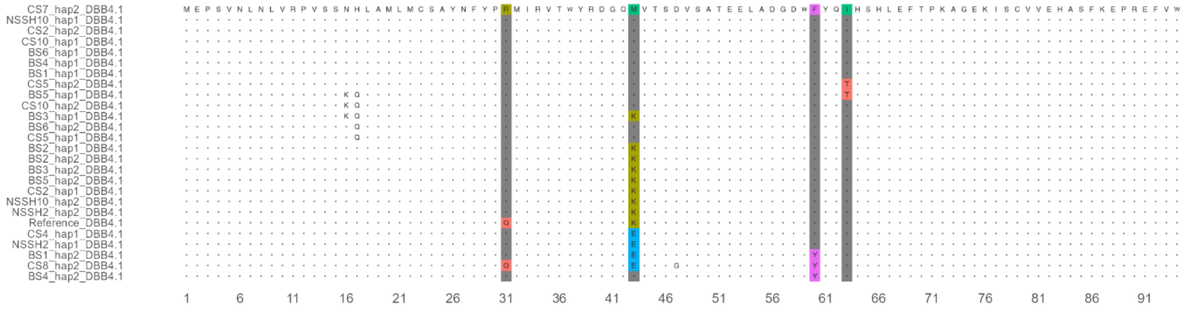

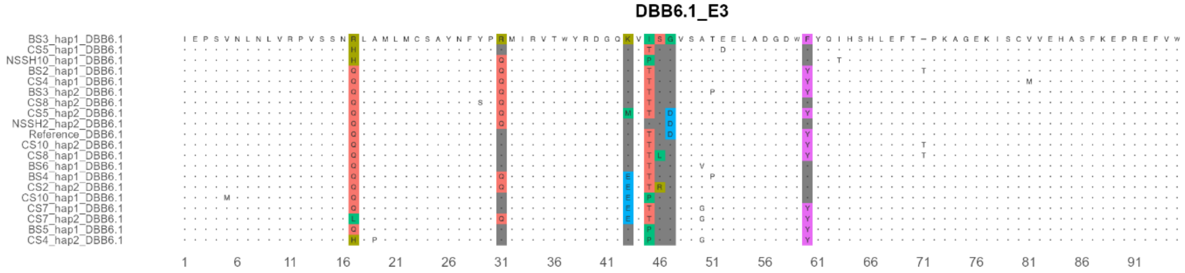


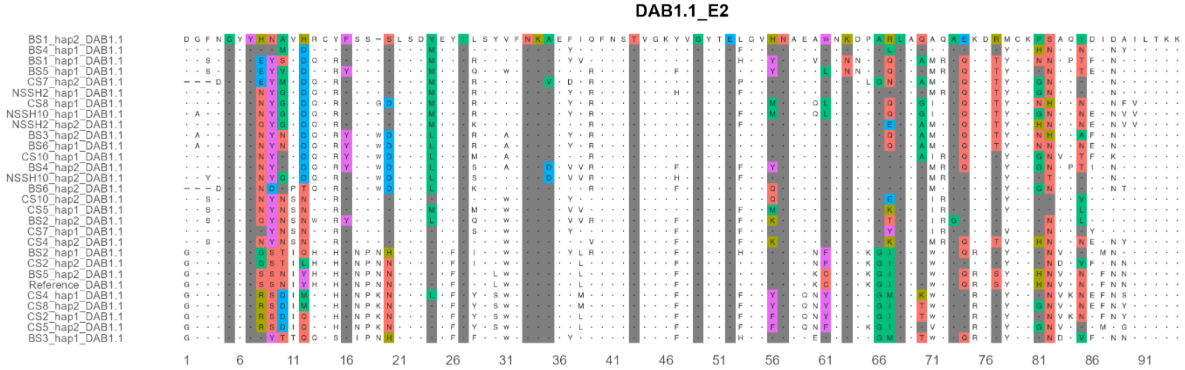


**Supplementary Fig. 3.** Amino acid sequence alignments of α1, α2, β1, and β2 domains of classical and non-classical genes used for *dN/dS* analysis. Colored residues represent positively selected sites predicted by M8 model of CODEML. The color shades represent the chemistry of amino acids where hydrophobic residues are in blue shades and hydrophilic residues are in red shades.

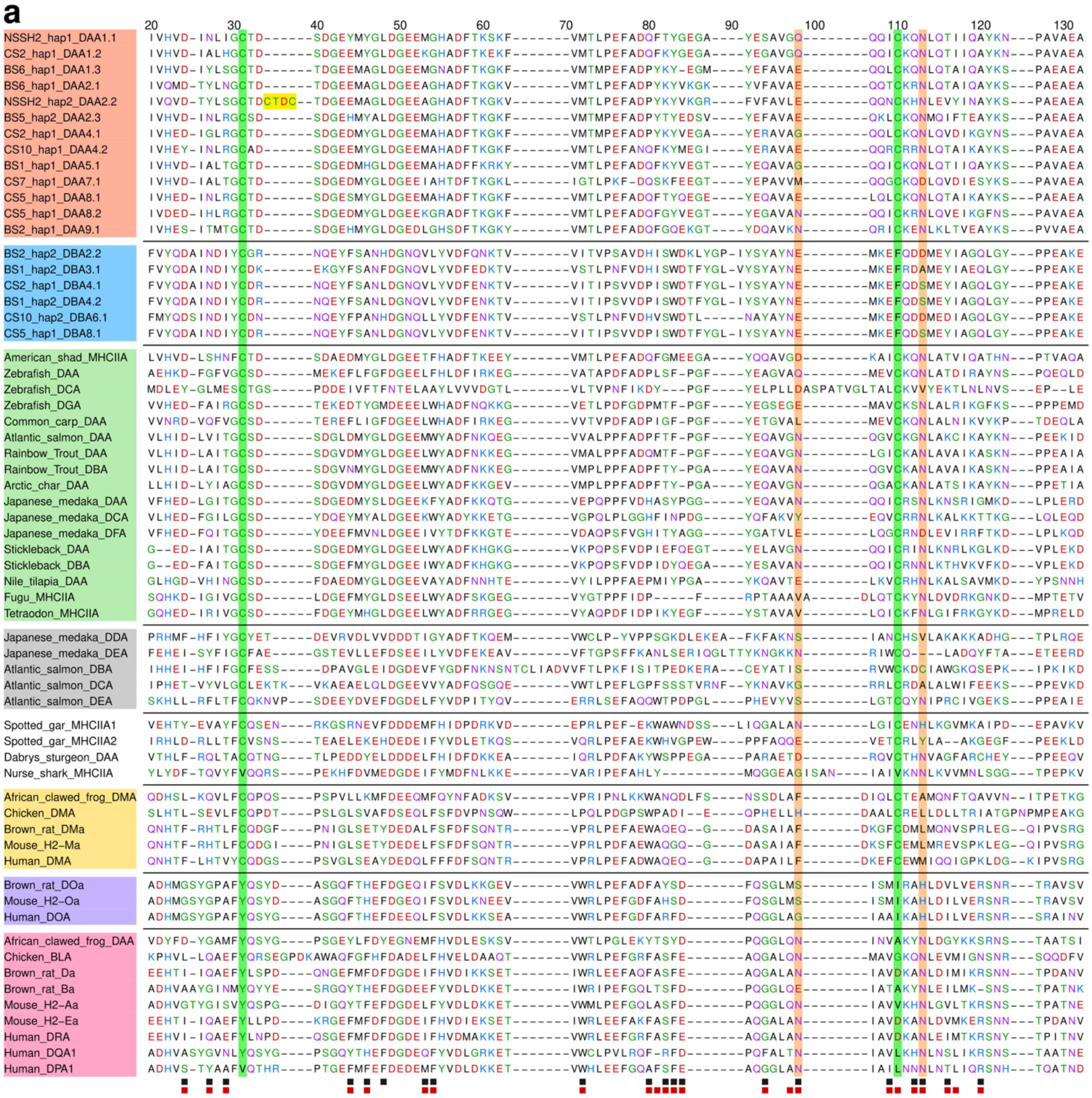
a
20
30
40
50
60
70
80
90
100
110
120
130
NSSH2_hap1_DAA1.1
CS2_hap1_DAA1.2
BS6_hap1_DAA1.3
BS6_hap1_DAA2.1
NSSH2_hap2_DAA2.2
BS5_hap2_DAA2.3
CS2_hap1_DAA4.1
CS10_hap1_DAA4.2
BS1_hap1_DAA5.1
CS7_hap1_DAA7.1
CS5_hap1_DAA8.1
CS5_hap1_DAA8.2
BS2_hap1_DAA9.1
BS2_hap2_DBA2.2
BS1_hap2_DBA3.1
CS2_hap1_DBA4.1
BS1_hap1_DBA4.2
CS10_hap2_DBA6.1
CS5_hap1_DBA8.1
American_shad_MHCIIA
Zebrafish_DAA
Zebrafish_DCA
Zebrafish_DGA
Common_carp_DAA
Atlantic_salmon_DAA
Rainbow_Trout_DAA
Rainbow_Trout_DBA
Arctic_char_DAA
Japanese_medaka_DAA
Japanese_medaka_DCA
Japanese_medaka_DFA
Stickleback_DAA
Stickleback_DBA
Nile_tilapia_DAA
Fugu_MHCIIA
Tetraodon_MHCIIA
Japanese_medaka_DDA
Japanese_medaka_DEA
Atlantic_salmon_DBA
Atlantic_salmon_DCA
Atlantic_salmon_DEA
Spotted_gar_MHCIIA1
Spotted_gar_MHCIIA2
Dabrys_sturgeon_DAA
Nurse_shark_MHCIIA
African_clawed_frog_DMA
Chicken_DMA
Brown_rat_DMa
Mouse_H2–Ma
Human_DMA
Brown_rat_DOa
Mouse_H2–Oa
Human_DOA
African_clawed_frog_DAA
Chicken_BLA
Brown_rat_Da
Brown_rat_Ba
Mouse_H2–Aa
Mouse_H2–Ea
Human_DRA
Human_DQA1
Human_DPA1

**b**

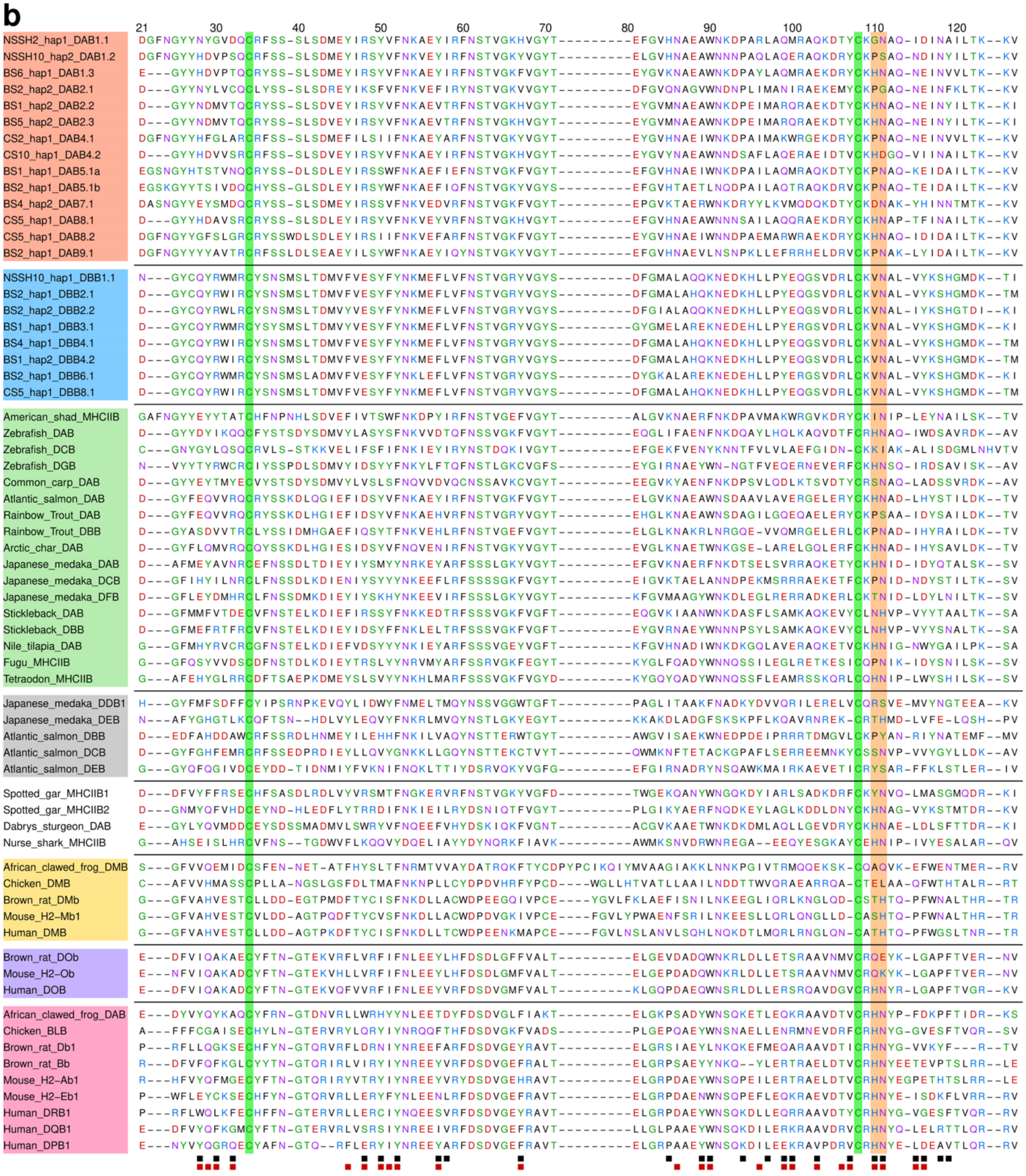

21
30
40
50
60
70
80
90
100
110
120
NSSH2_hap1_DAB1.1
NSSH10_hap2_DAB1.2
BS6_hap1_DAB1.3
BS2_hap2_DAB2.1
BS1_hap2_DAB2.2
BS5_hap2_DAB2.3
CS2_hap1_DAB4.1
CS10_hap1_DAB4.2
BS1_hap1_DAB5.1a
BS2_hap1_DAB5.1b
BS4_hap2_DAB7.1
CS5_hap1_DAB8.1
CS5_hap1_DAB8.2
BS2_hap1_DAB9.1
NSSH10_hap1_DBB1.1
BS2_hap1_DBB2.1
BS2_hap2_DBB2.2
BS1_hap1_DBB3.1
BS4_hap1_DBB4.1
BS1_hap2_DBB4.2
BS2_hap1_DBB6.1
CS5_hap1_DBB8.1
American_shad_MHCIIB
Zebrafish_DAB
Zebrafish_DCB
Zebrafish_DGB
Common_carp_DAB
Atlantic_salmon_DAB
Rainbow_Trout_DAB
Rainbow_Trout_DBB
Arctic_char_DAB
Japanese_medaka_DAB
Japanese_medaka_DCB
Japanese_medaka_DFB
Stickleback_DAB
Stickleback_DBB
Nile_tilapia_DAB
Fugu_MHCIIB
Tetraodon_MHCIIB
Japanese_medaka_DDB1
Japanese_medaka_DEB
Atlantic_salmon_DBB
Atlantic_salmon_DCB
Atlantic_salmon_DEB
Spotted_gar_MHCIIB1
Spotted_gar_MHCIIB2
Dabrys_sturgeon_DAB
Nurse_shark_MHCIIB
African_clawed_frog_DMB
Chicken_DMB
Brown_rat_DMb
Mouse_H2-Mb1
Human_DMB
Brown_rat_DOb
Mouse_H2-Ob
Human_DOB
African_clawed_frog_DAB
Chicken_BLB
Brown_rat_Db1
Brown_rat_Bb
Mouse_H2-Ab1
Mouse_H2-Eb1
Human_DRB1
Human_DQB1
Human_DPB1

**Supplementary Fig. 4.** Amino acid sequence alignments of representative Atlantic herring sequences alongside sequences from fish and tetrapods, shown for the **a**, α1 domain and **b**, β1 domain. Sequence names are color-coded as in Extended Data Fig. 1: green for teleost DA/DB lineages, red for Atlantic herring DA lineage, blue for Atlantic herring DB lineage, gray for teleost DE lineage, yellow for tetrapod DM lineage, purple for tetrapod DO lineage, and pink for classical tetrapod sequences. Amino acid positions highlighted in green correspond to teleost-specific conserved cysteine residues in *DAA* genes, and fully conserved cysteine residues in *DAB* genes, that form disulfide bonds. Highly conserved residues involved in hydrogen bonding with the backbone of antigenic peptide are shaded in light red. Residues highlighted in yellow mark a four-amino-acid insertion (CTDC) in herring *DAA* sequences, which forms an additional disulfide bridge. Dashes represent gaps introduced for alignment. A chemistry-based coloring scheme is used to indicate amino acid chemical properties. The scale above the alignment indicates amino acid positions, beginning with the first residue of exon 2 in the full-length Atlantic herring protein. The black and red squares below the plots represent human[18] and chicken PBRs[19], respectively.

**a**

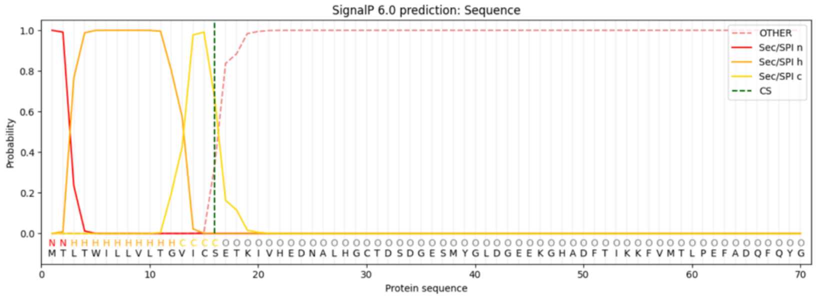


CS4_hap1_DAA2.1
Sequence prediction:
Signal peptide (Sec/SPI)
Cleavage site between pos. 16 and 17
Probability: 0.67

**b**

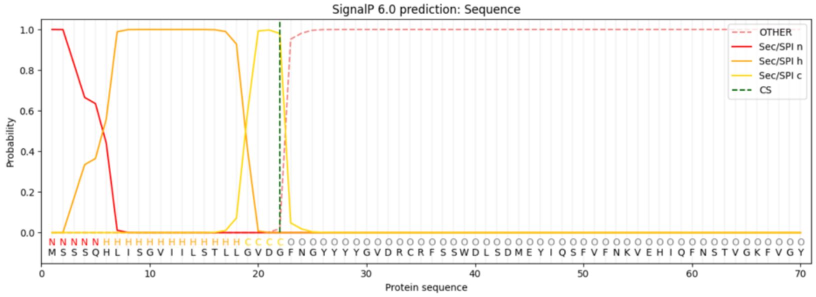


CS4_hap1_DAB2.1
Sequence prediction:
Signal peptide (Sec/SPI)
Cleavage site between pos. 22 and 23
Probability: 0.98

**c**

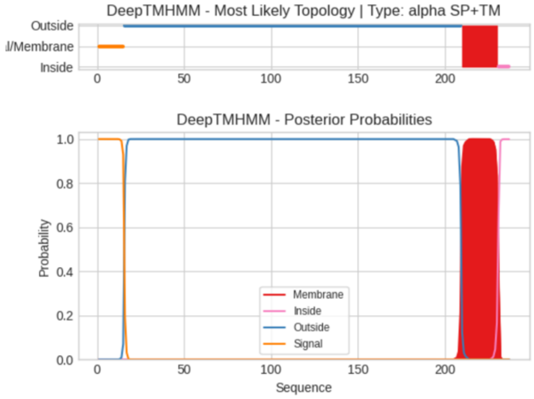


CS4_hap1_DAA2.1
Signal: 1 – 15
Outside: 16 – 209
TMhelix: 210 – 230
Inside: 231 – 237

**d**

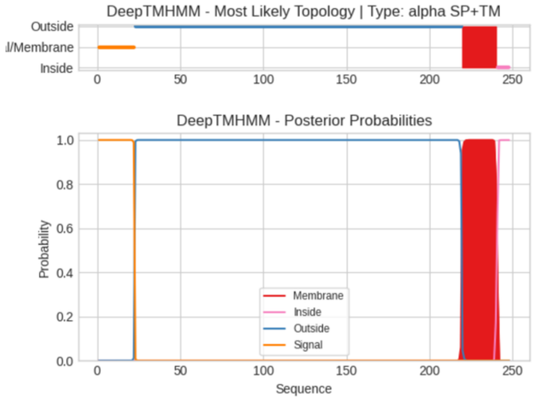


CS4_hap1_DAB2.1
Signal: 1 – 22
Outside: 23 – 219
TMhelix: 220 – 240
Inside: 241 – 248

**Supplementary Fig. 5.** Metrics of predicted MHC class II protein model formed by *CS4_hap1_DAA2.1* (alpha chain) and *CS4_hap1_DAB2.1* (beta chain). **a, b** Results from SignalP 6.0 signal peptide prediction of the alpha chain and beta chain, respectively. **c** and **d**, Results from DeepTMHMM transmembrane region prediction of the alpha chain and beta chain, respectively.

```
Reference_DAA2.1    IVHVDINLRG CSD----SDG EHMYALDGEE MGHADFTKGK FVMTMPEFAD PYTYEDSVYE FAVAEQKLCK QNMQIFTEAY KSPAEAE
BS1_hap1_DAA2.1     ..Q..TY.S. .T.CTDCT.. .E.AG..... A......... V...L..... ..K.VKGRFV ...L..QN.. H.LEVYIN.. .......
BS1_hap2_DAA2.1     ..Q..TY.S. .T.CTDCT.. .E.AG..... A......... V...L..... ..K.VKGRFV ...L..QN.. H.LEVYIN.. .......
BS3_hap2_DAA2.1     ..Q..TY.S. .T.CTDCT.. .E.AG..... A......... V...L..... ..K.VKGRFV ...L..QN.. H.LEVYIN.. .......
BS4_hap1_DAA2.1     ..Q..TY.S. .T.CTDCT.. .E.AG..... A......... V...L..... ..K.VKGRFV ...L..QN.. H.LEVYIN.. .......
BS5_hap1_DAA2.1     ..Q..TY.S. .T.CTDCT.. .E.AG..... A......... V...L..... ..K.VKGRFV ...L..QN.. H.LEVYIN.. .......
BS2_hap1_DAA2.2     ..Q..TY.S. .T.CTDCT.. .E.AG..... A......... V...L..... ..K.VKGRFV ...L..QN.. H.LEVYIN.. .......
BS5_hap2_DAA2.2     ..Q..TY.S. .T.CTDCT.. .E.AG..... A......... V...L..... ..K.VKGRFV ...L..QN.. H.LEVYIN.. .......
CS2_hap1_DAA2.2     ..Q..TY.S. .T.CTDCT.. .E.AG..... A......... V...L..... ..K.VKGRFV ...L..QN.. H.LEVYIN.. .......
CS2_hap2_DAA2.2     ..Q..TY.S. .T.CTDCT.. .E.AG..... A......... V...L..... ..K.VKGRFV ...L..QN.. H.LEVYIN.. .......
CS5_hap2_DAA2.2     ..Q..TY.S. .T.CTDCT.. .E.AG..... A......... V...L..... ..K.VKGRFV ...L..QN.. H.LEVYIN.. .......
CS8_hap1_DAA2.2     ..Q..TY.S. .T.CTDCT.. .E.AG..... A......... V...L..... ..K.VKGRFV ...L..QN.. H.LEVYIN.. .......
NSSH2_hap2_DAA2.2   ..Q..TY.S. .T.CTDCT.. .E.AG..... A......... V...L..... ..K.VKGRFV ...L..QN.. H.LEVYIN.. .......
```

**Supplementary Fig. 6.** Amino acid alignment of α1 domain sequences with an insertion of 'CTDC' domain, highlighted in a black box.

**a**

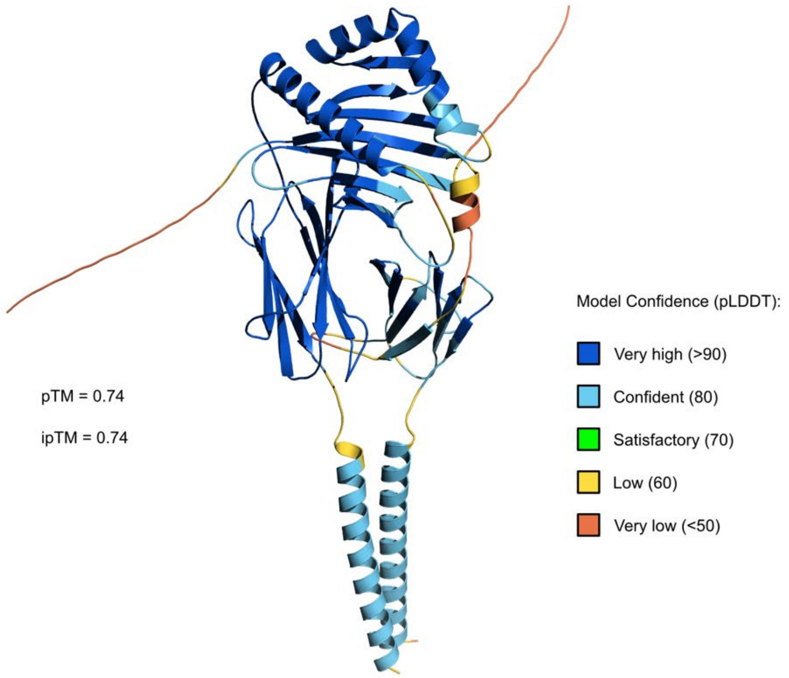

pTM = 0.74
ipTM = 0.74
Model Confidence (pLDDT):
Very high (>90)
Confident (80)
Satisfactory (70)
Low (60)
Very low (<50)


**b**

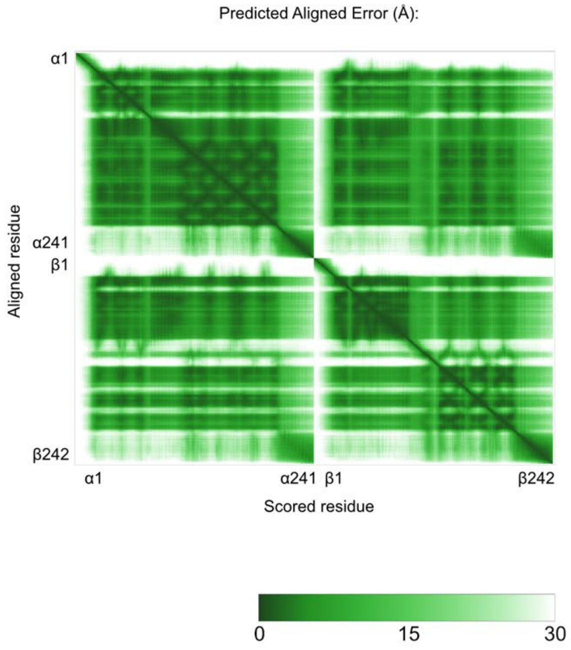

Predicted Aligned Error (Å):
α1
α241
β1
β242
Aligned residue
Scored residue
0
15
30

**Supplementary Fig. 7.** Predicted protein structure of the MHC class II heterodimer formed by *BS1_hap1_DAA2.1* (alpha chain) and *BS1_hap1_DAB2.1* (beta chain). **a** Model colored with the predicted local distance difference test values (pLDDT) of the alpha and beta chains. The predicted template modeling score (pTM) and interface predicted template modeling score (ipTM) are annotated. **b** Predicted aligned error (PAE) of the model. Amino acid residue number of alpha and beta chains are annotated by the axes of the plot.

a

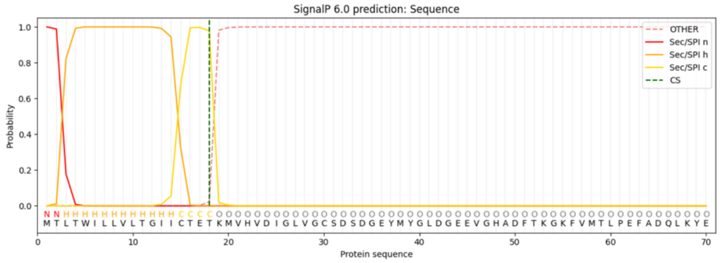


BS1_hap1_DAA2.1
Sequence prediction:
Signal peptide (Sec/SPI)
Cleavage site between pos.
18 and 19
Probability: 0.98

b

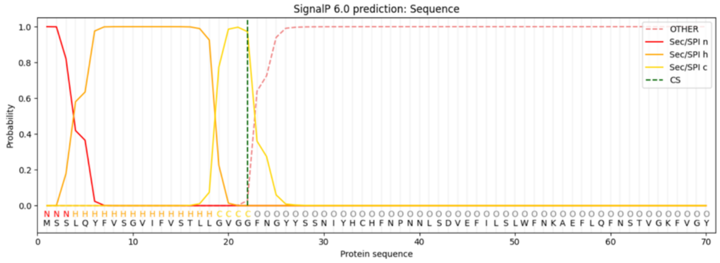


BS1_hap1_DAB2.1
Sequence prediction:
Signal peptide (Sec/SPI)
Cleavage site between pos.
22 and 23
Probability: 0.97

c

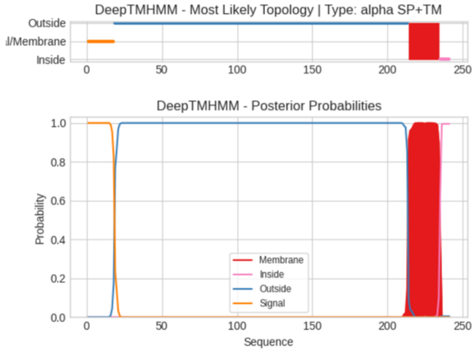


BS1_hap1_DAA2.1
Signal: 1 – 18
Outside: 19 – 213
TMhelix: 214 – 234
Inside: 235 – 241

d

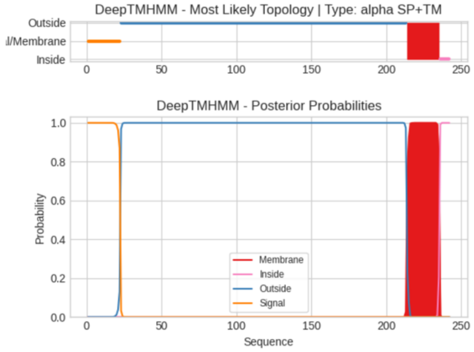


BS1_hap1_DAB2.1
Signal: 1 – 22
Outside: 23 – 213
TMhelix: 214 – 235
Inside: 236 – 242

**Supplementary Fig. 8.** Metrics of predicted MHC class II protein model *BS1_hap1_DAA2.1* (alpha chain) and *BS1_hap1_DAB2.1* (beta chain). **a, b** Results from SignalP 6.0 signal peptide prediction of the alpha chain and beta chain, respectively. **c, d** Results from DeepTMHMM transmembrane region prediction of the alpha chain and beta chain, respectively.

**Supplementary Table 1.** Non-synonymous substitutions (*dN*) and synonymous substitutions (*dS*) values computed by codon selection test in MEGA for exon2 and exon3 sequences of alpha and beta genes from DA and DB lineages along with their respective standard errors (S.E.).

| Gene_exon | No. sequences | *dN* ± S.E. | *dS* ± S.E. | *dN/dS* | P value |
|---|---|---|---|---|---|
| DAA1.1_Exon2 | 29 | 0.123 ± 0.019 | 0.095 ± 0.022 | 1.2891 | 0.13 |
| DAA1.2_Exon2 | 28 | 0.132 ± 0.020 | 0.084 ± 0.020 | 1.5652 | 0.02 |
| DAA1.3_Exon2 | 8 | 0.047 ± 0.009 | 0.027 ± 0.013 | 1.7562 | 0.09 |
| DAA2.1_Exon2 | 24 | 0.113 ± 0.019 | 0.040 ± 0.014 | 2.8140 | 0.0004 |
| DAA2.2_Exon2 | 24 | 0.141 ± 0.018 | 0.083 ± 0.019 | 1.6954 | 0.02 |
| DAA2.3_Exon2 | 6 | 0.056 ± 0.014 | 0.044 ± 0.020 | 1.2643 | 0.30 |
| DAA4.1_Exon2 | 28 | 0.107 ± 0.018 | 0.065 ± 0.019 | 1.6358 | 0.02 |
| DAA4.2_Exon2 | 5 | 0.146 ± 0.023 | 0.069 ± 0.022 | 2.1099 | 0.003 |
| DAA5.1_Exon2 | 28 | 0.001 ± 0.001 | 0 ± 0 | N/A | 0.16 |
| DAA7.1_Exon2 | 8 | 0.048 ± 0.011 | 0.033 ± 0.011 | 1.4846 | 0.15 |
| DAB1.1_Exon2 | 29 | 0.174 ± 0.021 | 0.178 ± 0.026 | 0.9787 | 1 |
| DAB1.2_Exon2 | 29 | 0.186 ± 0.022 | 0.128 ± 0.019 | 1.4553 | 0.005 |
| DAB1.3_Exon2 | 9 | 0.065 ± 0.013 | 0.046 ± 0.017 | 1.3950 | 0.18 |
| DAB2.1_Exon2 | 29 | 0.148 ± 0.018 | 0.071 ± 0.017 | 2.0785 | 0.0003 |
| DAB2.2_Exon2 | 25 | 0.177 ± 0.020 | 0.101 ± 0.021 | 1.7525 | 0.001 |
| DAB2.3_Exon2 | 6 | 0.116 ± 0.020 | 0.037 ± 0.015 | 3.1596 | 0.0006 |
| DAB4.1_Exon2 | 27 | 0.141 ± 0.018 | 0.082 ± 0.020 | 1.7249 | 0.006 |
| DAB4.2_Exon2 | 5 | 0.159 ± 0.021 | 0.076 ± 0.022 | 2.0862 | 0.002 |
| DAB5.1a_Exon2 | 29 | 0.040 ± 0.009 | 0.030 ± 0.013 | 1.3303 | 0.26 |
| DAB5.1b_Exon2 | 19 | 0.017 ± 0.003 | 0.011 ± 0.006 | 1.4593 | 0.19 |
| DAB7.1_Exon2 | 6 | 0.030 ± 0.008 | 0.055 ± 0.023 | 0.5551 | 1 |
| DBA3.1_Exon2 | 29 | 0.003 ± 0.003 | 0.001 ± 0.001 | 2.4259 | 0.28 |
| DBA4.1_Exon2 | 16 | 0.004 ± 0.003 | 0.007 ± 0.007 | 0.5942 | 1 |
| DBB2.1_Exon2 | 27 | 0 ± 0 | 0 ± 0 | N/A | N/A |
| DBB3.1_Exon2 | 28 | 0.001 ± 0.001 | 0.006 ± 0.003 | 0.1740 | 1 |
| DBB4.1_Exon2 | 26 | 0 ± 0 | 0.012 ± 0.005 | 0.0296 | 1 |
| DBB6.1_Exon2 | 20 | 0.004 ± 0.002 | 0.009 ± 0.006 | 0.4579 | 1 |
| DAA1.1_Exon3 | 29 | 0.054 ± 0.011 | 0.090 ± 0.019 | 0.6001 | 1 |
| DAA1.2_Exon3 | 28 | 0.052 ± 0.011 | 0.097 ± 0.019 | 0.5374 | 1 |
| DAA1.3_Exon3 | 8 | 0.026 ± 0.008 | 0.037 ± 0.013 | 0.7009 | 1 |
| DAA2.1_Exon3 | 24 | 0.036 ± 0.008 | 0.066 ± 0.017 | 0.5395 | 1 |
| DAA2.2_Exon3 | 24 | 0.030 ± 0.007 | 0.063 ± 0.015 | 0.4713 | 1 |
| DAA2.3_Exon3 | 6 | 0.015 ± 0.006 | 0.053 ± 0.019 | 0.2892 | 1 |
| DAA4.1_Exon3 | 28 | 0.048 ± 0.009 | 0.081 ± 0.020 | 0.5884 | 1 |

| Gene_exon | No. sequences | *dN* ± S.E. | *dS* ± S.E. | *dN/dS* | P value |
|---|---|---|---|---|---|
| DAA4.2_Exon3 | 5 | 0.043 ± 0.010 | 0.099 ± 0.026 | 0.4342 | 1 |
| DAA5.1_Exon3 | 28 | 0 ± 0 | 0 ± 0 | N/A | N/A |
| DAA7.1_Exon3 | 8 | 0.021 ± 0.006 | 0.042 ± 0.019 | 0.4899 | 1 |
| DAB1.1_Exon3 | 29 | 0.059 ± 0.011 | 0.119 ± 0.023 | 0.4917 | 1 |
| DAB1.2_Exon3 | 29 | 0.052 ± 0.011 | 0.096 ± 0.020 | 0.5427 | 1 |
| DAB1.3_Exon3 | 9 | 0.051 ± 0.011 | 0.090 ± 0.026 | 0.5639 | 1 |
| DAB2.1_Exon3 | 29 | 0.019 ± 0.006 | 0.064 ± 0.018 | 0.3028 | 1 |
| DAB2.2_Exon3 | 25 | 0.037 ± 0.008 | 0.089 ± 0.020 | 0.4119 | 1 |
| DAB2.3_Exon3 | 6 | 0.025 ± 0.008 | 0.070 ± 0.021 | 0.3524 | 1 |
| DAB4.1_Exon3 | 27 | 0.024 ± 0.007 | 0.067 ± 0.018 | 0.3556 | 1 |
| DAB4.2_Exon3 | 5 | 0.021 ± 0.008 | 0.069 ± 0.023 | 0.3003 | 1 |
| DAB5.1a_Exon3 | 29 | 0.004 ± 0.002 | 0.041 ± 0.015 | 0.0871 | 1 |
| DAB5.1b_Exon3 | 19 | 0.002 ± 0.001 | 0.033 ± 0.013 | 0.0715 | 1 |
| DAB7.1_Exon3 | 6 | 0.039 ± 0.008 | 0.048 ± 0.019 | 0.8087 | 1 |
| DBA3.1_Exon3 | 29 | 0.001 ± 0.000 | 0.002 ± 0.001 | 0.3070 | 1 |
| DBA4.1_Exon3 | 16 | 0.007 ± 0.003 | 0.031 ± 0.013 | 0.2407 | 1 |
| DBB2.1_Exon3 | 27 | 0.009 ± 0.003 | 0.042 ± 0.016 | 0.2063 | 1 |
| DBB3.1_Exon3 | 28 | 0.001 ± 0.000 | 0.007 ± 0.003 | 0.0959 | 1 |
| DBB4.1_Exon3 | 26 | 0.009 ± 0.004 | 0.025 ± 0.010 | 0.3459 | 1 |
| DBB6.1_Exon3 | 20 | 0.019 ± 0.006 | 0.025 ± 0.010 | 0.7673 | 1 |

**Supplementary Table 2.** Summary of studies on MHC II genes on teleost

| **Species** | **Approach** | **MHC class II gene/exon** | **Ref.** |
|---|---|---|---|
| Cold water adapted icefish (*Chionodraco hamatus*) | A total of 54 samples were studied, of which cloning and PCR was performed on 10 and Ion Torrent sequencing was performed on 44, resulting in 41 sequences in total with maximum 9 sequences in a single individual. | DAB, exon2 | 28 |
| European eel (*Anguilla anguilla*) | A total of four samples were studied using PCR for entire gene, resulting in a total of 14 alpha and 16 beta sequences. | DAA and DAB, entire gene | 29 |
| Tongue sole (*Cynoglossus semilaevis*) | A total of six samples were studied where PCR was performed on exon2, intron2, and exon3, resulting in six alpha sequences from DA lineage and seven from DB lineage. | DAA and DBA, exon2, intron2, and exon3 | 30 |
| Stone flounder (*Kareius bicoloratus*) and Japanese flounder (*Paralichthys olivaceus*) | A total of eight samples were studied where PCR was performed on exons 1-4 (total 6 exons), resulting in 28 alleles. | Beta | 31 |
| Nile tilapia (*Oreochromis niloticus*) | A total of 60,000 BAC clones were used to build genomic organization, and genes were amplified by PCR, resulting in 9 alpha and 15 beta genes across six regions, DA through DF. | Alpha and beta genes | 32 |
| Midas cichlid (*Amphilophus citrinellus)* | A total of 13 samples were studied resulting in 69 alleles, with upto 25 alleles per individual. | Beta | 33 |
| Species of *O. niloticus* (old world cichlids from East African Great Lakes) | Species from five families were studied using PCR resulting in 17 polymorphic loci. | Beta | 34 |
| Nicaraguan Midas cichlid (*Amphilophus spp.*) | A total of 287 samples from six sympatric assemblages were studied, resulting in 152 unique exon2 alleles. | Beta | 35 |
| Cichlid genera | Sequences were retrieved from public datasets, resulting in 155 exon2 sequences belonging to 28 cichlid taxa and 40 exon3 sequences from 17 species. | Beta, exon 2 and exon 3 | 36 |

| **Species** | **Approach** | **MHC class II gene/exon** | **Ref.** |
|---|---|---|---|
| European bitterling (*Rhodeus amarus*) | A total of 221 samples were used, resulting in 36 sequences with 1-4 allelic variants per individual. | DAB1 | 37 |
| Rhodeus sinensis *(bitterling species)* | A total of 50 samples were used, resulting in 140 allelic sequences. | DAB | 38 |
| Sichuan taimen *(Hucho bleekeri)* | A total of 7 samples were used, resulting in 16 alpha and 16 beta sequences | Alpha and beta genes | 39 |

**Footnote.** 28. Hofmann, M. J., Bracamonte, S. E., Eizaguirre, C. & Barluenga, M. Molecular characterization of MHC class IIB genes of sympatric Neotropical cichlids. *BMC Genet.* **18**, 1–17 (2017). 29. Gerdol, M. *et al.* Molecular and Structural Characterization of MHC Class II β Genes Reveals High Diversity in the Cold-Adapted Icefish Chionodraco hamatus. *Sci. Rep.* **9**, 1–14 (2019). 30. Jeon, H. B., Won, H. & Suk, H. Y. Polymorphism of MHC class IIB in an acheilognathid species, Rhodeus sinensis shaped by historical selection and recombination. *BMC Genet.* **20**, 74 (2019). 31. Chen, Y. *et al.* Characterization, expression, and polymorphism of MHC II α and MHC II β in Sichuan taimen (Hucho bleekeri). *Comp. Biochem. Physiol. Part A Mol. Integr. Physiol.* **299**, 111767 (2025). 32. Thesis Seraina, M. S., Bracamonte, E. & Bracamonte, S. E. *Characterization and evolution of MHC II genes in the European eel (Anguilla anguilla)* (2013). 33. Li, C., Jiang, J., Zhang, Q. & Wang, X. Duplicated major histocompatibility complex class II genes in the tongue sole (Cynoglossus semilaevis). *Int. J. Immunogenet.* **45**, 210–224 (2018). 34. Jiang, J., Li, C., Zhang, Q. & Wang, X. Locus Number Estimation of MHC Class II B in Stone Flounder and Japanese Flounder. *Int. J. Mol. Sci. 2015, Vol. 16, Pages 6000-6017* **16**, 6000–6017 (2015). 35. Sato, A., Dongak, R., Hao, L., Shintani, S. & Sato, T. Organization of Mhc class II A and B genes in the tilapiine fish Oreochromis. *Immunogenetics* **64**, 679–690 (2012). 36. Málaga-Trillo, E. *et al.* Linkage Relationships and Haplotype Polymorphism Among Cichlid Mhc Class II B Loci. *Genetics* **149**, 1527–1537 (1998). 37. Bracamonte, S. E., Hofmann, M. J., Lozano-Martín, C., Eizaguirre, C. & Barluenga, M. Divergent and non-parallel evolution of MHC IIB in the Neotropical Midas cichlid species complex. *BMC Ecol. Evol.* **22**, 1–17 (2022). 38. Lozano-Martín, C., Bracamonte, S. E. & Barluenga, M. Evolution of MHC IIB Diversity Across Cichlid Fish Radiations. *Genome Biol. Evol.* **15**, (2023). 39. Talarico, L., Bryjová, A., Čížková, D., Douda, K. & Reichard, M. Individual copy number variation and extensive diversity between major MHC-DAB1 allelic lineages in the European bitterling. *Immunogenetics* **74**, 497–505 (2022).

**Supplementary Table 3**. Summary of null allele occurrences and mutations

| Gene | Number of assemblies | Description of a mutation |
|---|---|---|
| *DAA1.2* | 1 | A 1-bp insertion (C) at the beginning of exon 3 |
| *DAA2.1* | 5 | Only exon 1 and 2 were found, exons 3 and 4 are seemingly missing |
| *DAA2.2* | 2 | A 1-bp insertion (C) in the beginning of exon2 leading to stop codon in exon2; Absence of exon 1 in addition to deletion of first base pair of exon 2 which resulting in a frameshift mutation |
| *DAB2.2* | 1 | A 2-bp deletion in exon 1 (third codon) leading to frameshift mutation. |
| *DAA2.3* | 2 | A 1-bp insertion (C) at the beginning of exon 3 |
| *DAB2.3* | 2 | A 2-bp deletion in exon 1 |
| *DAA5.1* | 1 | A 106-bp deletion in exon 3 causing frameshift mutation. |
| *DAB5.1b* | 10 | An SNV in exon 2 caused a stop codon (nonsense mutation) |
| *DAB7.1* | 2 | Part of exon 1 is missing; Stop gain mutation |
| *DBB1.1* | 1 | Absence of exon 1 |
| *DBA4.1* | 14 | A 5-bp deletion in exon 3; A 147-bp insertion in exon 3 |
| *DBA6.1* | 20 | Truncated one or more exons |
| *DBB1.1* | 19 | Truncated one or more exons |
| *DBB2.1* | 1 | A 7-bp deletion at exon 3 caused a frameshift |
| *DBB4.1* | 3 | A 1-bp insertion in exon 2; Truncated one or more exons |
| *DBB6.1* | 7 | A 22-bp deletion in Exon 3; A 2-bp deletion in exon 3; A 1-bp insertion in exon 3; A 442-bp insertion in exon 3; Truncated one or more exons |

**Supplementary Table 4.** Length (bp) of intron 1 in the *DAA1.3* gene from eight haplotypes

| Haplotype | Intron1 length bp) |
|---|---|
| BS3_hap1 | 20,902 |
| BS4_hap2 | 18,216 |
| BS5_hap1 | 21,149 |
| BS6_hap1 | 20,219 |
| CS2_hap2 | 17,809 |
| CS4_hap2 | 19,496 |
| CS7_hap1 | 20,217 |
| CS8_hap1 | 23,335 |

**Supplementary Table 5.** List of MHC class II α and β chain sequences from other species used in phylogenetic analyses, along with their corresponding accession numbers or gene IDs.

| Common name | Scientific name | gene lineage | Source DB | Accession number/Gene ID | |
|---|---|---|---|---|---|
| | | | | Alpha chain | Beta chain |
| American shad | *Alosa sapidissima* | DA | NCBI | MN364683 | MN364682 |
| Zebrafish | *Danio rerio* | DA | NCBI | NP_001004521 | NP_571551 |
| Zebrafish | *Danio rerio* | DC | NCBI | NP_001410940 | NP_001007036 |
| Zebrafish | *Danio rerio* | DG | NCBI | NP_001007206 | NP_001007207 |
| Common carp | *Cyprinus carpio* | DA | NCBI | CAA64707 | CAA64709 |
| Atlantic salmon | *Salmo salar* | DA | IPD | FISH08192 | FISH08214 |
| Atlantic salmon | *Salmo salar* | DB | NCBI | XP_013992518 | XP_013992517 |
| Atlantic salmon * | *Salmo salar* | DC | NCBI | DY704572 | KC316031 |
| Atlantic salmon | *Salmo salar* | DE | NCBI | XP_045569620 | XP_045569629 |
| Rainbow Trout | *Oncorhynchus mykiss* | DA | IPD | FISH08121 | FISH08122 |
| Rainbow Trout | *Oncorhynchus mykiss* | DB | NCBI | XP_036806345 | NP_001182463 |
| Arctic char | *Salvelinus alpinus* | DA | NCBI | ACI05079 | ACI05078 |
| Japanese medaka | *Oryzias latipes* | DA | NCBI | JQ743249 | JQ743258 |
| Japanese medaka | *Oryzias latipes* | DC | NCBI | JQ743250 | JQ743259 |
| Japanese medaka | *Oryzias latipes* | DD | NCBI | JQ743252 | JQ743261 |
| Japanese medaka | *Oryzias latipes* | DE | NCBI | JQ743255 | JQ743265 |
| Japanese medaka | *Oryzias latipes* | DF | NCBI | JQ743256 | JQ743266 |

| **Common name** | **Scientific name** | **gene lineage** | **Source DB** | **Accession number/Gene ID** | |
|---|---|---|---|---|---|
| | | | | **Alpha chain** | **Beta chain** |
| Three-spined stickleback | *Gasterosteus aculeatus* | DA | NCBI | AAU01917 | AAU01918 |
| Three-spined stickleback | *Gasterosteus aculeatus* | DB | NCBI | AAU01919 | AAU01920 |
| Nile tilapia | *Oreochromis niloticus* | DA | NCBI | NP_001269817 | QJD21088 |
| Fugu * | *Takifugu rubripes* | | Ensembl | ENSTRUG00000017444 | ENSTRUG00000017449 |
| Tetraodon * | *Tetraodon nigroviridis* | | Ensembl | ENSTNIG00000005593 | ENSTNIG00000005592 |
| Spotted gar * | *Lepisosteus oculatus* | | NCBI | JH591501 | JH591501 |
| Spotted gar * | *Lepisosteus oculatus* | | NCBI | JH591615 | JH591615 |
| Dabry's sturgeon | *Acipenser dabryanus* | DA | NCBI | MK923681 | MK923693 |
| Nurse shark | *Ginglymostoma cirratum* | | NCBI | AAA49311 | L20274 |
| African clawed frog | *Xenopus laevis* | DA | NCBI | AF454374 | BAA08759 |
| African clawed frog | *Xenopus laevis* | DM | NCBI | AAH61681 | ABB85336 |
| Chicken | *Gallus gallus* | BL | NCBI | AAR14673 | AAA48948 |
| Chicken | *Gallus gallus* | DM | NCBI | CAA18966 | CAA18967 |
| House mouse | *Mus musculus* | H2-A | NCBI | NP_034508 | NP_996988 |
| House mouse | *Mus musculus* | H2-E | NCBI | NP_034511 | NP_034512 |
| House mouse | *Mus musculus* | H2-M | NCBI | NP_001347459 | NP_034517 |
| House mouse | *Mus musculus* | H2-O | NCBI | NP_032232 | NP_034519 |
| Brown rat | *Rattus norvegicus* | RT1-B | NCBI | NP_001008831 | NP_001004084 |
| Brown rat | *Rattus norvegicus* | RT1-D | IPD | RT108424 | RT108427 |
| Brown rat | *Rattus norvegicus* | RT1-DM | NCBI | NP_942036 | NP_942035 |

| Common name | Scientific name | gene lineage | Source DB | Accession number/Gene ID | |
|---|---|---|---|---|---|
| | | | | **Alpha chain** | **Beta chain** |
| Brown rat | *Rattus norvegicus* | RT1-DO | NCBI | NP_898874 | NP_001008846 |
| Human | *Homo sapiens* | HLA-DR | IPD | HLA00662 | HLA00664 |
| Human | *Homo sapiens* | HLA-DQ | IPD | HLA00619 | HLA00622 |
| Human | *Homo sapiens* | HLA-DP | IPD | HLA00499 | HLA00514 |
| Human | *Homo sapiens* | HLA-DM | IPD | HLA00485 | HLA00489 |
| Human | *Homo sapiens* | HLA-DO | IPD | HLA00494 | HLA01098 |

* Retrieved from Dijkstra et al., 2013.
