## Supplementary Figure 9 for "Unheralded high MHC Class II polymorphism in the abundant Atlantic herring resolved by long-read sequencing"

**Supplementary Fig. 9.** Dotplots showing pairwise alignments of MHC class II haplotypes at Locus 2 from 28 phased haploid genome assemblies. Each page compares one assembly (y-axis) to all others (x-axes). The aligned sequence includes the full MHC class II region spanning the DA and DB lineage α- and β-chain genes, intergenic regions, and 20 kb of flanking sequence on each side. Red and blue lines indicate alignments in the same and opposite orientations, respectively. Gene positions are indicated by vertical and horizontal colored bars: green for DAA, blue for DAB, purple for DBA, and pink for DBB.

### CS2_hap1 vs. all assemblies − Locus 2

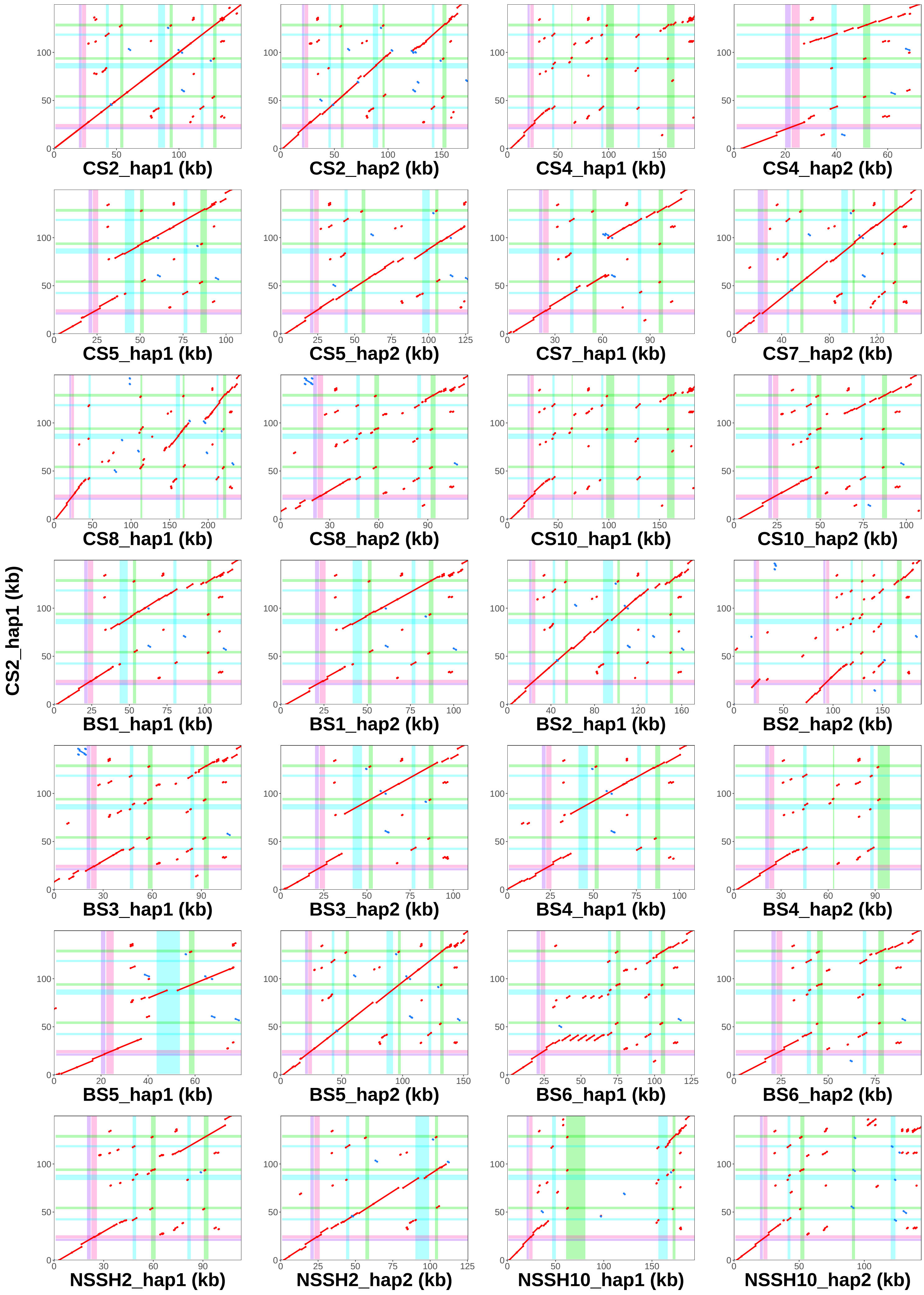

### CS2_hap2 vs. all assemblies − Locus 2

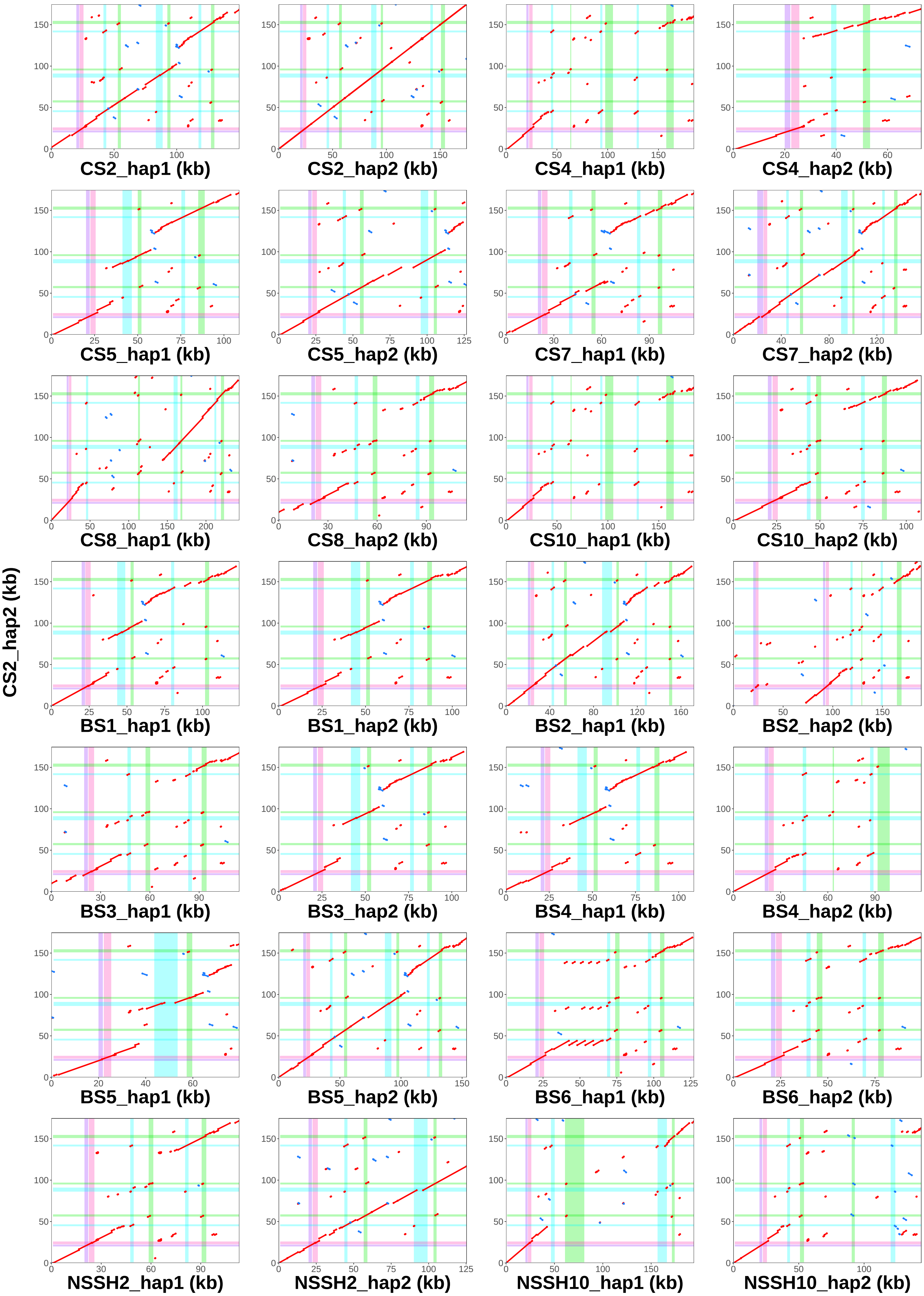

### CS4_hap1 vs. all assemblies − Locus 2

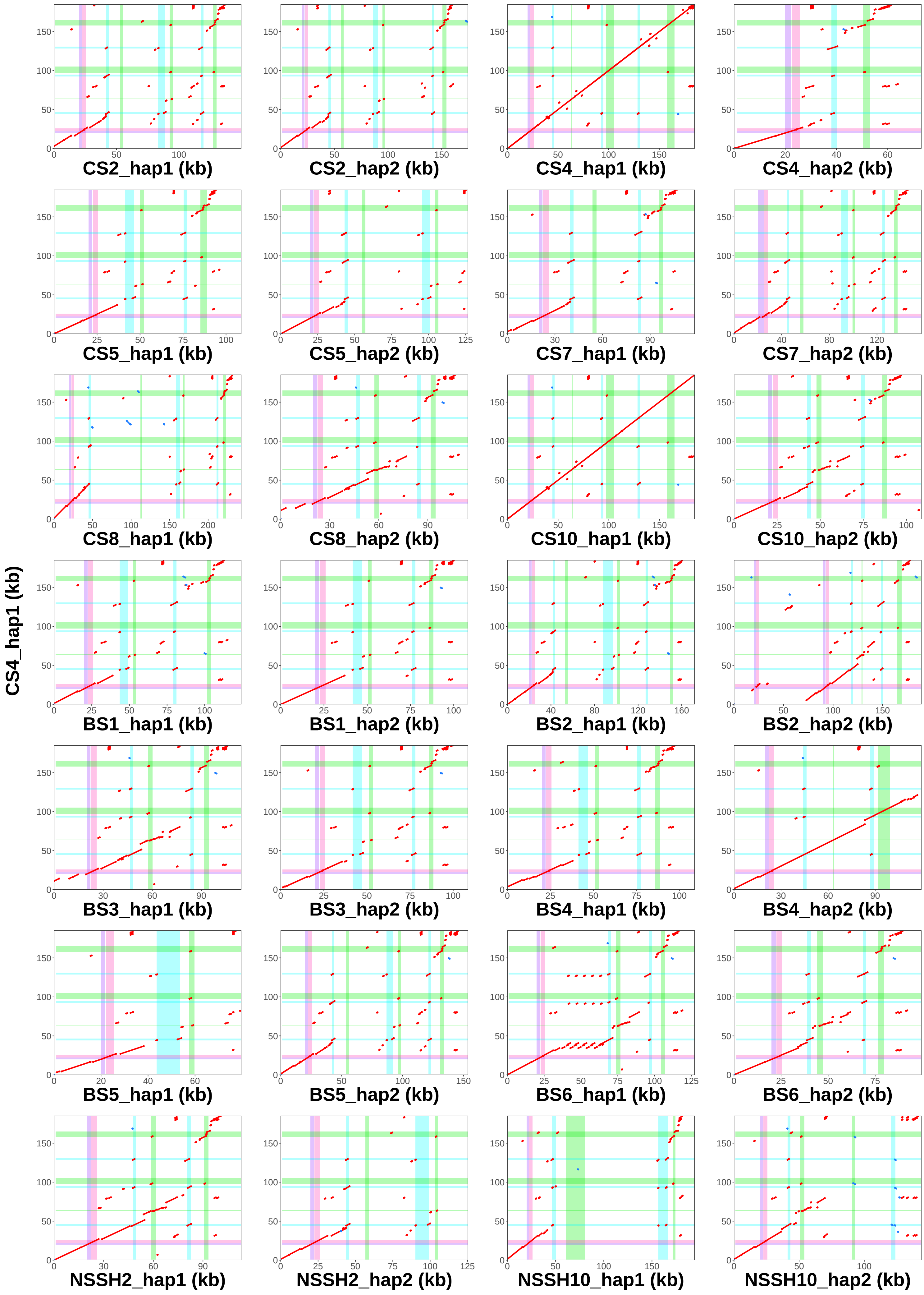

### CS4_hap2 vs. all assemblies − Locus 2

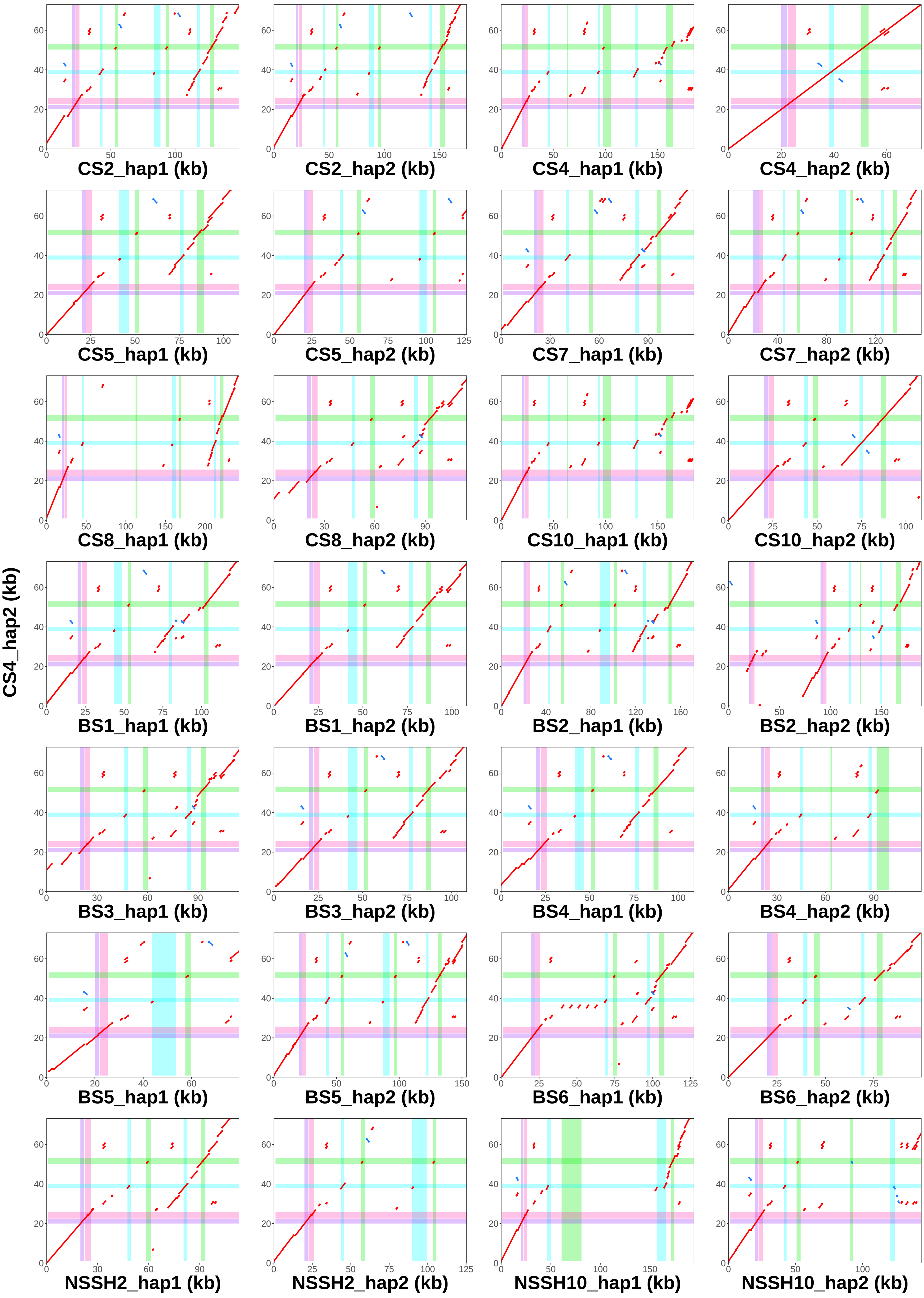

### CS5_hap1 vs. all assemblies − Locus 2

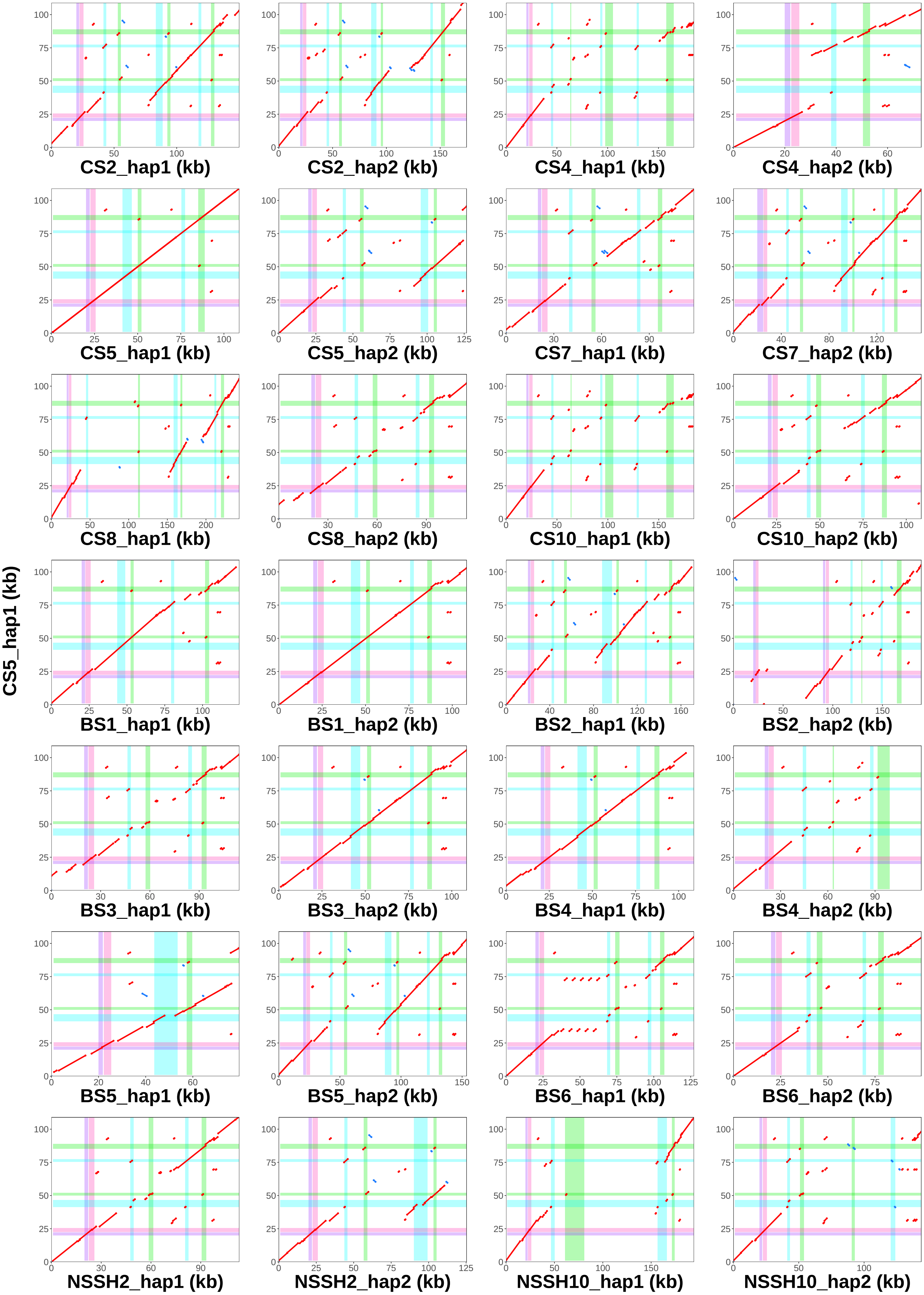

### CS5_hap2 vs. all assemblies − Locus 2

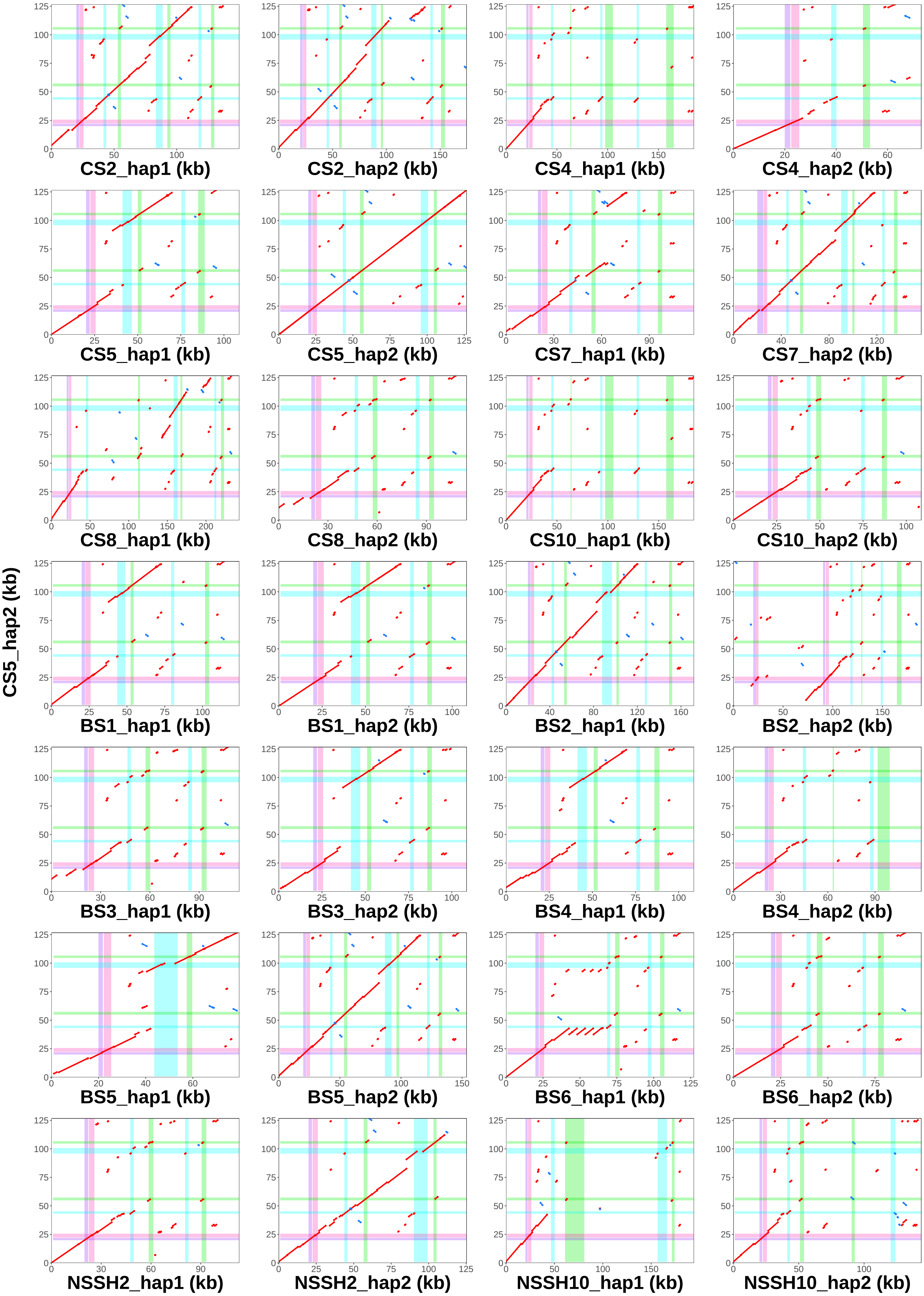

### CS7_hap1 vs. all assemblies − Locus 2

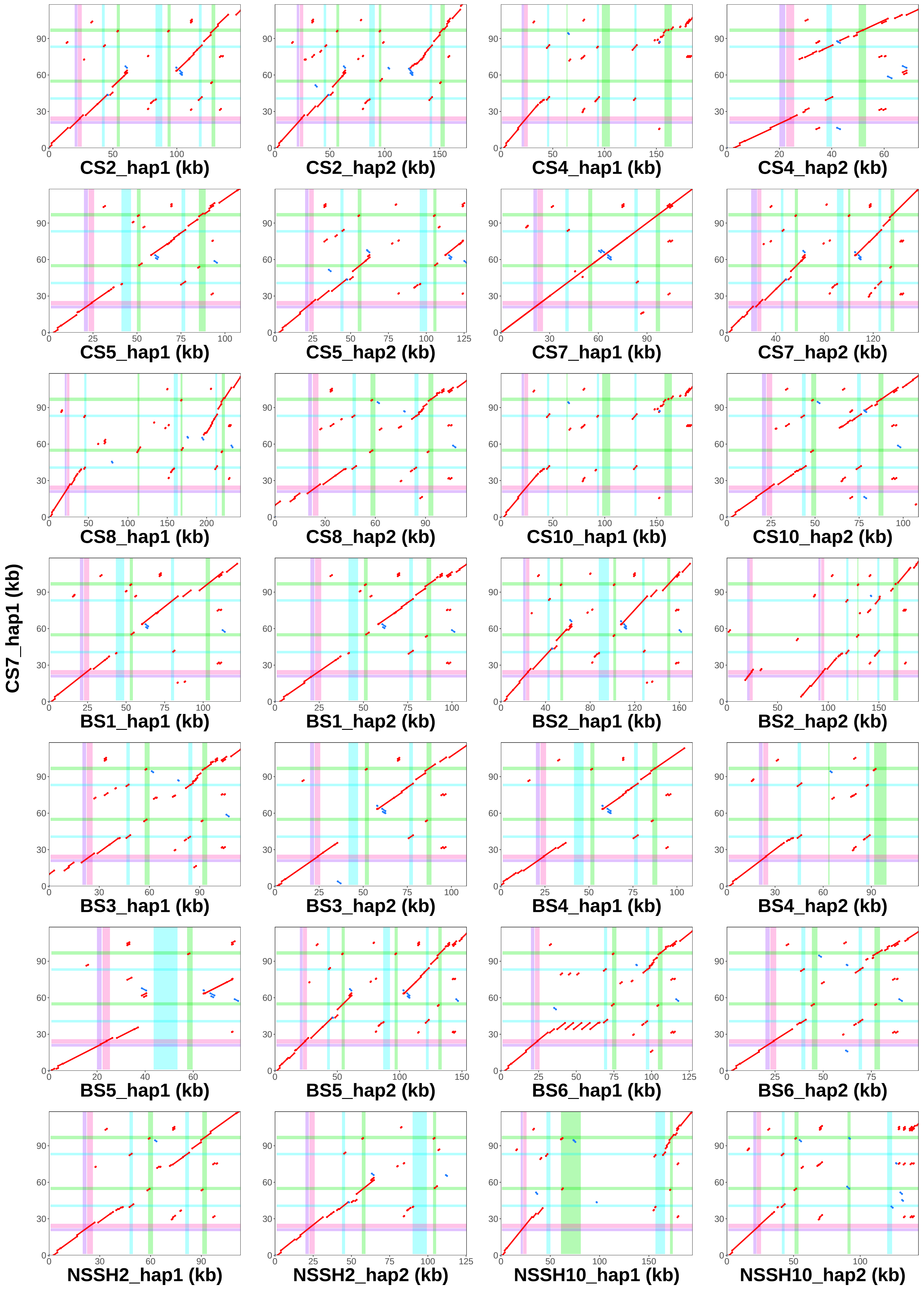

### CS7_hap2 vs. all assemblies − Locus 2

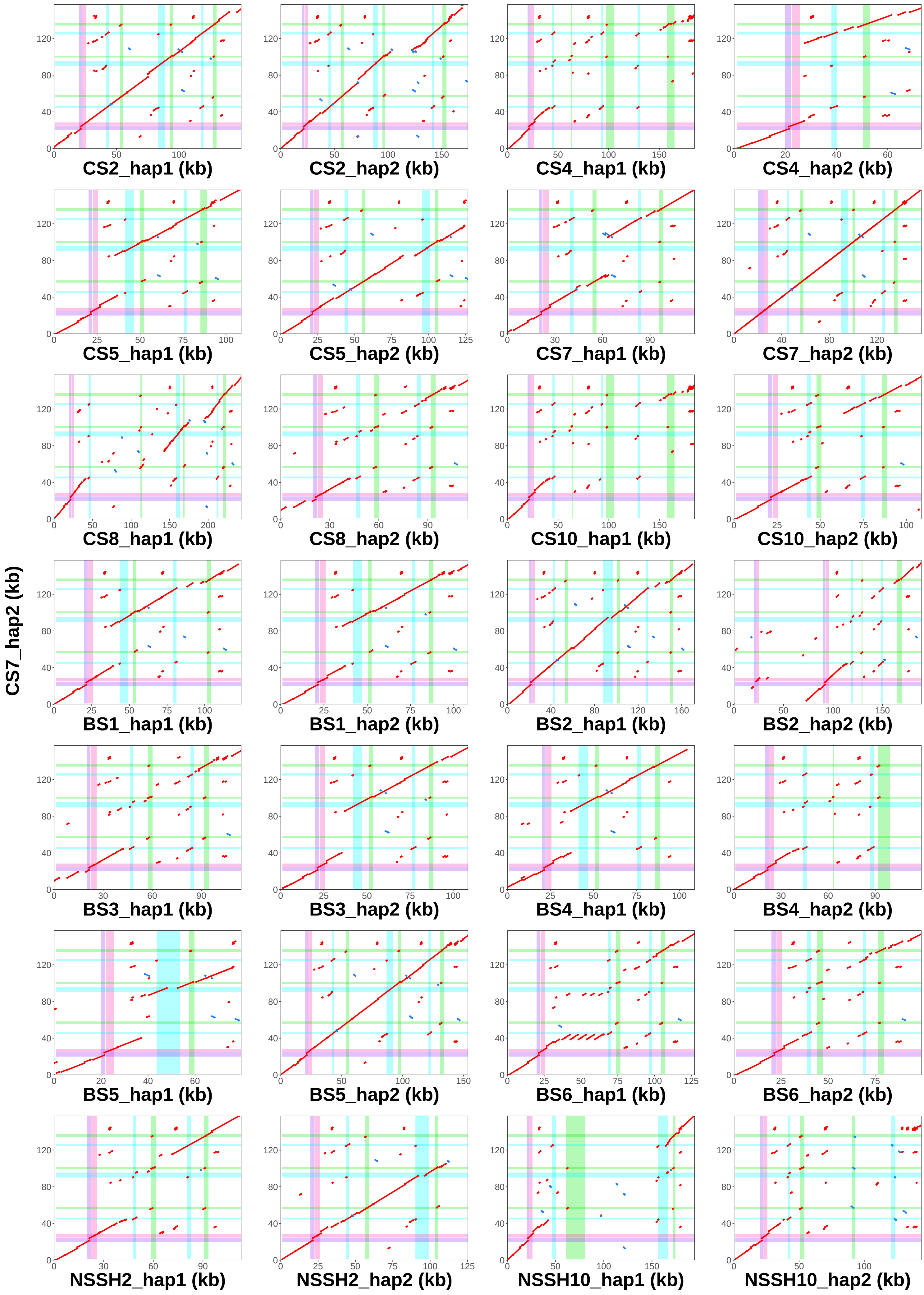

### CS8_hap1 vs. all assemblies − Locus 2

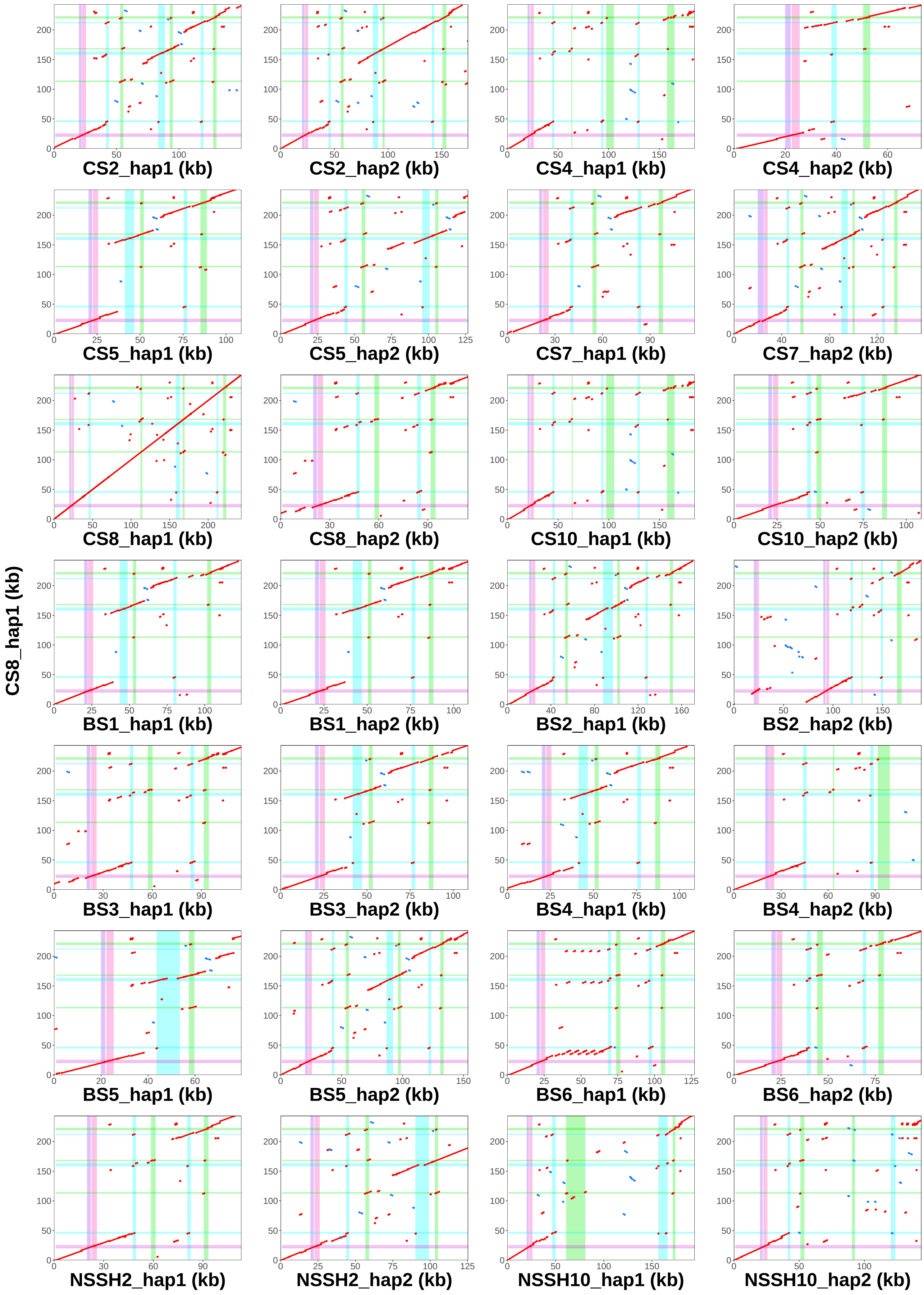

### CS8_hap2 vs. all assemblies − Locus 2

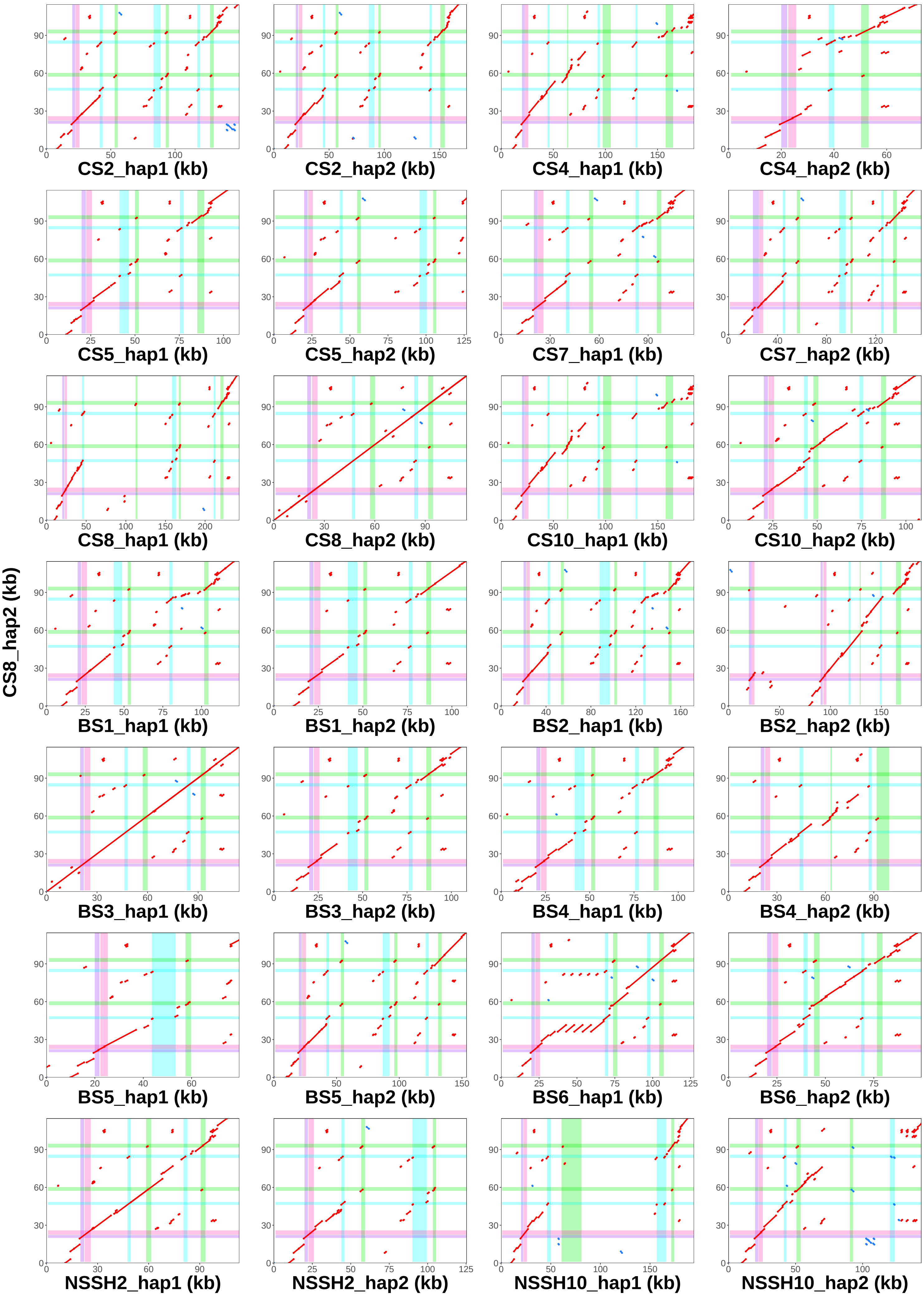

### CS10_hap1 vs. all assemblies − Locus 2

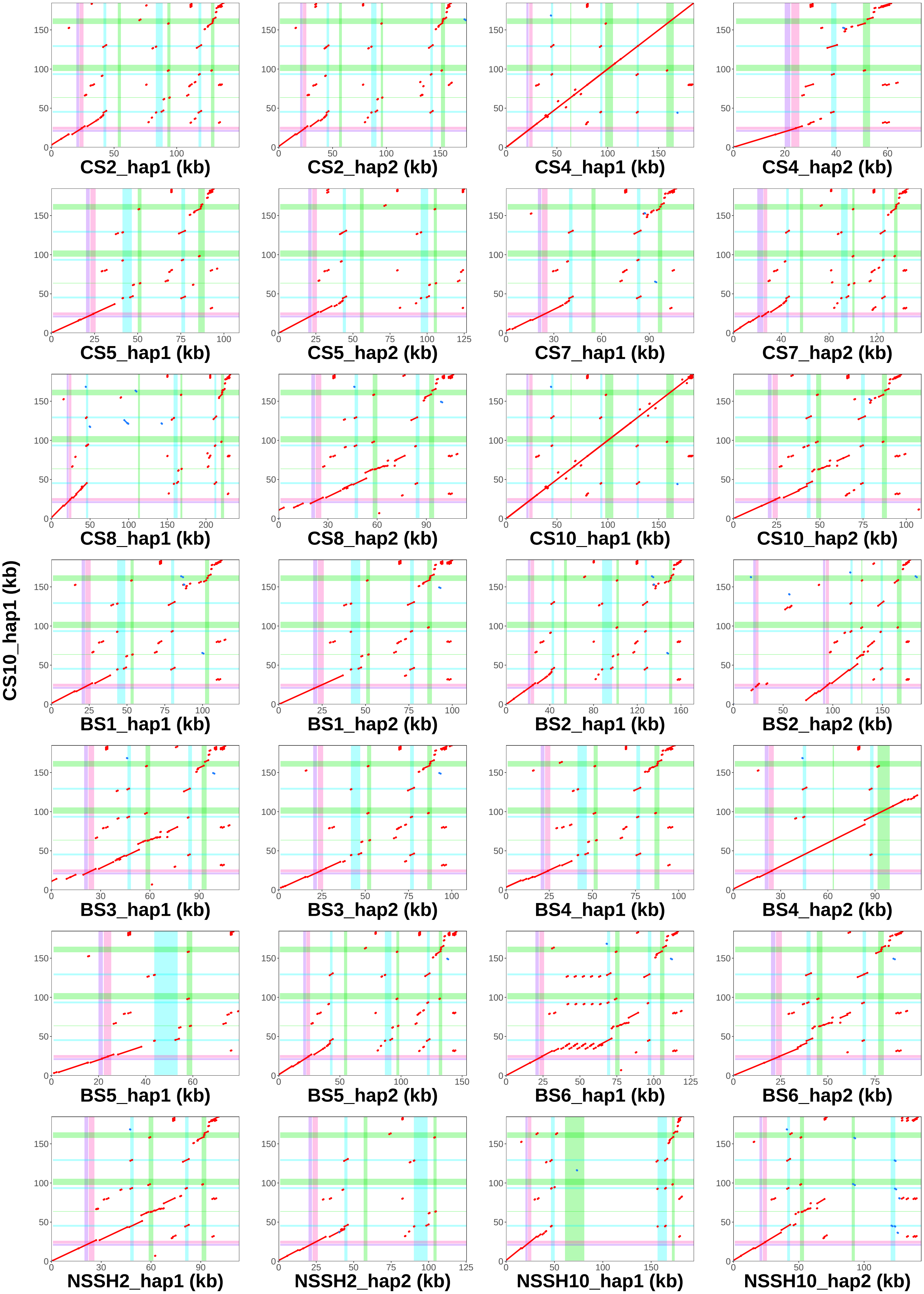

### CS10_hap2 vs. all assemblies − Locus 2

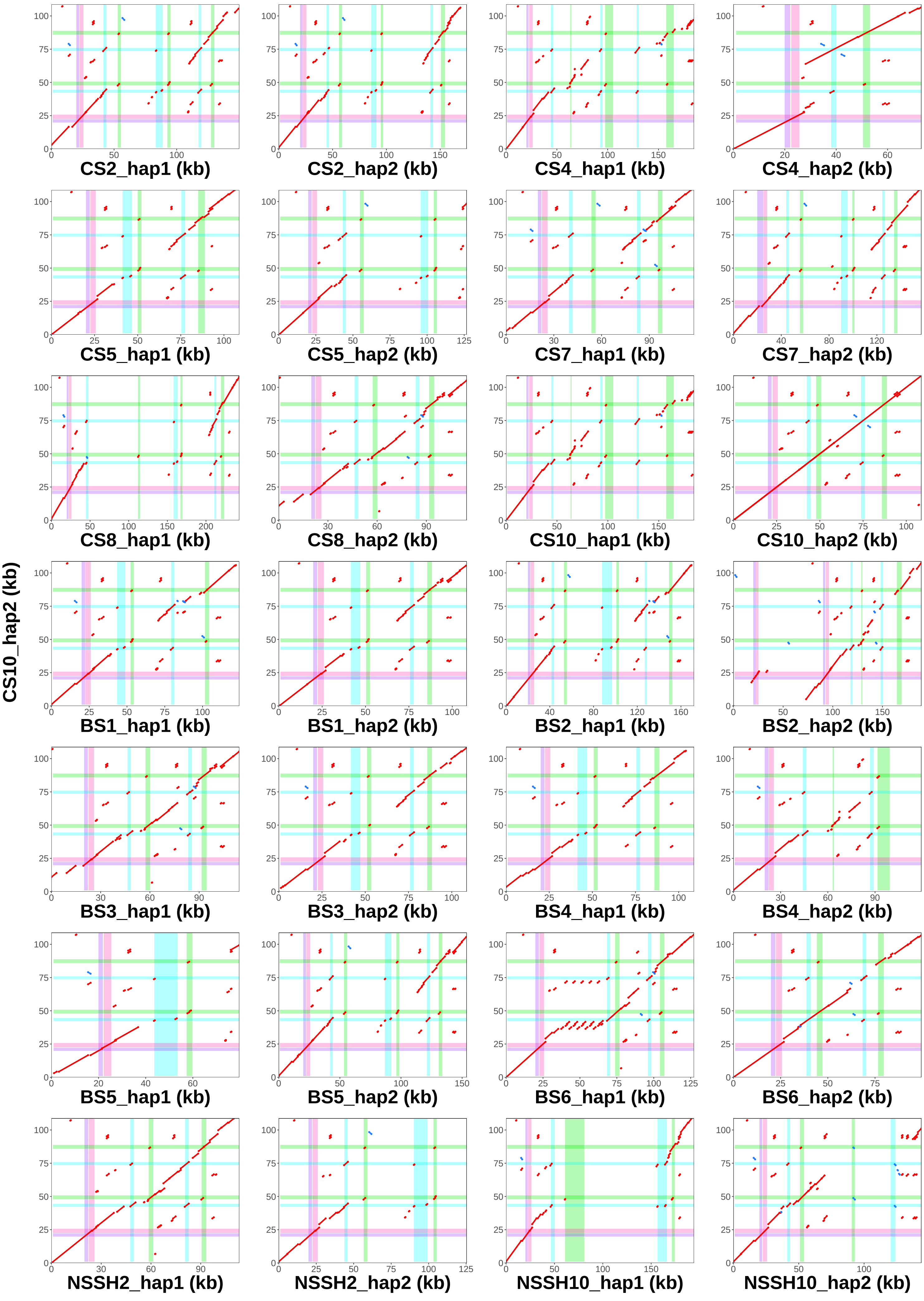

### BS1_hap1 vs. all assemblies − Locus 2

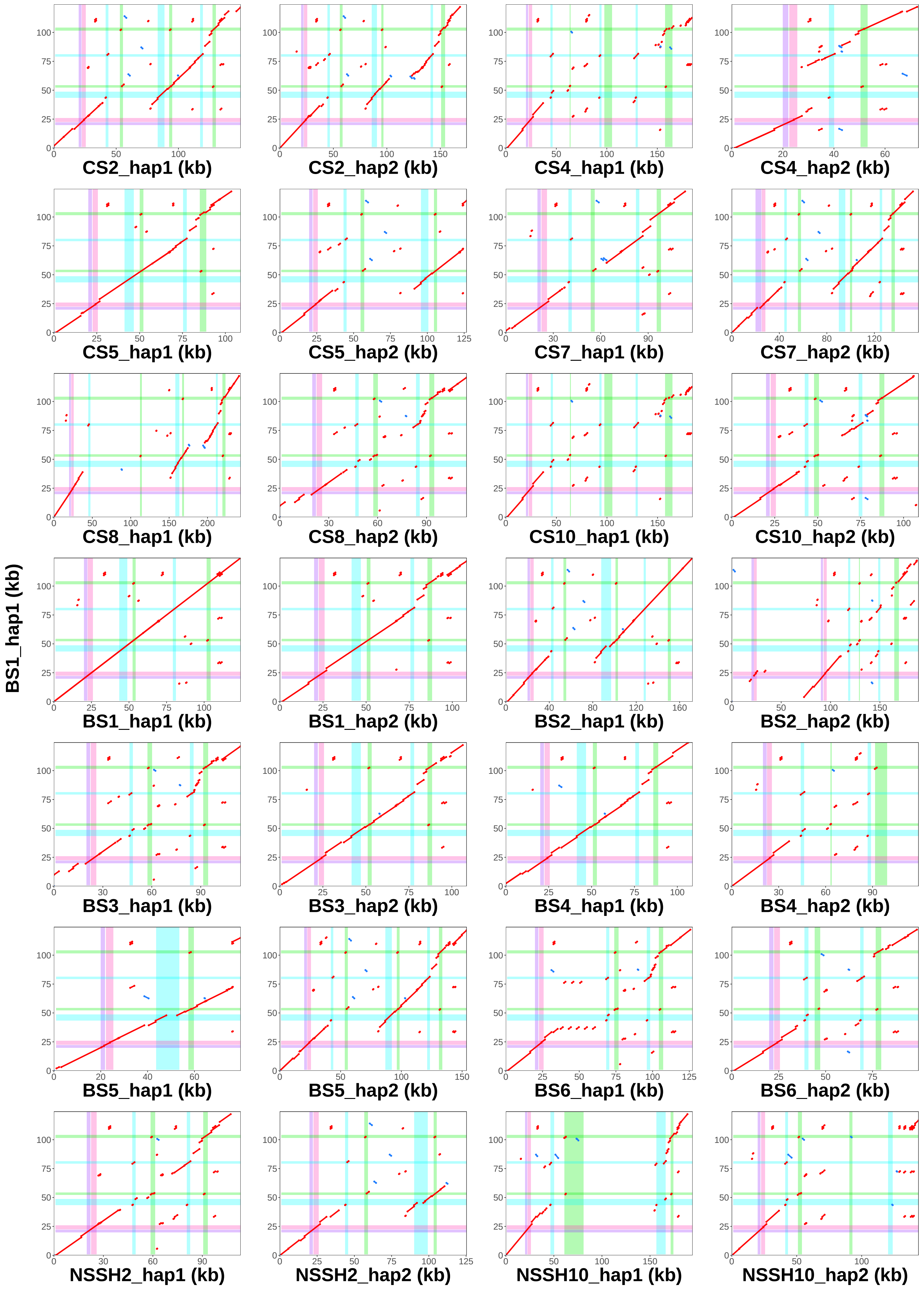

### BS1_hap2 vs. all assemblies − Locus 2

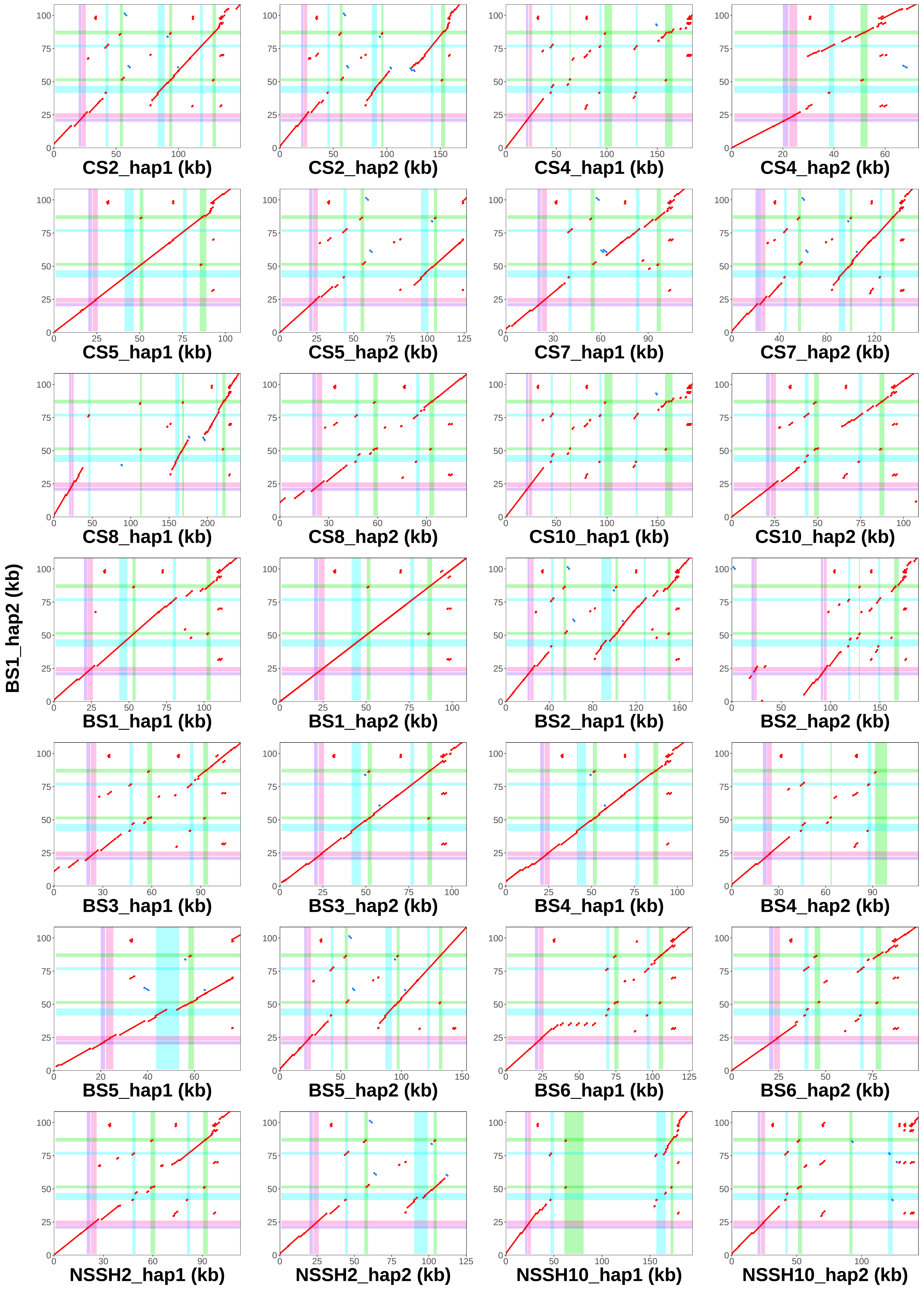

### BS2_hap1 vs. all assemblies − Locus 2

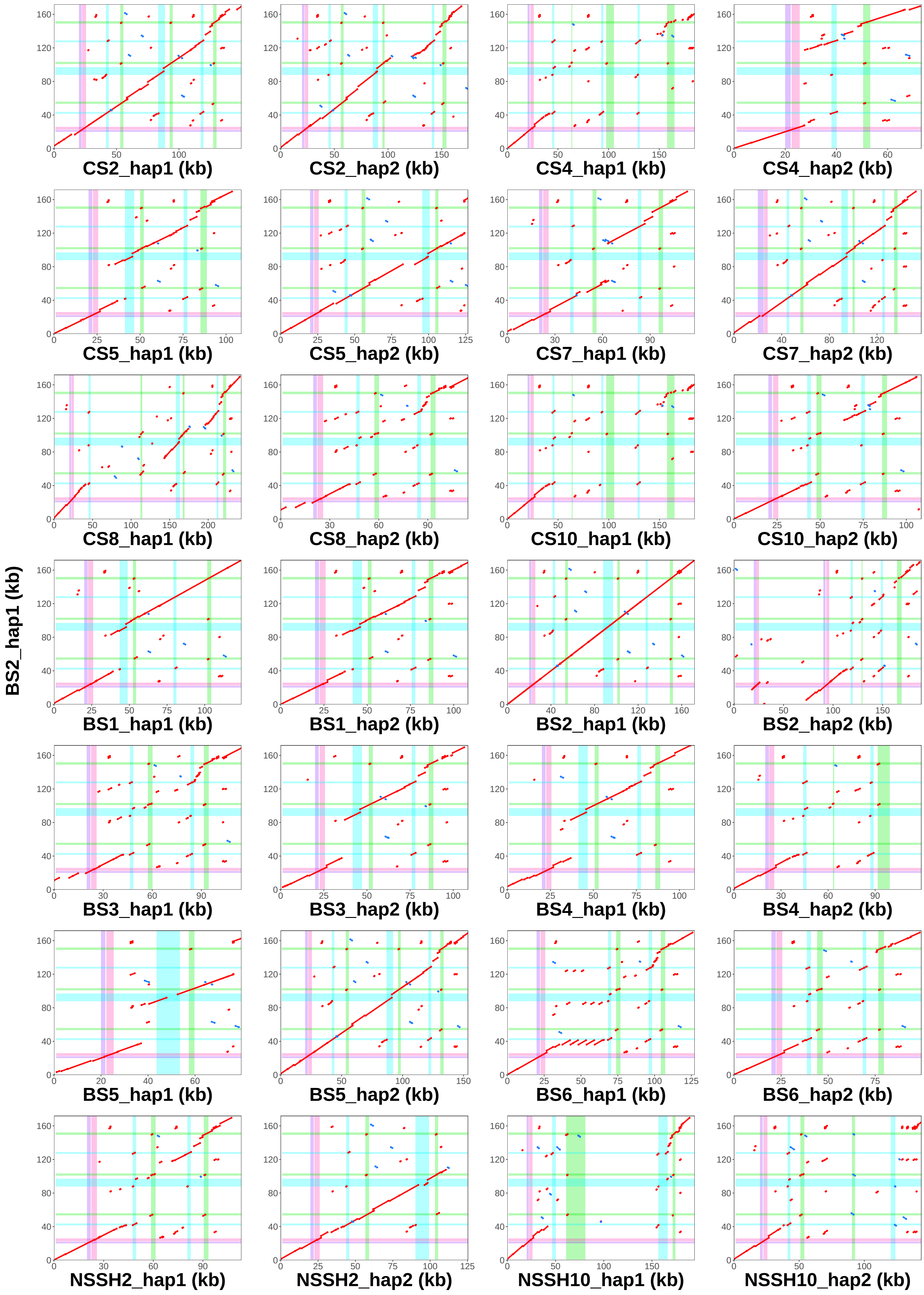

### BS2_hap2 vs. all assemblies − Locus 2

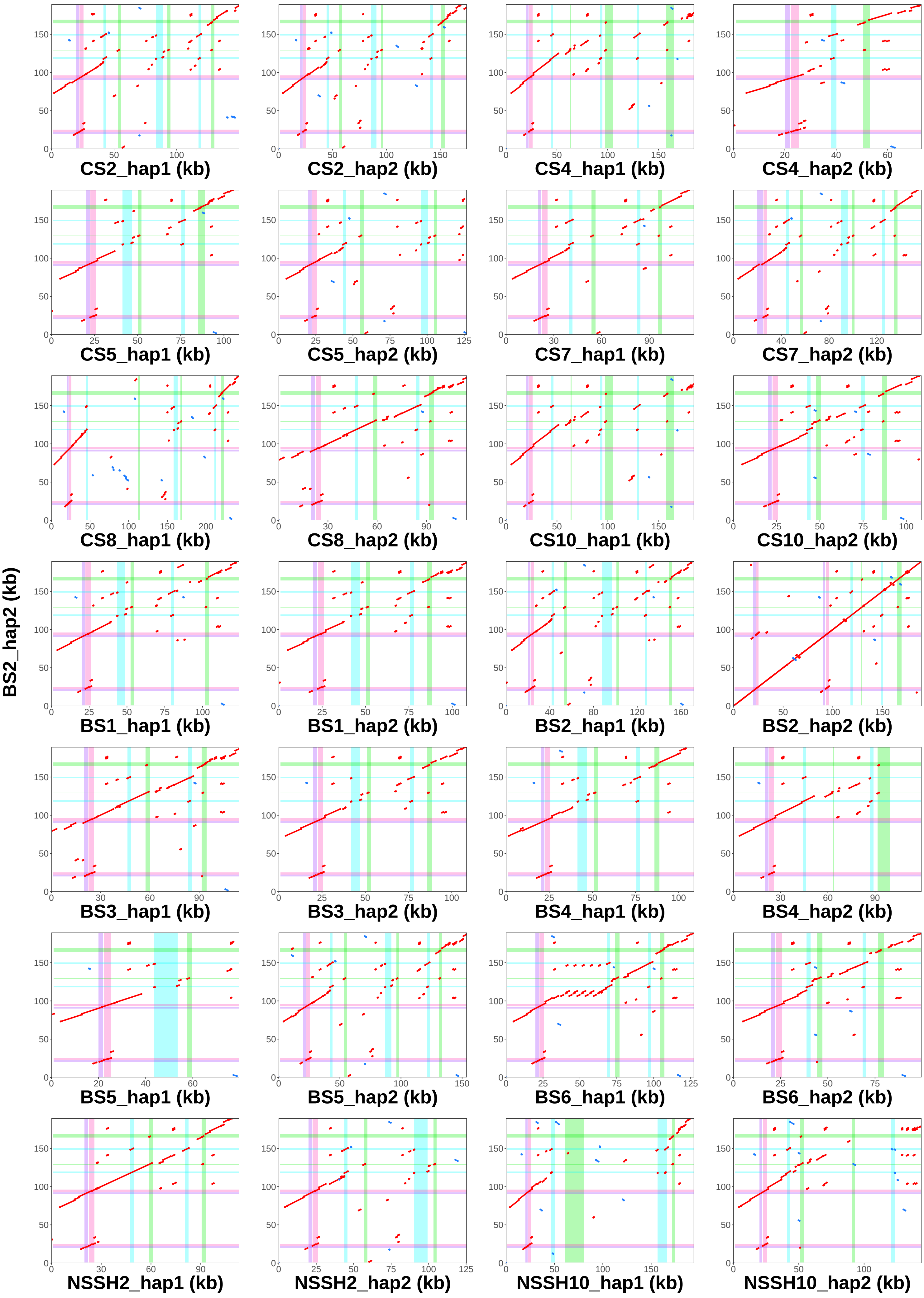

### BS3_hap1 vs. all assemblies − Locus 2

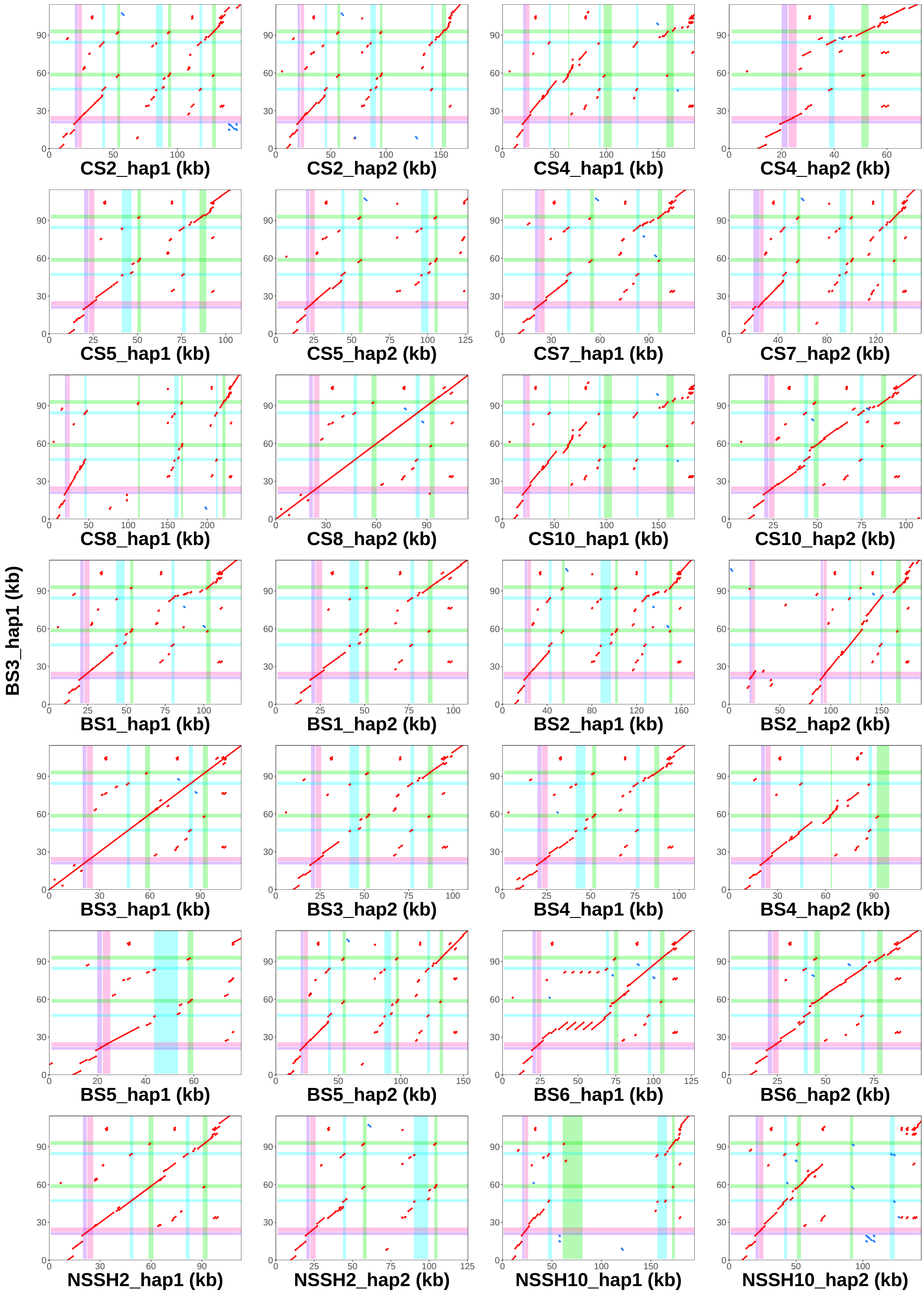

### BS3_hap2 vs. all assemblies − Locus 2

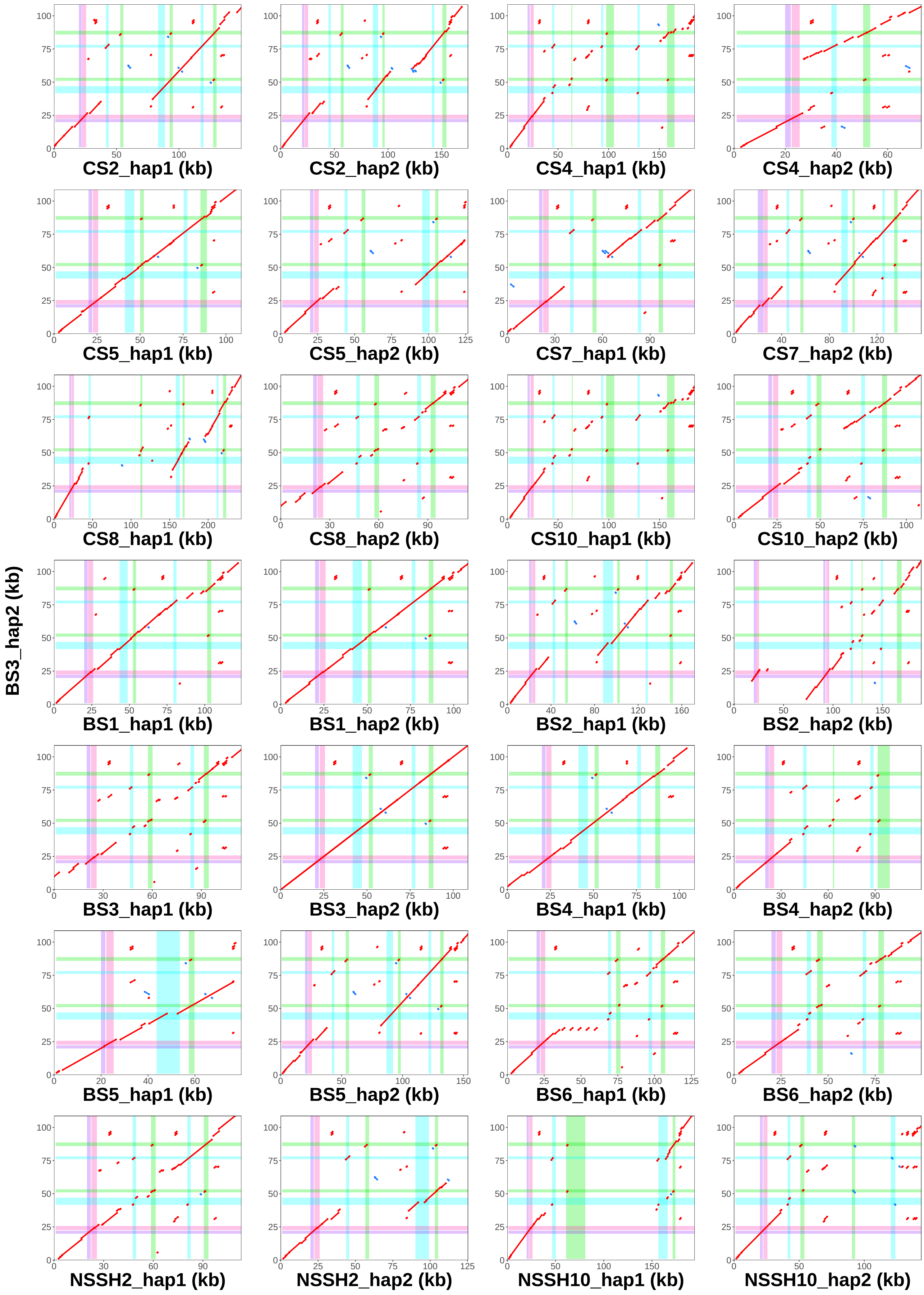

### BS4_hap1 vs. all assemblies − Locus 2

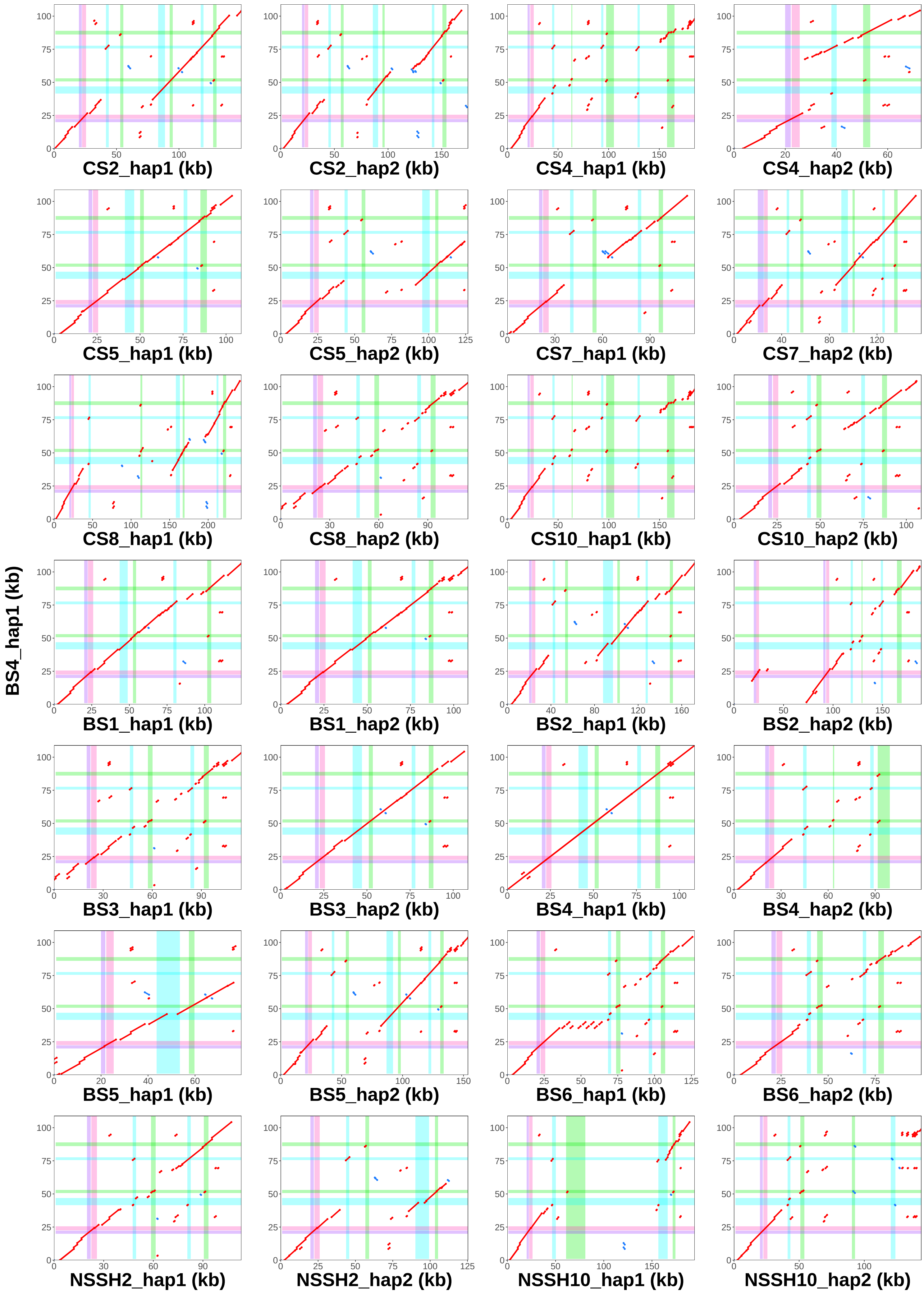

### BS4_hap2 vs. all assemblies − Locus 2

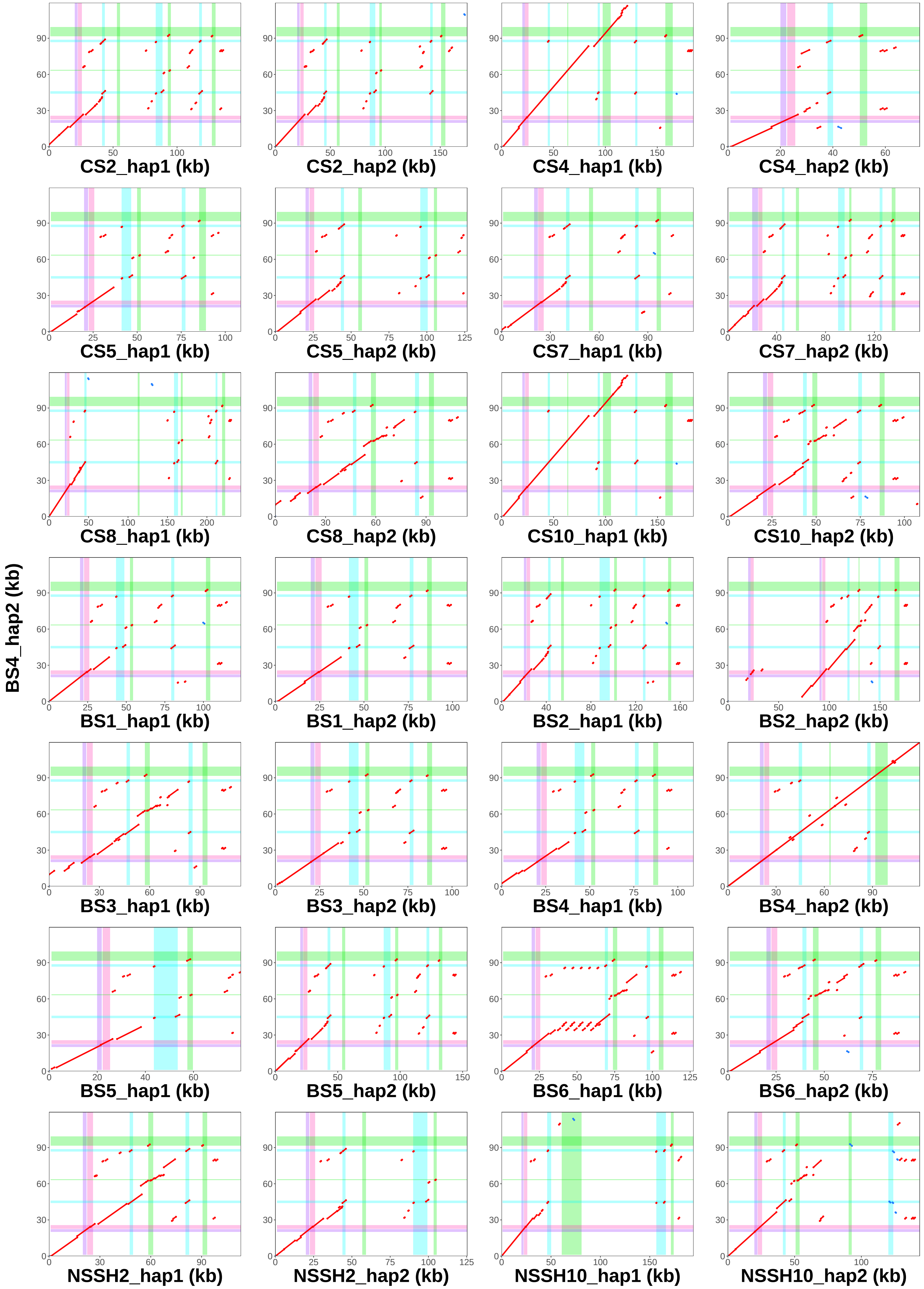

### BS5_hap1 vs. all assemblies − Locus 2

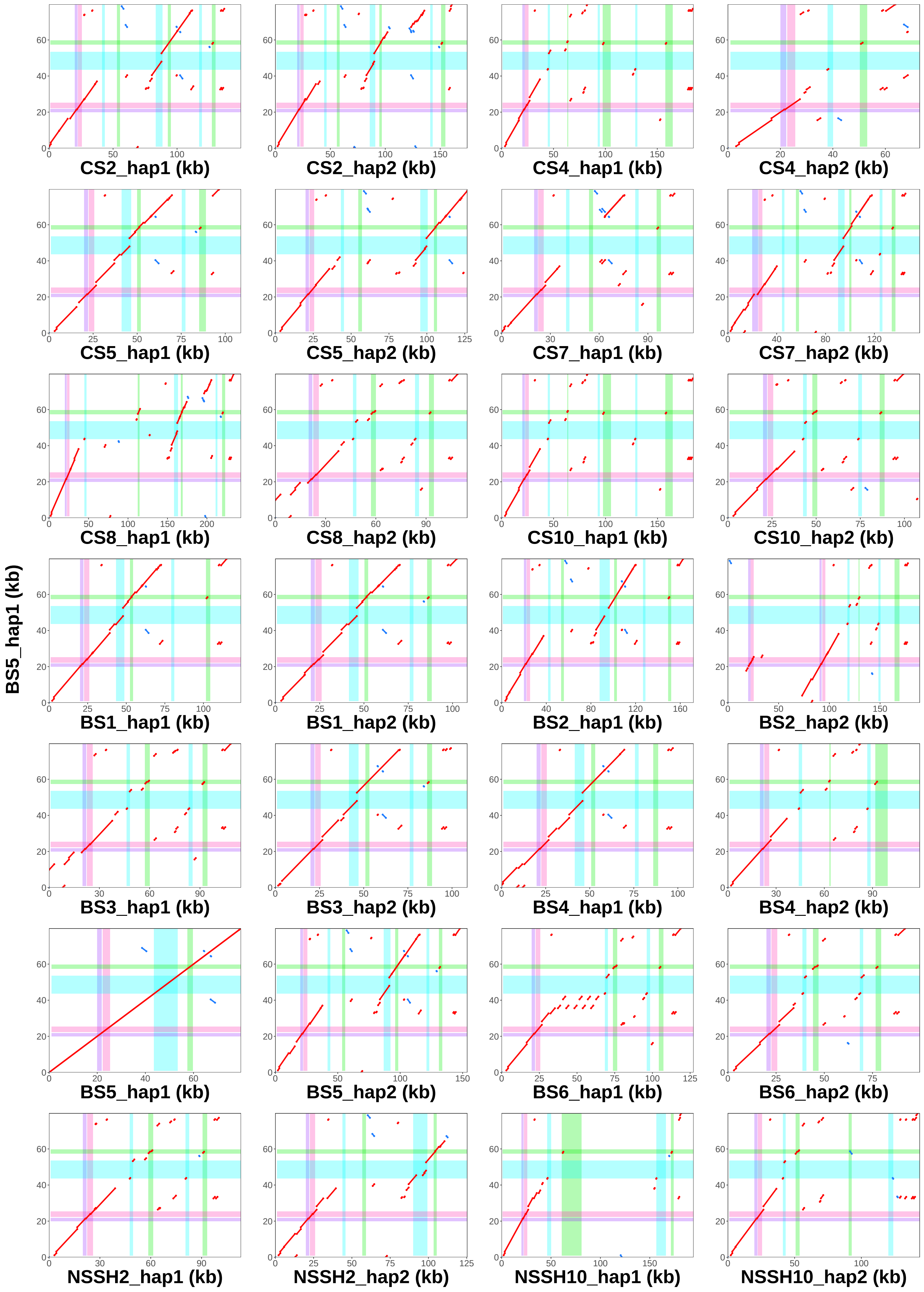

### BS5_hap2 vs. all assemblies − Locus 2

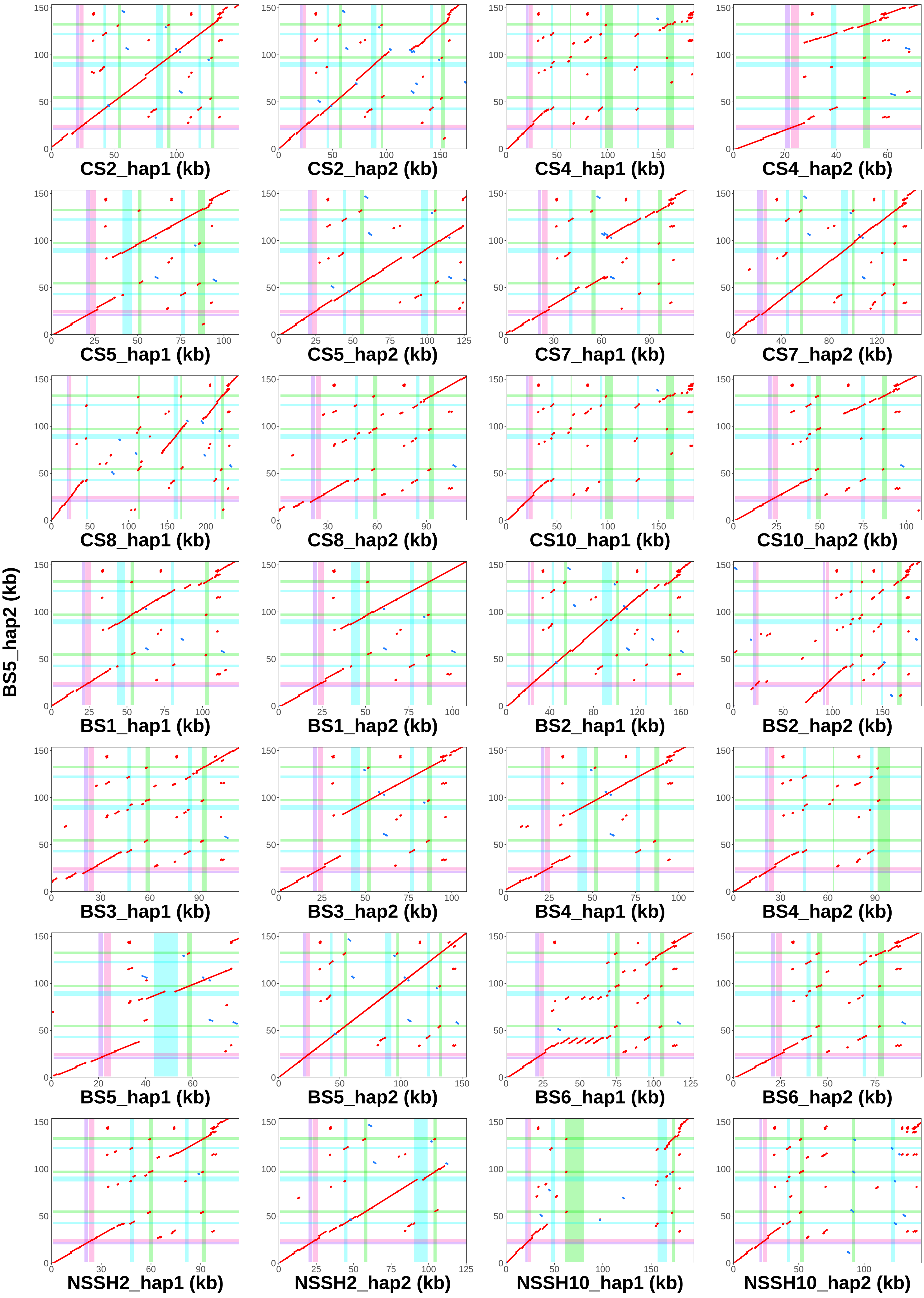

### BS6_hap1 vs. all assemblies − Locus 2

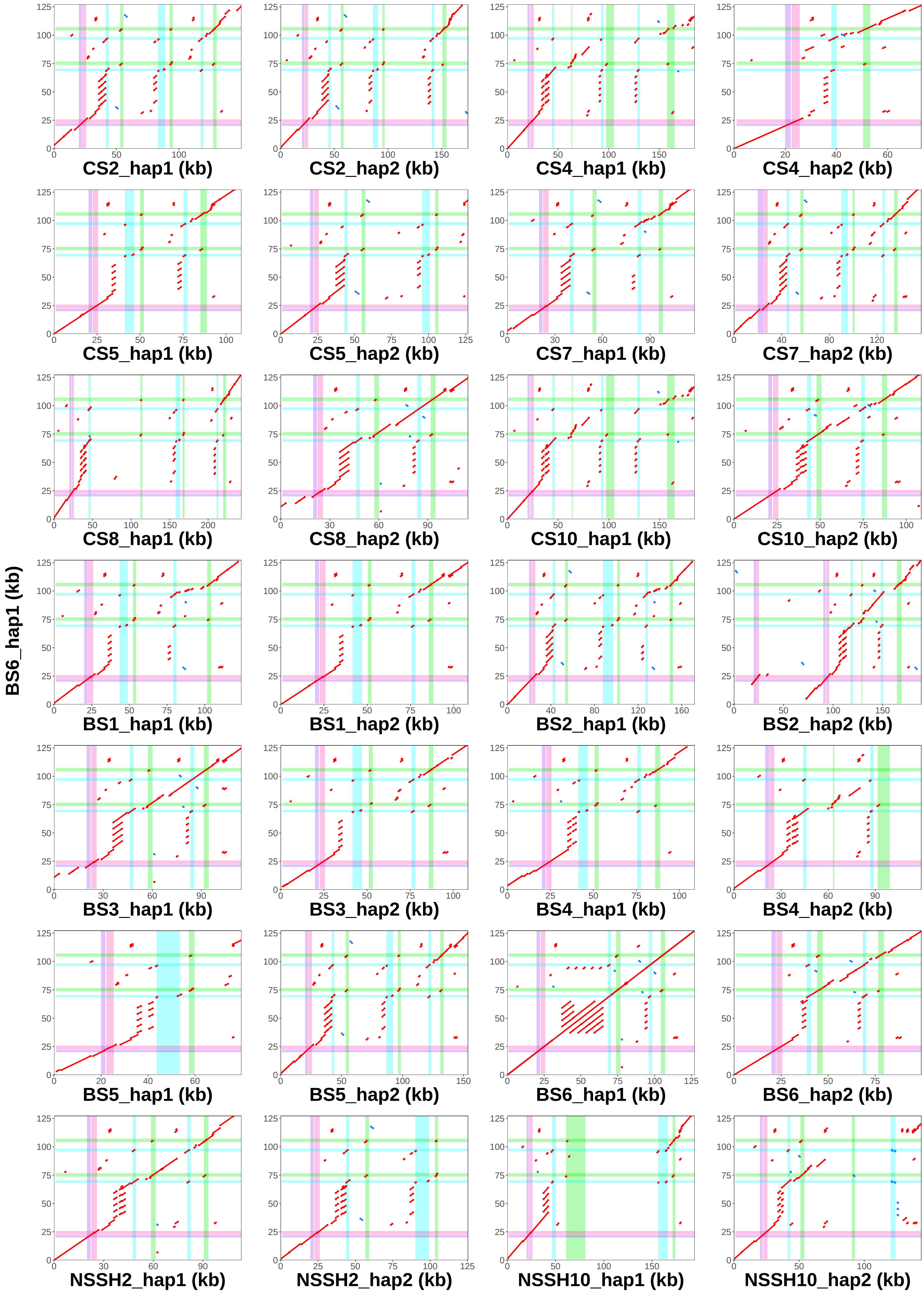

### BS6_hap2 vs. all assemblies − Locus 2

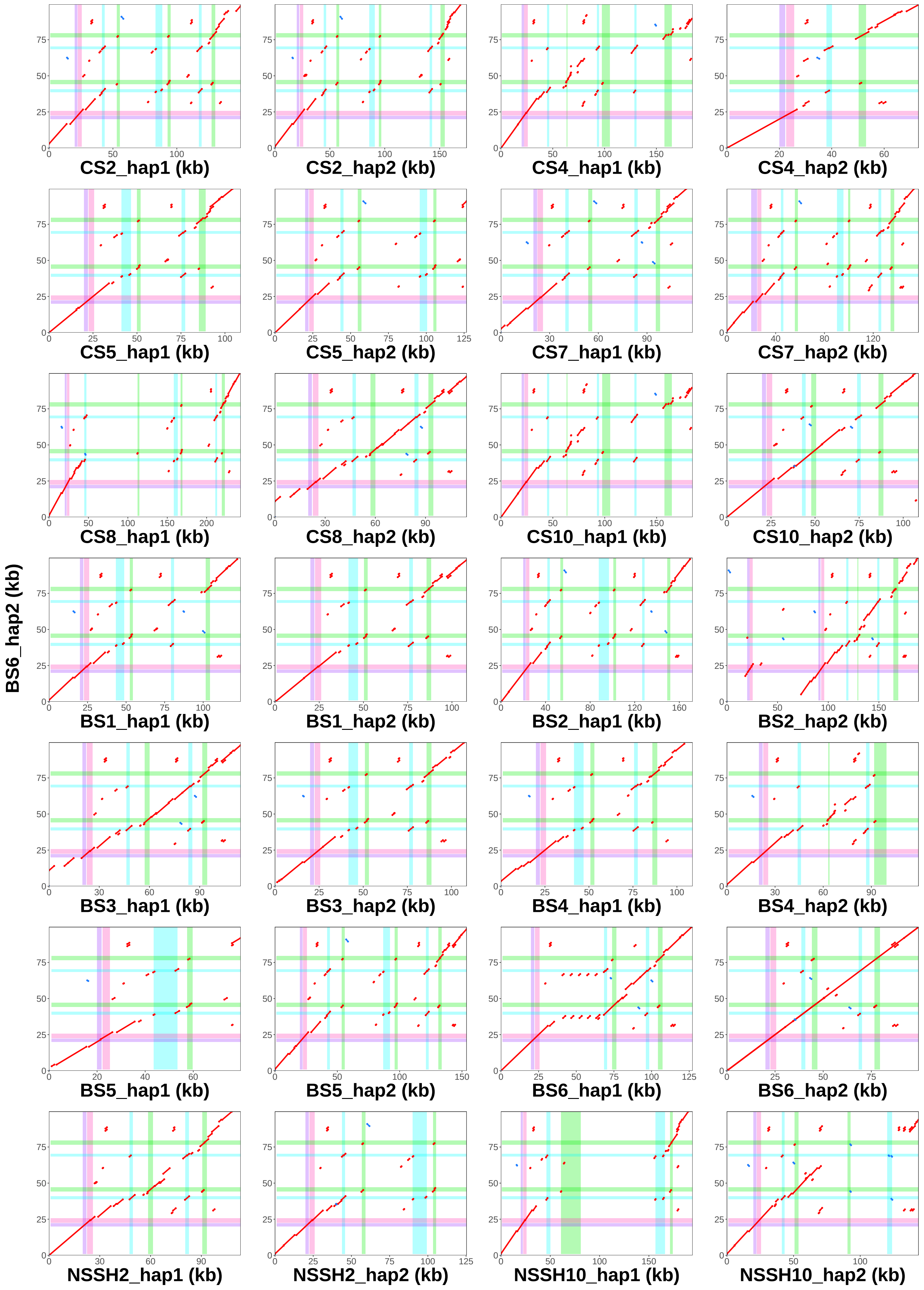

### NSSH2_hap1 vs. all assemblies − Locus 2

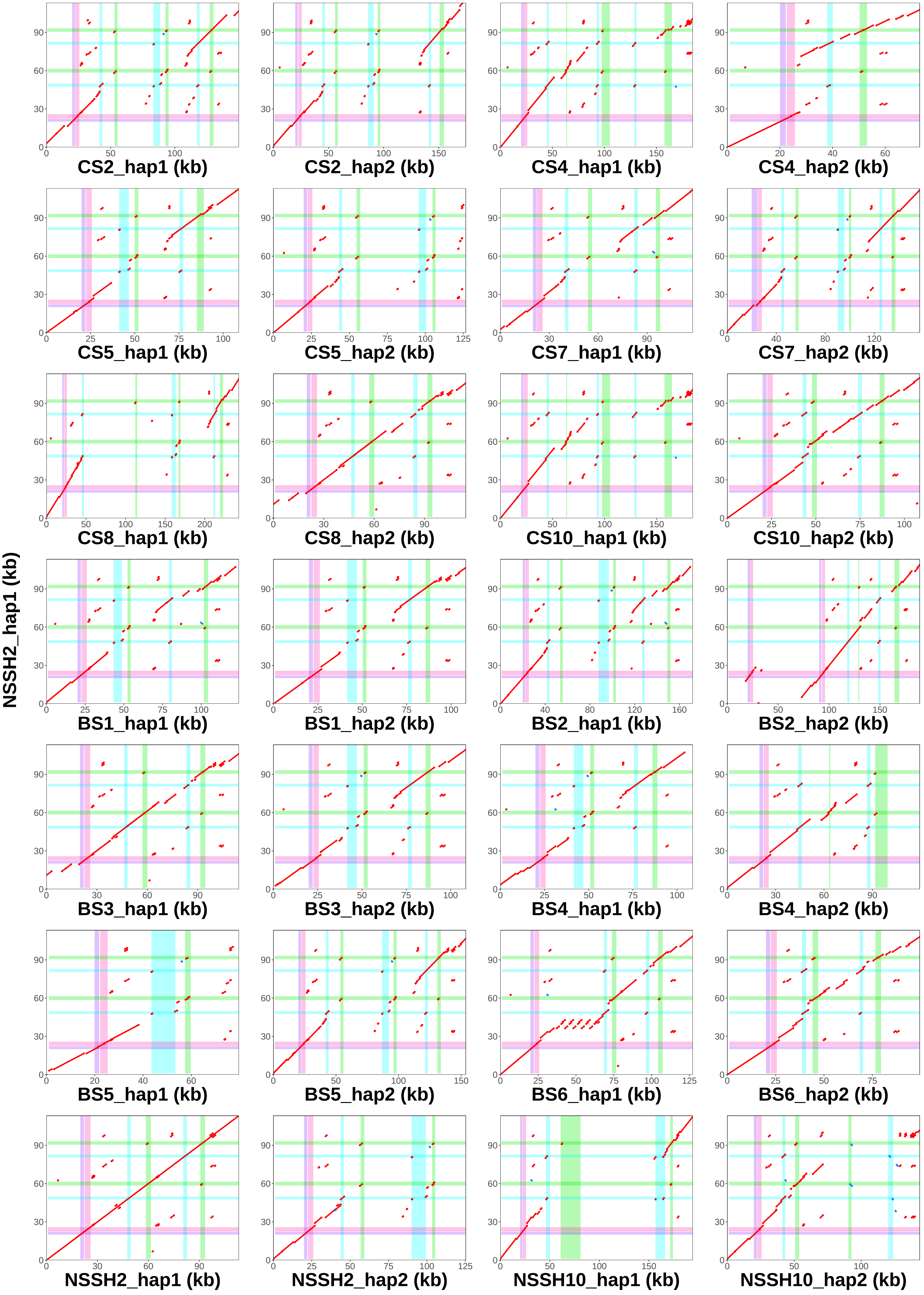

### NSSH2_hap2 vs. all assemblies − Locus 2

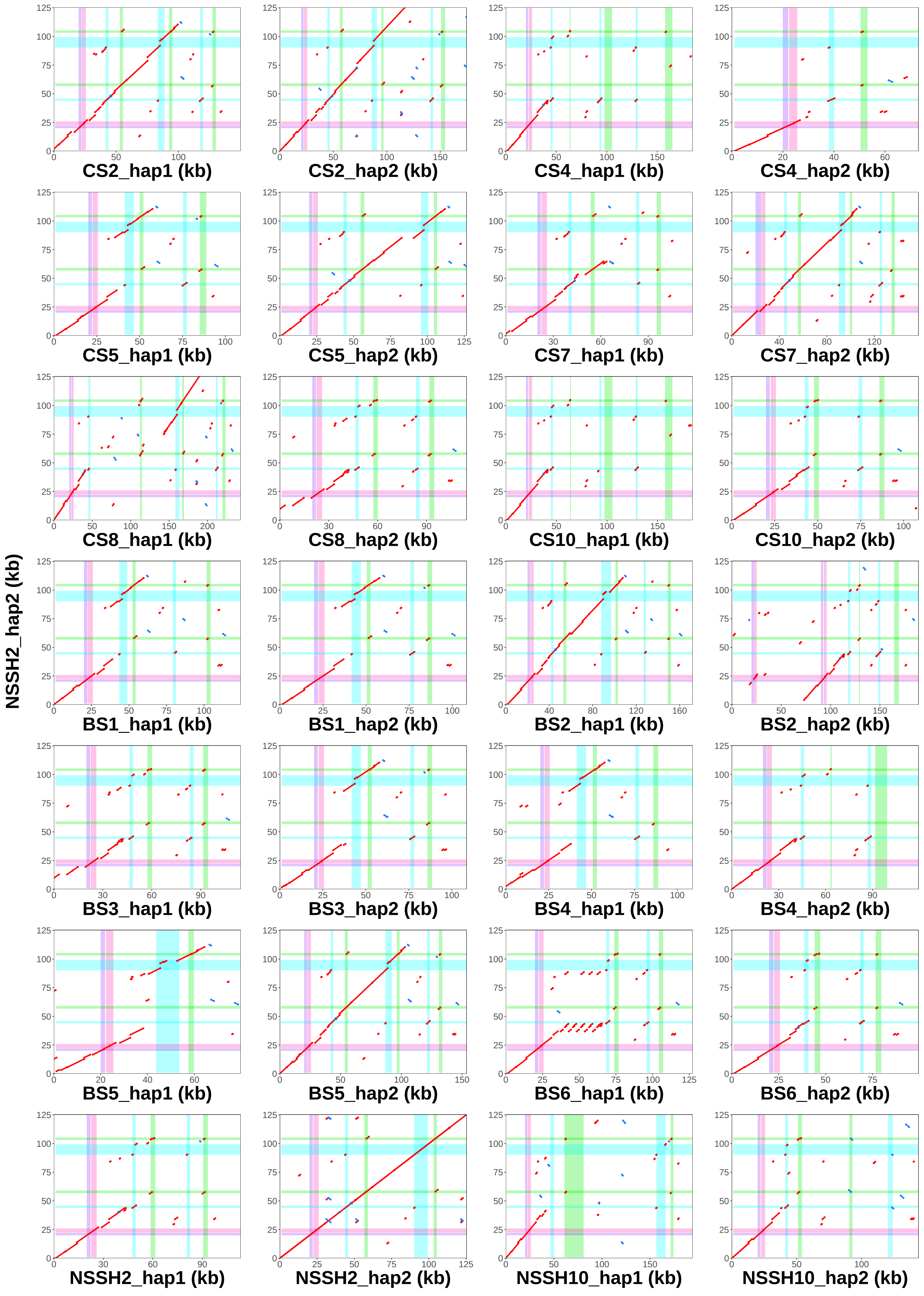

### NSSH10_hap1 vs. all assemblies − Locus 2

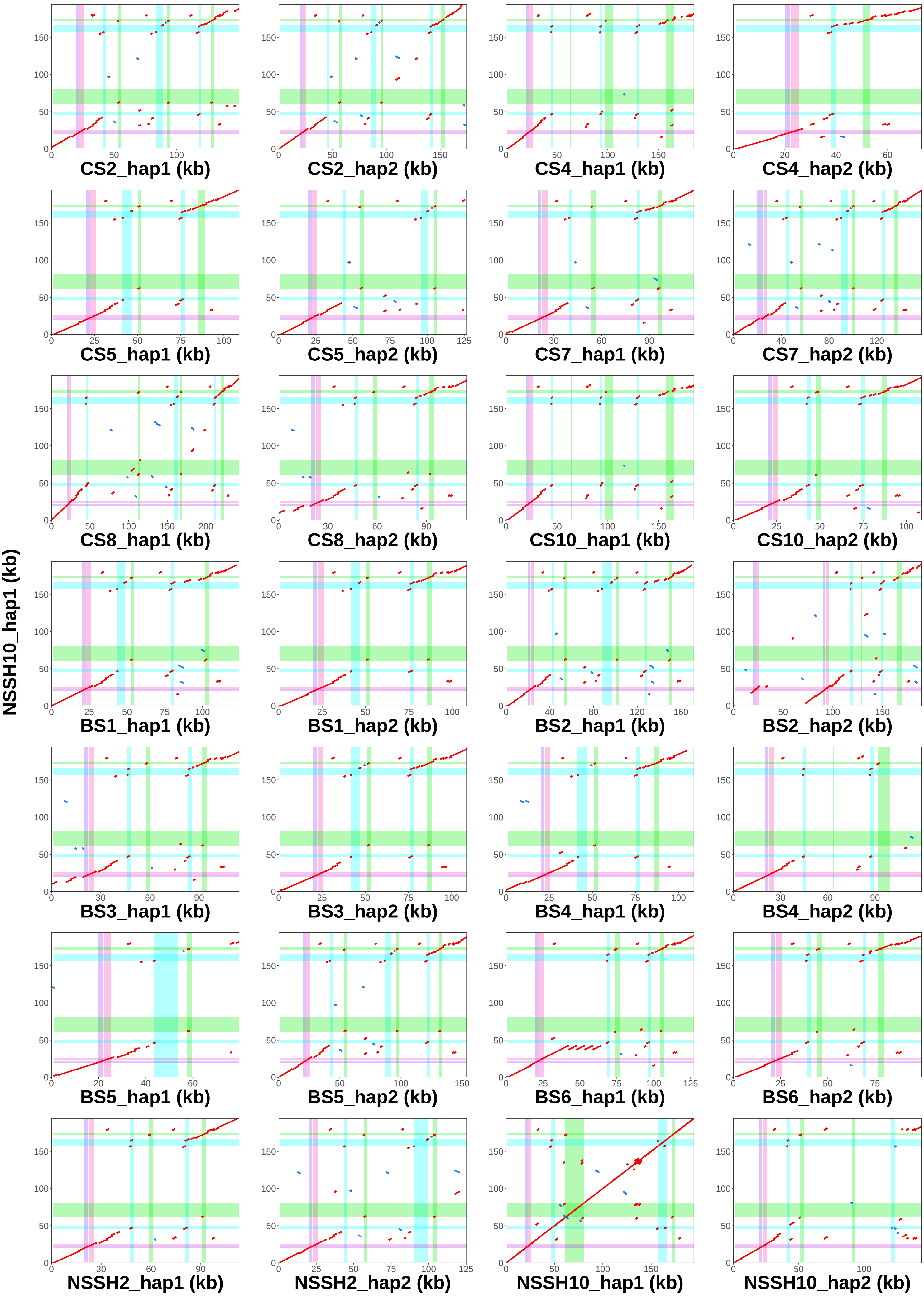

### NSSH10_hap2 vs. all assemblies − Locus 2

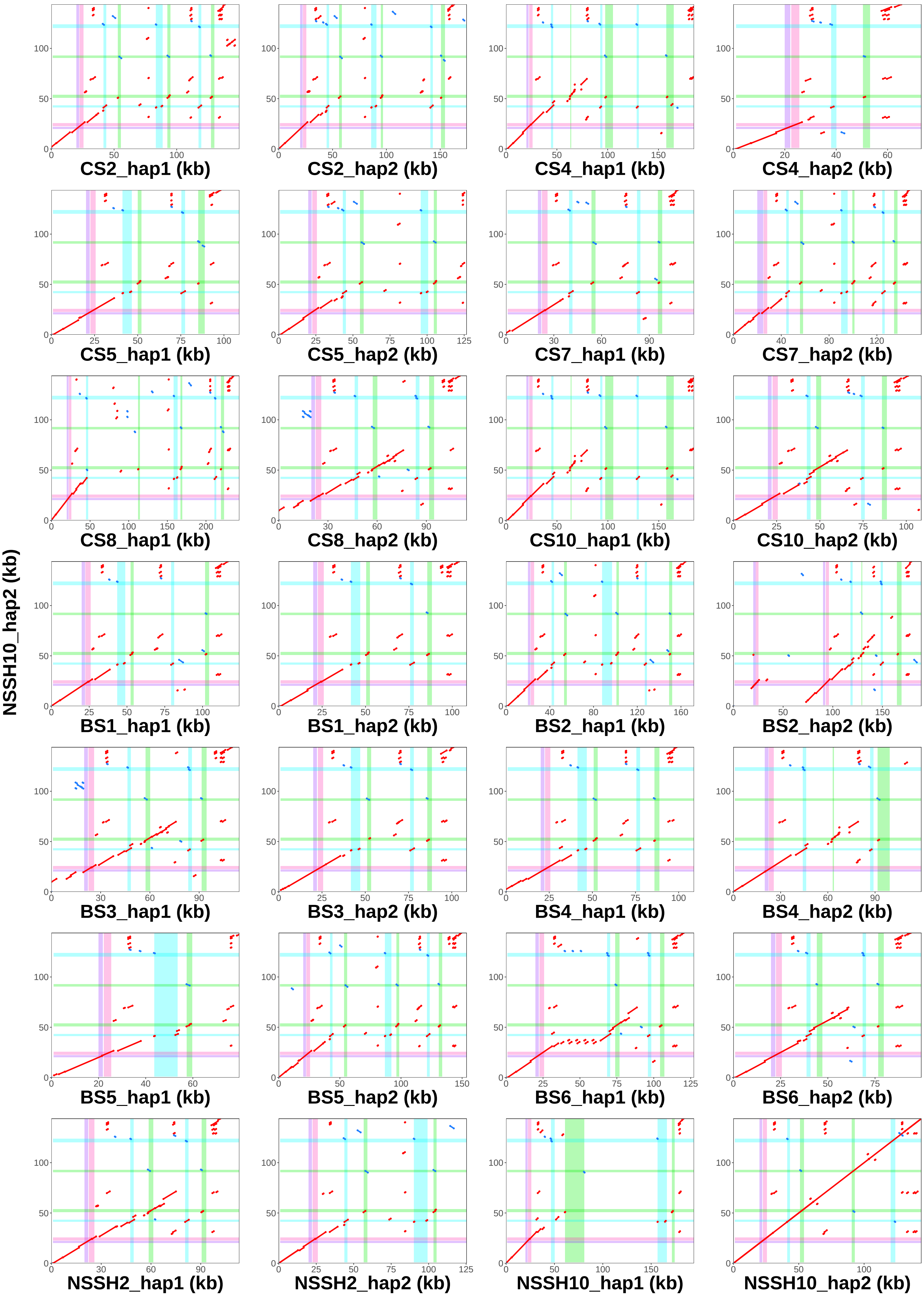
